# A KaiA Conformational Switch Links Day-Night Transitions to Clock Robustness

**DOI:** 10.64898/2026.10.13.758887

**Authors:** Athira C. Chandradasan, Arnaud Taton, Archana G. Chavan, Ning Zhang, Carla A. Omaga, Annia Ariuska Gonzalez Crespo, Luz Mónica Vásquez Pérez, Luis F. Larrondo, Mingxu Fang, Susan S. Golden, Andy LiWang

**Affiliations:** Department of Chemistry & Biochemistry, University of California, Merced, CA, 95343, United States; Department of Molecular Biology, University of California, San Diego, 9500 Gilman Dr., La Jolla, CA, 92093, United States; Department of Microbiology, The Ohio State University, Columbus, OH, 43210, United States; Center for Circadian Biology and Medicine, University of California, San Diego, La Jolla, CA, 92093, United States; Genentech, San Francisco, CA, 94080, United States; Current address: Qingdao Institute of Bioenergy and Process, Chinese Academy of Sciences, Qingdao 266101, China; Facultad de Ciencias Biológicas Pontificia, Universidad Católica de Chile, Avenida Libertador General Bernardo O’Higgins 340 Santiago, Casilla 8331150, Chile; ANID-Millennium Science Initiative Program - Millennium Institute for Integrative Biology (iBio), Santiago 8331150, Chile

## Abstract

Biological clocks require mechanisms that enable transition between temporal phases and buffer against environmental fluctuations. In the cyanobacterial KaiABC-based circadian oscillator, KaiA has been regarded as a daytime activator that is passively sequestered at night. Here we define KaiA as a conformationally regulated activator and an active catalyst of the night-to-dawn transition. We identify a single linker helix, α5, that acts as a structural switch, mediating a domain-driven “tug-of-war” that sets the open–closed equilibrium in KaiA. Structure-guided mutations that shift this balance in either direction abolish stable rhythmicity both *in vitro* and in cells. Also, we show that substoichiometric KaiA dismantles nighttime KaiB–KaiC complexes through interactions with the KaiC C-terminal tails and likely initiates an autocatalytic feedback loop that resets the phosphorylation cycle at dawn. Furthermore, we present two independent kinetic models that offer distinct, deterministic explanations for the clock’s low-temperature amplitude decline and failure at 20 °C.

## Introduction

Circadian clocks enable organisms to anticipate daily environmental change by generating an internal representation of time ^1,2^. In cyanobacteria, this timing system is built from a minimal biochemical oscillator composed of the proteins KaiA, KaiB, and KaiC ^3^, which together drive rhythmic cycles of KaiC phosphorylation that can be reconstituted *in vitro* ^4^ and control global gene expression *in vivo* ^5,6^. KaiC, a hexameric ATPase with tandem CI and CII domains ^7^, undergoes ordered autophosphorylation and autodephosphorylation on two conserved residues in its CII domain ^8,9^, while KaiB and KaiA modulate this cycle by binding distinct KaiC surfaces over the course of the day–night cycle ^10–12^. During the circadian cycle, KaiA, KaiB, and KaiC undergo dynamic conformational changes and assemble into distinct protein complexes at different times of the day ^12^. Metamorphic transitions in KaiB ^13,14^ and changes associated with KaiC phospho-state have roles at the dynamics ^10,11,15^ and structural ^16–18^ levels.

KaiA is a domain-swapped homodimer ^19^ in which each protomer contains an N-terminal *pseudo*-receiver domain ^20^ and a C-terminal four-helix bundle ^21^ that binds the C-terminal tails of KaiC ^11,22,23^. These domains are connected by a flexible linker that includes helix α5, which can engage either the N- or C-terminal domain to produce distinct quaternary conformations ^12^ (**Fig. 1, Supplementary Fig. 1**). Structural and biochemical data indicate that KaiA samples an “open” conformation, in which α5 associates with the N-terminal domain while the C-terminal KaiC-binding surface is exposed, and a “closed” conformation, in which α5 packs against the C-terminal domain and occludes this site.

**Figure 1.**
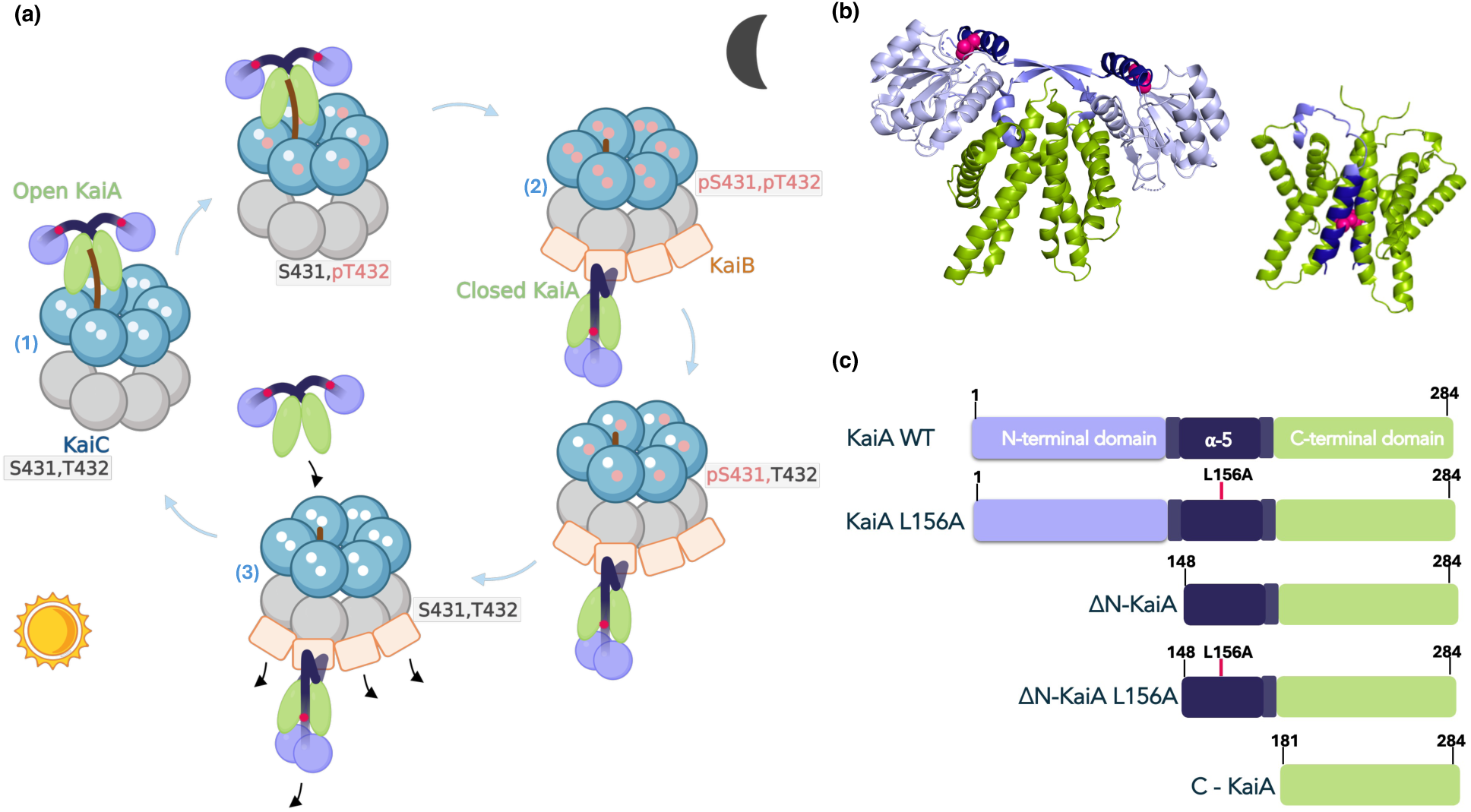
KaiA architecture, conformations, and roles in the KaiABC cycle. **(a)** Schematic of the KaiA–KaiB–KaiC oscillator over 24 h, showing daytime KaiA–KaiC interaction (1), nighttime KaiB–KaiC–KaiA sequestration (2), and morning KaiA-driven KaiB–KaiC disassembly (3). **(b) Left**: Surface/ribbon views of KaiA in the “open” (α5 bound to KaiA N-terminal domain). **Right**: “closed” (α5 packed on KaiA C-terminal domain) conformations. Magenta represents residue L156 in open KaiA (from *S. elongatus*; PDB ID 1R8J) and L155 in closed KaiA (from *Thermosynechococcus* elongatus; PDB ID 5JWR). The closed conformation of KaiA was previously solved using the ΔN KaiA mutant (PDB ID 5JWR). **(c)** Linear cartoons for KaiA WT, KaiA L156A, ΔN KaiA (residues 148-284), ΔN KaiA L156A, and C-KaiA (residues 181-284).

In the canonical model, open KaiA promotes KaiC autophosphorylation during the subjective day, whereas KaiB antagonizes closed KaiA at night by sequestering it within KaiB–KaiC complexes whenever residue S431 is phosphorylated, hereafter referred to as pKaiC. Recent work, however, reframes the day–night transition as a hysteretic switch driven by an underlying bistability in the protein-interaction network, in which KaiA operates within a double-negative feedback loop, actively opposing its own inhibition by catalyzing disassembly of its own suppressor, the KaiB–pKaiC complex ^24^. This model raises two questions that we address here: what tunes KaiA’s open–closed equilibrium and how does that equilibrium shape daytime activation and rhythmicity; and does KaiA actively remodel near-dawn KaiB–KaiC complexes to reset the clock.

As for other circadian systems, cyanobacterial clocks are temperature compensated over a broad physiological range. A temperature-dependent shift in the equilibrium between the two folds of KaiB is likely to play a role in this process ^14^. We therefore sought to address both questions raised above in the context of temperature-dependent regulation. While KaiA was recently found to contribute to temperature compensation by sensing the redox/metabolic state of the cell ^25^, the mechanism by which it feeds this information to the oscillator is not fully resolved.

Here we combine structural analysis, *in vitro* reconstitution of the KaiABC oscillator (IVO), *in vivo* reporter assays, KaiB-KaiC equilibrium binding assays, and kinetic modeling to define KaiA’s roles across the day–night cycle and how they are modulated by temperature. First, we show that helix α5 mediates a domain-driven tug-of-war that sets the balance between open ⇌ closed KaiA, and that this balance governs daytime activation of KaiC and the ability to sustain rhythms. Second, we demonstrate that dephosphorylated KaiC (dpKaiC) forms stable complexes with KaiB at high temperature and that KaiA catalyzes their disassembly through interactions with the KaiC C-terminal tails. Third, we quantify how the KaiB–dpKaiC interaction weakens with cooling and show that this change promotes premature nighttime-complex breakdown at low temperatures. Finally, we show that two independent models, a kinetic model built from these measured processes (the Autocatalytic Dawn model) and a recently published bistability model of the KaiA–KaiB antagonism ^24^, both reproduce the amplitude decline and the 20 °C collapse as deterministic oscillator death; the bistability model additionally reproduces the oscillator’s temperature-compensated period.

## Results

### An α5-helix–mediated tug-of-war between KaiA domains controls day–night conformations

To understand how KaiA’s structure encodes its distinct daytime and nighttime activities, we examined how the N-terminal (N-KaiA) and C-terminal (C-KaiA) domains, linked by helix α5 (residues 148–164), regulate its open ⇌ closed conformational equilibrium. KaiA adopts an open conformation in which α5 packs against N-KaiA and leaves the C-KaiA dimer surface available to bind KaiC C-terminal tails ^11,22^, and a closed conformation in which α5 instead packs against C-KaiA and occludes its KaiC-binding site ^12^. (**Fig. 1**) We reasoned that mutually exclusive interactions of α5 with each domain would create a domain-driven tug-of-war that tunes the relative populations of these conformations and thereby regulates KaiA’s function across the circadian cycle.

### Removing the N-terminal domain biases KaiA toward a closed conformation

We generated a truncation lacking N-KaiA (ΔN-KaiA), while retaining α5 and C-KaiA, to test whether N-KaiA is required to stabilize the open state. Size-exclusion chromatography coupled to multi-angle light scattering (SEC-MALS) showed that ΔN-KaiA forms a stable homodimer of the expected mass (**Fig. 2a–b, Supplementary Fig. 2**). However, ΔN-KaiA minimally stimulated KaiC autophosphorylation (**Fig. 2c, Supplementary Fig. 3**) and failed to support circadian rhythms *in vitro*, in contrast to KaiA WT (wild-type KaiA) (**Fig. 3a, Supplementary Fig. 4**). KaiA WT delays the onset of KaiB–KaiC binding (**Supplementary Fig. 5-6**), consistent with KaiA’s antagonistic effect on KaiB–KaiC interactions, but no such delay was observed with ΔN-KaiA (**Fig. 3a**; compare amber and purple peak times, **Supplementary Fig. 6**), indicating that ΔN-KaiA lacks this antagonistic activity. A reporter strain expressing firefly luciferase from the *kaiBC* promoter (P*kaiBC-luc*) together with ΔN-KaiA in place of KaiA WT failed to sustain rhythms *in vivo*: reporter oscillations were strongly damped under 12-h light:12-h dark (LD) cycles and were undetectable after ∼24 h under constant light (LL), resembling a KaiA-null strain (**Fig. 3b**). Thus, although ΔN-KaiA is dimeric like KaiA WT, it is functionally compromised, suggesting that in the absence of the N-KaiA domain, α5 engages the C-KaiA domain to favor a closed, KaiC-inaccessible conformation, as observed in the closed KaiA structure (PDB 5JWR, chains E and F; ^12^).

**Figure 2.**
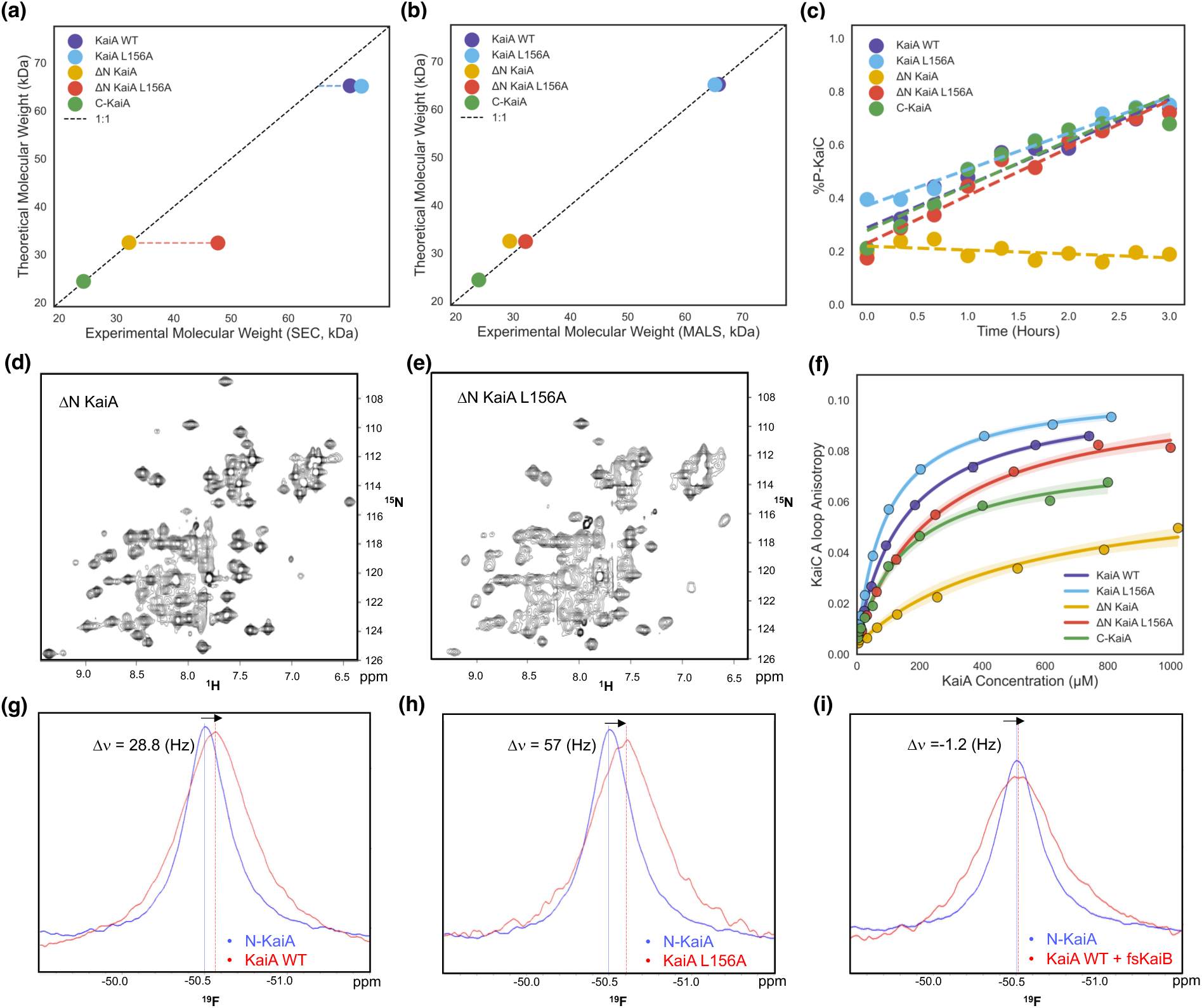
The α5-helix–mediated tug-of-war between KaiA domains tunes its open–closed equilibrium. **(a)** Comparison of theoretical and size-exclusion chromatography-determined molecular weights for KaiA mutants (Supplementary Table 11). **(b)** Comparison of theoretical and multi-angle light scattering (MALS) determined molecular weights for KaiA mutants (Supplementary Table 11). **(c)** KaiC initial phosphorylation kinetic profiles in the presence of KaiA mutants. (Supplementary Table 6). **(d)** 15N, 1H heteronuclear single-quantum correlation (HSQC) spectra of uniformly 15N, 2H-enriched ΔN KaiA and **(e)** ΔN KaiA L156A. **(f)** FA profiles of KaiC A loop peptides as a function of KaiA mutants. (Supplementary Table 7). **(g)** ^19^F–Trp NMR of N-KaiA in comparison to KaiA WT, **(h)** KaiA L156A, **(i)** KaiA WT with fold-switched KaiB (Supplementary Table 8).

**Figure 3.**
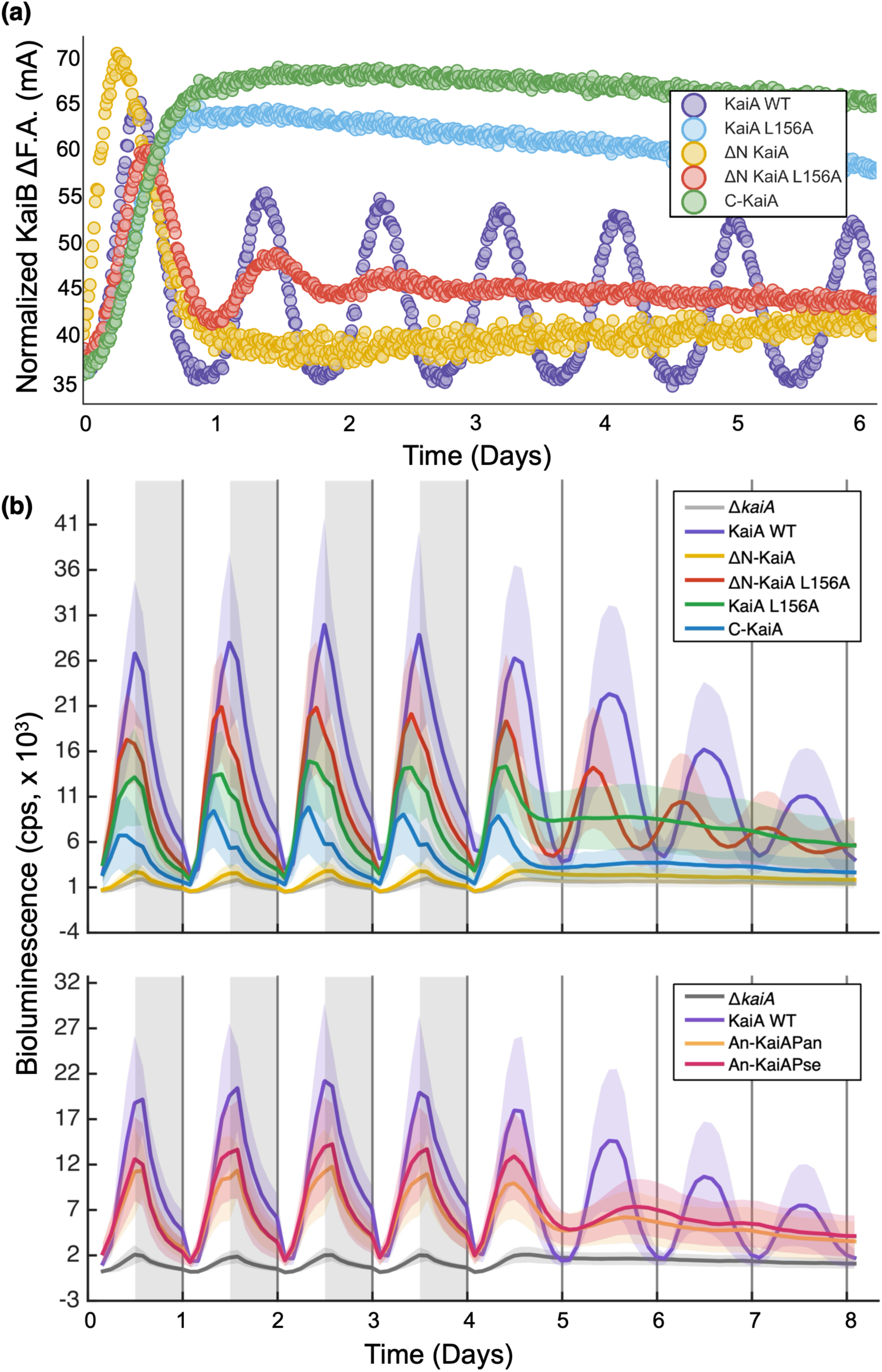
*In vitro* and *in vivo* assays. **(a)** *In vitro* oscillator anisotropy assays with different KaiA variants at 30°C with 6IAF-labeled 50 nM KaiB as the probe. Each reaction includes KaiC WT (3.45 μM), KaiB WT (3.45 μM), and KaiA mutant (1.2 μM). KaiA WT has robust rhythms with a period of 22.38 ± 0.32 h (n=8), and ΔN-KaiA L156A rescues in vitro rhythms with a longer period (23.29 ± 0.54 h, n=7). ΔN-KaiA, KaiA L156A, and C-KaiA anisotropies are arrhythmic. **(b)** P*_kaiBC_*-*luc* bioluminescence over an 8-day time course, including 4 days under 12 h light:12 h dark, followed by 4 days under continuous light conditions at 30 °C, in *S. elongatus* Δ*kaiA* strains expressing ectopically *S. elongatus* KaiA WT (*T*: 24.20 ± 0.67 *ɸ*: 11.98 ± 0.59 h), ΔN-KaiA, ΔN-KaiA L156A (*T*: 21.82 ± 0.72 *ɸ*: 10.20 ± 0.73 h), KaiA L156A, and C-KaiA (upper plot), or *S. elongatus* KaiA WT (*T*: 23.74 ± 0.43 *ɸ*: 12.31 ± 0.43 h), *Anabaena* PCC 7120 KaiA, either An-KaiAPan expressed from its native promoter or An-KaiAPse expressed from *the S. elongatus kaiA* promoter (bottom plot). Δ*kaiA* strains carrying the corresponding antibiotic resistance gene alone were used as negative controls, respectively. Period (*T*) and acrophase (*ɸ*) were estimated by fitting a chirped damped cosine model to the detrended bioluminescence time series; values are shown as mean ± SD (n>9, incl. 3-4 individual clones).

### Weakening α5–C-KaiA packing by an L156A substitution re-opens ΔN-KaiA

Using the homologous protein from *Thermosynechococcus elongatus*, we previously found that substituting its α5 residue L155 (analogous to *S. elongatus* L156) with alanine (L155A) weakened KaiA’s ability to be sequestered by the KaiB–pKaiC complex (Tseng et al., 2017) (**Supplementary Fig. 1**). To probe the role of α5 contacts in stabilizing the closed state of *S. elongatus* KaiA, we made the analogous L156A substitution. In the ΔN background, ΔN-KaiA L156A eluted with a larger apparent hydrodynamic radius by SEC-MALS than ΔN-KaiA did, consistent with an ensemble shifted toward more extended, open-like conformations (**Fig. 2a, Supplementary Fig. 2**). NMR spectra of ^15^N,^2^H-enriched proteins showed that many resonances present in ΔN-KaiA are broadened in ΔN-KaiA L156A (**Fig. 2d-e**), consistent with ΔN-KaiA populating a stable closed-like state and ΔN-KaiA L156A adopting a more flexible conformation with greater µs–ms internal motions.

Fluorescence anisotropy (FA) using a labeled KaiC A-loop peptide ^11^ showed that ΔN-KaiA L156A binds the A-loop with substantially higher affinity than ΔN-KaiA does, and with a temperature dependence resembling KaiA WT, reinforcing the conclusion that weakening α5– C-KaiA packing shifts the equilibrium toward an open, KaiC-binding competent state (**Fig. 2f, Supplementary Fig. 7**). Functionally, ΔN-KaiA L156A restored KaiC autophosphorylation *in vitro* to near wild-type levels (**Fig. 2c, Supplementary Fig. 3**) and supported clear, though damped, rhythms for two cycles in reconstitution assays (**Fig. 3a, Supplementary Fig. 4**). *In vivo*, reporter strains expressing ΔN-KaiA L156A in place of KaiA WT sustained high-amplitude oscillations under LD and LL, with gradual damping over several cycles and a slightly shorter period and phase advance relative to KaiA WT (**Fig. 3b, Supplementary Fig. 8**), consistent with partial rescue. The N-terminal domain is therefore not essential for entrainment, nor is it essential for resetting **(Supplementary Fig. 9)**. Together, these data show that removing N-KaiA biases KaiA toward a closed state, and that weakening α5–C-KaiA packing by L156A re-opens ΔN-KaiA and partially restores its ability to support rhythms.

Notably, adding ΔN-KaiA to ΔN-KaiA L156A produced more robust *in vitro* rhythms with shorter periods (**Supplementary Fig. 10**), which is like the effect of CikA on suboptimal levels of KaiA WT *in vitro* ^26,27^. Even so, these rhythms desynchronized over days, underscoring the role of the N-terminal domain in maintaining robust synchronization (**Supplementary Fig. 10**).

### L156A in full-length KaiA produces a constitutively open, dysregulated activator

We next asked how destabilizing the α5–C-KaiA interface affects the full-length protein. Full-length KaiA L156A behaved like KaiA WT by SEC-MALS (**Fig. 2a–b, Supplementary Fig. 2**) but bound the KaiC peptide more tightly than all the other variants tested (**Fig. 2f, Supplementary Fig. 7**), indicating that it populates the open conformation more than KaiA WT does. In phosphorylation assays, KaiA L156A stimulated KaiC autophosphorylation to WT or slightly higher levels (**Fig. 2c, Supplementary Fig. 3**), yet *in vitro* reconstitutions driven by KaiA L156A failed to oscillate in FA, indicating persistently high KaiB–KaiC binding. (**Fig. 3a, Supplementary Fig. 4**). To confirm constitutive stabilization of the open conformation in KaiA L156A, we ran SEC-MALS with KaiC EE 505 in the presence of fold-switched KaiB. This KaiC variant carries the double phosphomimetic mutations S431E and T432E to promote KaiB binding and lacks the C-terminal tail to eliminate direct KaiA-KaiC interactions. KaiA L156A could not form a KaiABC complex in contrast with KaiA WT, confirming that a constitutively open KaiA cannot be sequestered. (**Supplementary Fig. 2**).

Because KaiA WT (∼64 kDa as a dimer) is challenging for conventional NMR, we used ^19^F tryptophan NMR to monitor residue W10 in the N-terminal domain, the only tryptophan residue in the protein (**Fig. 2g–i**). Comparing 6-fluoro- and 7-fluoro-Trp labeling, 7-fluoro-Trp yielded narrower linewidths (**Supplementary Fig. 11)** and was used throughout. The ^19^F NMR signal of the isolated N-terminal domain represents the fully closed conformation - there is no α5 helix for N-KaiA to bind, as happens in the open conformation. In contrast, the ^19^F frequency of KaiA L156A, which destabilizes the closed state, is assumed to represent the fully open conformation. The KaiA L156A signal was 57 Hz upfield of the N-KaiA signal at 14.1 T and 30 °C, whereas KaiA WT was only 29 Hz upfield, indicating an open ⇌ closed equilibrium constant K_eq_ ≈ 1 (**Fig. 2h, Supplementary Fig. 11**). Adding a KaiB variant locked in the fold-switched conformation that sequesters KaiA on the KaiC CI ring at night (fsKaiB) shifted the KaiA WT ^19^F resonance downfield to coincide with that of N-KaiA (**Fig. 2i**), indicating that fsKaiB drives the equilibrium far toward the closed state. *In vivo*, expression of KaiA L156A severely disrupted rhythms: under LD, reporter expression oscillated similarly to the ΔN-KaiA L156A strain but with slightly lower amplitude and became arrhythmic under LL after ∼24 h (**Fig. 3b, Supplementary Fig. 8**). Thus, KaiA L156A compromises clock function while still stimulating KaiC autophosphorylation and light-driven class-1 gene expression.

### The isolated C-terminal domain enables autophosphorylation but cannot sustain rhythms

Several cyanobacteria, such as *Anabaena* PCC 7120, possess a KaiA that comprises the C-terminal domain alone ^28,29^. Having shown that N-KaiA and α5 govern the open–closed transition, we examined a C-terminal-only variant, KaiA 181–284 (C-KaiA), that mimics a constitutively open conformation as KaiA L156A does. C-KaiA efficiently stimulates KaiC phosphorylation (**Fig. 2c, Supplementary Fig. 3**) *in vitro* but fails to support stable oscillations (**Fig. 3a–b, Supplementary Fig. 4**), reinforcing that excessive stabilization of the open state is detrimental to rhythmicity. Measurements of reporter gene expression *in vivo* showed that the C-KaiA strain, like the KaiA L156A strain, did not sustain rhythms in LL, although it showed bioluminescence higher than the KaiA-null strain and lower than the KaiA L156A strain; under LD its reporter oscillated similarly to KaiA L156A but with lower amplitude. *S. elongatus* C-KaiA behaves more like a molten globule than a cooperatively folded protein ^21^, which may account for its lower amplitude. In the same assay, *S*. *elongatus* reporter strains with *Anabaena* KaiA (whose sequence begins near position 183 relative to *S. elongatus* KaiA) gave higher bioluminescence than did C-KaiA and a higher-amplitude LD rhythm, with an acrophase at the light-to-dark transition matching wild-type *S. elongatus* KaiA. However, it also failed to sustain rhythms under LL. Thus, single-domain KaiAs and the constitutively open KaiA L156A can recapitulate an hourglass-like clock in *S. elongatus* in a rhythmic environment but cannot support self-sustained oscillation, suggesting that additional species-specific features (perhaps an as-yet-undiscovered “KaiD”) are required for autonomous rhythmicity in single-domain KaiA systems.

### KaiA variants differentially affect the temperature dependence of *in vivo* versus *in vitro* experiments

Temperature compensation, maintenance of a near-24-h period across physiological temperatures, is a defining property of circadian clocks ^30,31^. In *S. elongatus*, KaiC and KaiB are both contributors: the KaiC ATPase is remarkably slow and temperature-insensitive ^32,33^, and KaiB likely contributes through its temperature-dependent free-energy landscape ^14^. We asked whether KaiA’s open–closed equilibrium is a third contributor by separately examining the temperature sensitivities of the KaiA-stimulated autophosphorylation rate and period in the reconstituted oscillator using our different KaiA constructs.

Across the panel of KaiA variants that span the closed–open axis, the temperature sensitivity of autophosphorylation was largely unchanged. ΔN-KaiA, which is biased toward the closed conformation, failed to stimulate autophosphorylation at any temperature, precluding a meaningful Q₁₀ determination. The remaining constructs gave similar values: ΔN-KaiA L156A, Q₁₀ = 1.3 ± 0.2; wild-type KaiA, Q₁₀ = 1.4 ± 0.1; the constitutively open KaiA L156A, Q₁₀ = 1.4 ± 0.2; and C-KaiA, Q₁₀ = 1.3 ± 0.1 (mean ± SD, n = 2 biological replicates; **Supplementary Fig. 3, Supplementary Table 9**). These values are not significantly different and are all modestly greater than 1, indicating that autophosphorylation is mildly temperature-sensitive, as was also shown previously ^24^, and that shifting KaiA’s open–closed set-point does not measurably change this sensitivity. In other words, the lack of a trend across the closed–open series argues that KaiA’s conformational equilibrium is not a mechanism by which the phosphorylation phase is temperature-compensated.

The reconstituted oscillator remained temperature-compensated with respect to period: ΔN-KaiA L156A supported oscillations with a period Q₁₀ of 0.98 ± 0.02 comparable to KaiA WT (Q₁₀ = 1.11 ± 0.05), albeit with a slightly longer period and lower amplitude (**Supplementary Fig. 4**). *In vivo*, however, the ΔN-KaiA L156A strain showed lower-amplitude rhythms, a shorter period, and a phase advance relative to KaiA WT at 30 °C (**Fig. 3b**); its period over-compensated across 27–35 °C (Q₁₀ = 0.78 ± 0.01) when *kaiA* was expressed at NS1 with native *kaiA* knocked out, versus Q₁₀ = 1.04 ± 0.01 in the control strain; **Fig. Supplementary Fig. 8**). Reporter gene rhythms from cyanobacteria when ΔN-KaiA L156A was expressed from the native locus produced still stronger over-compensation (Q₁₀ = 0.54 ± 0.03) but less robust, harder-to-interpret rhythms (**Supplementary Fig. 8**). This *in vitro*/*in vivo* discrepancy points to a second, cell-autonomous route to compensation that our reconstituted assays do not capture: robust timekeeping *in vivo* requires sensing of metabolism and redox state and feeding that information into the oscillator, in part through the N-terminal domain of KaiA and through CikA ^25,34^. Indeed, N-terminal KaiA point mutations produce over-compensated periods (Q₁₀ < 1) *in vivo* and heighten the clock’s sensitivity to redox perturbation ^25^, the same direction of effect seen for our N-terminally truncated ΔN-KaiA L156A strain. We therefore distinguish two routes by which a clock component can shape temperature compensation: first, an intrinsic route embodied by the temperature-insensitive KaiC ATPase, the KaiB fold-switch equilibrium, and the balancing of opposing temperature dependencies within the bistable switch ^24^; and second, a metabolic/redox-sensing route localized in part to the N-terminal domain of KaiA. Our in vitro data suggest that KaiA’s open–closed equilibrium is not a temperature-compensating element of the phosphorylation phase, while the *in vivo* behavior of the ΔN construct is consistent with the N-terminal domain contributing to compensation through metabolic and redox coupling.

KaiA is a major determinant of *S. elongatus* fitness under LD conditions ^35^. To assess how KaiA structure influences LD growth, we tested KaiA mutants under four temperatures and three light intensities (**Supplementary Fig. 12**). KaiA WT strain grew well under all conditions from 25°C at 150 µmol photons m^−2^ s^−1^ to 40°C at 75 µmol photons m^−2^ s^−1^, whereas strains carrying “locked” variants were sensitive to LD regimes in all conditions, regardless of whether the KaiA variants were locked in fully closed (C-KaiA) or fully open (ΔN-KaiA) conformations. In contrast, a fully open KaiA retaining the N-terminal domain (KaiA L156A) nearly complemented KaiA WT under standard growth conditions but resulted in LD sensitivity under high light and high temperature. Likewise, a variant that retains both open and closed conformations but lacks the N-terminal domain (ΔN-KaiA L156A) showed near-wild-type fitness under standard conditions, yet showed LD sensitivity under high light and high temperature.

### KaiA catalyzes KaiB–dpKaiC disassembly to restart KaiC autophosphorylation

In the canonical picture, KaiA is sequestered within KaiB–pKaiC assemblies at night and re-emerges passively as KaiC dephosphorylates. We asked whether KaiA plays an active role in remodeling KaiB–dpKaiC complexes near dawn. Our data support a model in which KaiA actively catalyzes KaiB–dpKaiC disassembly, initiating a new phosphorylation cycle before dawn.

### KaiB–dpKaiC complexes are long-lived at 30–35 °C without KaiA

FA of labeled KaiB showed that, in the absence of KaiA at 30–35 °C, KaiB binds dpKaiC with high apparent affinity and forms complexes that dissociate only slowly relative to the *in vitro* period (**Fig. 4**). Equilibrium binding yielded low-micromolar apparent dissociation constants (**Supplementary Fig. 13–14, Supplementary Table 10**). These complexes are therefore intrinsically stable at higher temperatures, and passive dissociation alone is too slow to account for the rapid morning transition to renewed autophosphorylation.

**Figure 4.**
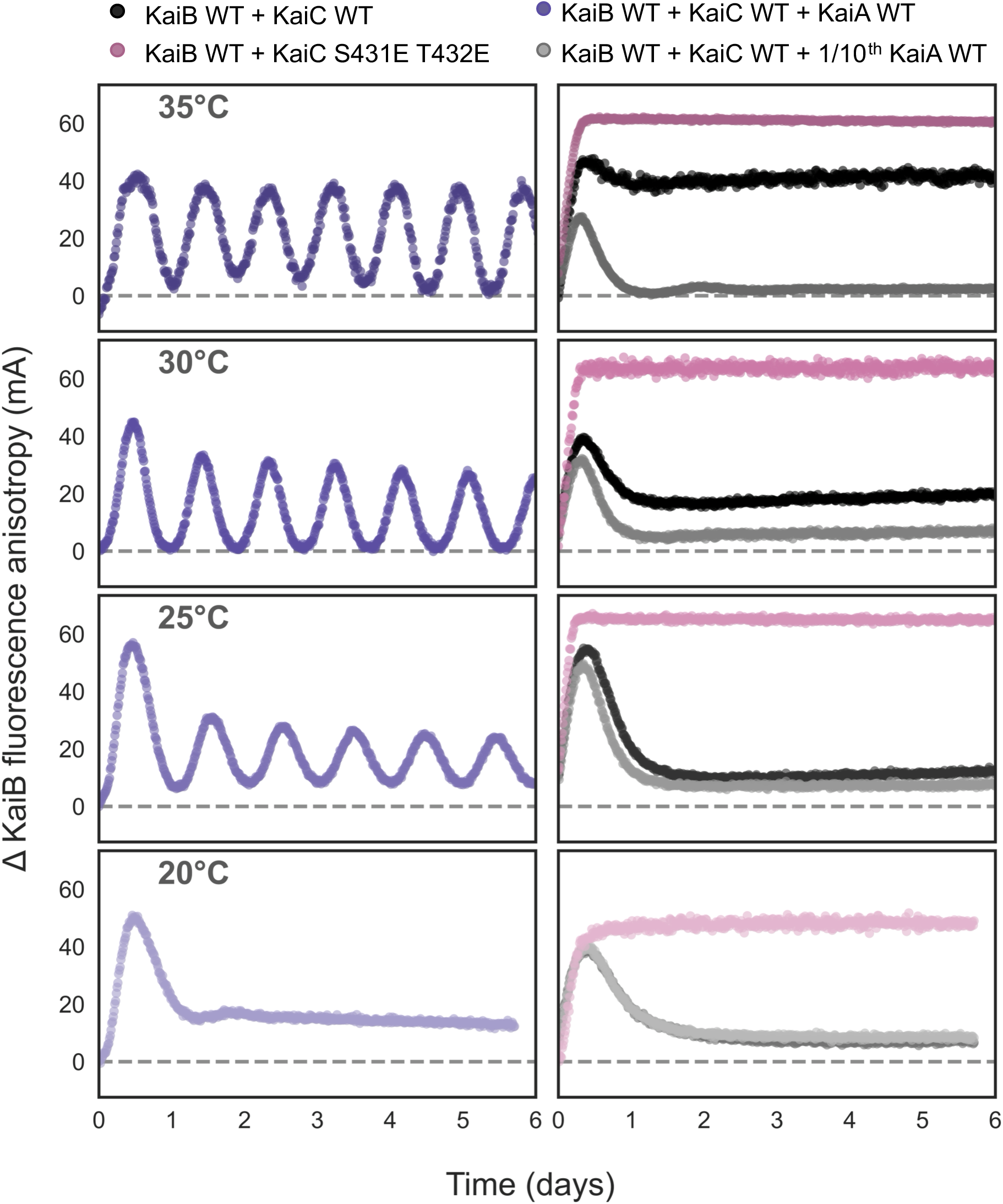
KaiA catalyzes disassembly of stable KaiB–KaiC complexes to restart KaiC phosphorylation. FA of labeled KaiB in the presence of KaiC mutants at different temperatures, where anisotropy was subtracted by that of KaiB WT (50 nM labeled KaiB with KaiB WT (3.45 μM)). The temperature decreases from 35°C in the top row to 20°C in the bottom row. **Left column**: FA of 50 nM labeled KaiB in the presence of KaiC WT (3.5 μM), KaiB WT (3.45 μM), and KaiA WT (1.2 μM). **Right column**: (Black) FA of 50 nM labeled KaiB in the presence of KaiC WT (3.5 μM) and KaiB WT (3.45 μM). (Grey) FA after adding substoichiometric KaiA WT to the KaiB WT–KaiC WT complex. (Pink) FA of KaiB in the presence of phosphomimic KaiC S431E T432E. The grey dashed line in each plot represents the anisotropy levels of 50 nM labeled KaiB with KaiB WT (3.45 μM) at each temperature.

### Substoichiometric KaiA accelerates KaiB–dpKaiC disassembly

Adding a low concentration of KaiA (1/10th of the standard 1.2 µM) to preformed KaiB– dephosphorylated KaiC complexes (dpKaiC) caused FA to decrease markedly over time, indicating KaiA acting catalytically to release KaiB (**Fig. 4**). Concomitantly, KaiC phosphorylation began to rise as complexes disassembled (**Supplementary Fig. 15-16**). Thus, KaiA binds KaiC to both stimulate autokinase activity and catalytically dismantle KaiB–dpKaiC complexes. Mechanistically, this dismantling is part of a nucleotide-exchange event gated by the KaiC CI domain: ADP release is the rate-limiting step of the KaiC ATPase cycle, and KaiA must accelerate ADP-to-ATP exchange within CI to cause KaiB dissociation at the cusp of the phosphorylation phase ^36,37^. Consistent with an allosteric route from the KaiC C-terminus to the distant KaiB-binding site, a KaiC truncation lacking the flexible C-terminal tail (KaiC 1–505), which weakens direct KaiA binding ^38^, showed impaired KaiA-mediated disassembly (**Supplementary Fig. 16**). Together with prior work on CI–CII coupling across nearly 10 nm ^38,39^, these findings indicate that KaiA destabilizes the complex by engaging the KaiC C-terminal tails, providing a long-range allosteric mechanism for actively disassembling KaiB–dpKaiC complexes to restart autophosphorylation each dawn.

### The KaiB–dpKaiC interaction and KaiA-mediated disassembly are both temperature-gated

The *S. elongatus* clock is temperature-compensated across a broad range, yet ensemble rhythms fail near and below 20 °C ^40,41^. We therefore examined how both processes, KaiB– dpKaiC interaction and KaiA-mediated disassembly, change with temperature. FA measurements of KaiB binding to dpKaiC revealed a pronounced temperature dependence: at 30–35 °C, KaiB bound with low-micromolar apparent dissociation constants and the anisotropy signal indicated a stable complex, whereas at 20 °C the FA signal decreased (**Fig. 4**), approaching the level of free KaiB. Because the period trend in Tris buffer is also influenced by pH ^42^, we confirmed that these FA changes are not significantly affected by pH (**Supplementary Fig. 17**); a phosphate buffer with a temperature-independent p*K*a was avoided because it yielded a significantly less robust IVO in our hands. By contrast, the late-night phosphomimetic KaiC (S431E/T432; KaiC-ET) sustained high KaiB binding across 20–35 °C (**Supplementary Fig. 15**), and preformed KaiB–KaiC-ET complexes did not disassemble appreciably upon adding 0.1× KaiA at any temperature tested. The temperature dependence is thus specific to the dephosphorylated complex. Adding 0.1× KaiA to KaiB + dpKaiC preincubated at 30–35 °C drove FA down to the level of free KaiB, but at 20 °C little complex remained on which KaiA could act (**Fig. 4**). These observations suggest that at 30–35 °C, when S431 finally dephosphorylates at the night-to-dawn transition, un-sequestered substoichiometric KaiA catalytically disassembles the otherwise stable complexes, producing an autocatalytic feedback loop as more free KaiA is released (note that some amount of free KaiA exists at every phase of the oscillator ^43^; we postulate that this mechanism supports robust rhythms at 30–35 °C. Near 20 °C, by contrast, the KaiB–dpKaiC complex is intrinsically too weak to require KaiA for its dissolution, so the KaiA-catalyzed autocatalytic step is bypassed. Together, these measured temperature-dependent processes, the weakening of the KaiB–dpKaiC complex, the KaiA-catalyzed autocatalytic disassembly, and the temperature-dependent KaiB fold-switch that supplies binding-competent KaiB suggested a molecular basis for the temperature-dependent failure of the oscillator, which we set out to test in a kinetic model.

### The Autocatalytic Dawn (AD) model reproduces the amplitude collapse and oscillator death

We first asked whether the temperature-dependent processes we observed, the weakening of the KaiB–dpKaiC complex, the KaiA-catalyzed autocatalytic disassembly, and the temperature-dependent KaiB fold-switch could, on their own, account for the oscillator’s behavior by incorporating them into a model we named the Autocatalytic Dawn (AD) model (**Supplementary Fig. 18**). The model reproduced the decline in amplitude with cooling and the virtual loss of rhythm at 20 °C, with low-temperature failure occurring as deterministic oscillator death, the individual limit cycle collapsing through a Hopf bifurcation (**Fig. 5**).

**Figure 5.**
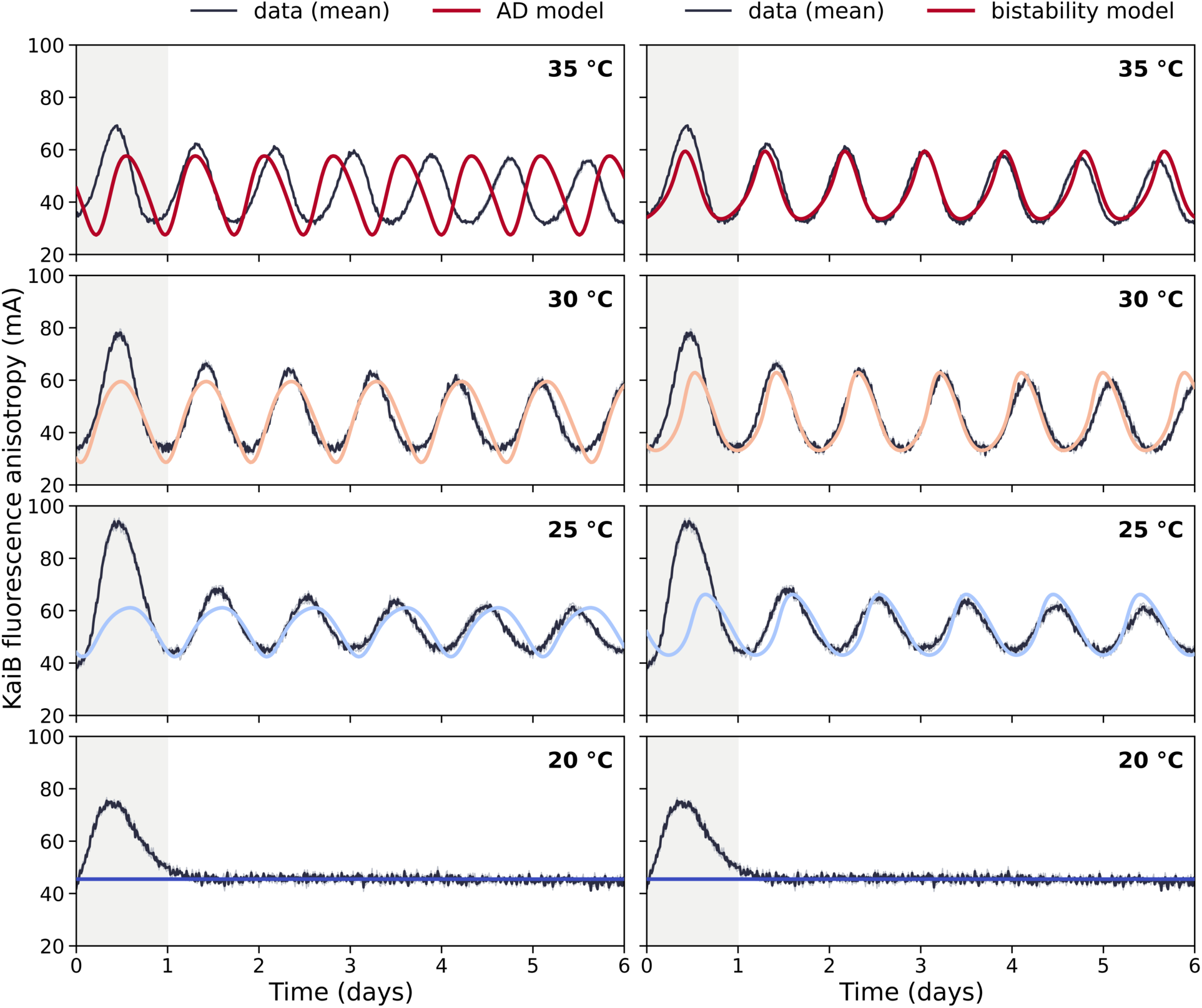
Temperature-dependent AD model and bistability model recapitulate the reconstituted oscillator across temperature. FA of KaiB in the reconstituted oscillator (IVO) at 35, 30, 25, and 20 °C (grey, individual replicates; dark, replicate mean) overlaid with the Autocatalytic Dawn (AD) model (Left panel) and KaiA-KaiB bistability model (Right panel). The shaded first 24 h is the post-mixing equilibration period and is excluded from fitting.

Different temperature-dependent processes act together to produce this collapse. First, the KaiB–dpKaiC complex weakens as temperature decreases: the intrinsic off-rate of the dephosphorylated complex rises considerably between 35 and 20 °C (**Supplementary Fig. 13**), so that at low temperature these complexes dissolve spontaneously rather than persisting until free KaiA disassembles them. Second, this weakening undermines the KaiA-catalyzed autocatalytic feedback that sharpens the dawn transition. At 30–35 °C, where the dephosphorylated complex is long-lived, substoichiometric free KaiA initiates a cooperative disassembly cascade; each KaiA released by one dissolving complex accelerates the disassembly of the next, producing a switch-like dawn transition (**Fig. 4**). At 20 °C, spontaneous dissociation overwhelms the cascade (the ratio of autocatalytic to intrinsic off rates falls from ∼1.3 to ∼0.14), so the positive feedback that sustains oscillation amplitude is effectively lost. Third, the AD model incorporates the temperature-dependent shift of the gsKaiB ⇌ fsKaiB equilibrium toward fsKaiB at low temperature ^14^. With the gsKaiB ⇌ fsKaiB equilibrium held temperature-independent, the modeled 20 °C oscillation persists at roughly three times the measured amplitude (**Supplementary Fig. 18**), confirming that the fold-switch is a significant contributor; however, no single factor alone accounts for the full amplitude collapse.

The period of the AD model is temperature compensated in the broad sense (frequency Q₁₀ ≈ 1.3) over 25–35 °C, within the 0.8–1.4 range characteristic of circadian clocks, though less tightly than the oscillator itself (Q₁₀ ≈ 1.1) and, notably, the period is ∼11.5% shorter at 35 °C (≈18.15 h versus 20.5 h observed) ^44^. The observed processes are thus sufficient to generate the amplitude collapse and oscillator death; capturing the tight temperature compensation and the observed periods led us to examine a second, independent model.

### A temperature-dependent bistability model recapitulates the temperature dependence of the reconstituted oscillator

This second model is built on a bistability in the protein-interaction network. KaiA’s antagonism of the KaiB–pKaiC interaction, inhibiting complex assembly and stimulating its disassembly together with the sequestration of KaiA by the complex, provides the ingredients of a double-negative (net positive) feedback loop that renders the KaiA–KaiB–pKaiC network bistable ^24^. In this bistability model, KaiA regulates KaiB binding to pKaiC, and the oscillation is a relaxation cycle in which slow KaiC autophosphorylation drives the system across a hysteretic switch whose width sets the amplitude. Because KaiA’s disassembly of the KaiB–pKaiC complex, the rate at the heart of the positive feedback, is strongly temperature dependent (Q₁₀ ≈ 3.5), the width of the bistable region should itself vary with temperature. We therefore asked whether the model can quantitatively account for the temperature dependence of our reconstituted oscillator.

We adopted the published six-state kinetic model of the KaiABC network that couples ordered CII phosphorylation to KaiA-regulated KaiB binding ^24^, in which free KaiA both inhibits KaiB– pKaiC assembly and stimulates KaiB–pKaiC disassembly, and computed the model’s KaiB– pKaiC complex concentration as the analog of our observable FA (*Materials and Methods*). Without adjusting any of the model’s mechanistic parameters, we mapped its complex output onto empirical FA units with a single global amplitude scale and set the overall timescale with a single global rate-scale factor.

The model reproduced all three signatures of our data **(Fig. 5).** It remained temperature-compensated in period, with a frequency Q₁₀ of 1.09 across 25–35 °C. Its FA amplitude declined progressively with cooling and tracked the measured traces closely, with residuals ≤ 3.8 mA against ≈25-35 mA oscillations and correlations of 0.91–0.94 at 25–30 °C. At 20 °C, the model’s oscillation vanished: the bistable region, computed directly from the KaiA-regulated binding parameters, narrows roughly sevenfold between 30 and 23 °C and closes entirely at 20 °C, so that no stable limit cycle exists and the FA signal flatlines, reproducing the arrhythmia we observe.

Two aspects of this result require emphasis. First, the model reproduces the loss of rhythm at 20 °C deterministically: the failure is an amplitude collapse caused by the closing of the bistable switch. Second, the bistable region is set by the KaiA-regulated KaiB-binding rates, so its temperature-dependent narrowing, and thus the change in oscillator amplitude, arises from the temperature dependence of those binding rates (the phosphotransfer rates, which are also temperature-dependent, set the reaction speed).

## Discussion

Our work reframes KaiA from a purely daytime activator into a factor that acts at both ends of the KaiC phosphorylation cycle. First, the α5 helix links KaiA’s N- and C-terminal domains in a domain-driven tug-of-war that tunes the balance between an open, KaiC-activating conformation and a closed, KaiC binding-occluded conformation. This balance sets how strongly KaiA activates KaiC during the day and how readily KaiA can be sequestered at night. Perturbing this balance in either direction, by removing the N-terminal domain or weakening α5–C-terminal contacts, disrupts sustained rhythmicity. Second, substoichiometric KaiA catalytically dismantles near-dawn KaiB–dpKaiC complexes, and this activity requires the KaiC C-terminal tails. While the autonomous post-translational oscillator is generated by ATP-driven protein-interactions, *in vivo* timekeeping additionally relies on sensing intracellular redox and metabolism through KaiA and CikA ^25,34^. Truncating the N-terminal domain disrupts this metabolic integration, destabilizing the clock *in vivo* even where the reconstituted oscillator remains temperature-compensated *in vitro*. The N-terminal domain is critical for synchronization and robustness *in vivo* as well as *in vitro*. *In vivo*, KaiA N-terminal metabolic/redox integration buffers KaiA activity against environmental changes, maintaining organism fitness under various LD conditions. Strains carrying locked open or closed KaiA variants that lack the N-terminal domain are LD sensitive. In contrast, strains carrying KaiA that can adopt both open and closed conformations, or that are locked open but carry the N-terminal domain, show near-wild-type fitness under standard LD conditions; however, those strains become LD sensitive under high light and high temperature. Our modeling followed two independent routes that converge on the same conclusion. The AD model, built upon our temperature-dependent FA data, and a recently developed bistability model of the KaiA–KaiB antagonism ^24^ both reproduce the decline in amplitude with cooling and the loss of rhythm at 20 °C. That two mechanistically distinct models reach the same conclusion strengthens the inference that the cold failure is deterministic oscillator death. The two differ in how precisely they capture the oscillator’s timing: both are temperature compensated in the broad sense (period Q₁₀ within the 0.8–1.4 range typical of circadian clocks ^44^. The bistability of the Liu et al. model arises from KaiA’s regulation of KaiB binding to pKaiC, and the temperature dependence of those KaiA-regulated binding rates sets the width of the bistable region, and hence the amplitude. The model’s temperature dependence has a second source: the dephosphorylation of Ser431 (Q₁₀ ≈ 5.8), which sets the reaction speed. Temperature compensation of the period emerges from the balance between these two effects. The model does not include the KaiB–dpKaiC complex, the temperature-dependent weakening of that complex, the autocatalytic KaiA-catalyzed disassembly, or the temperature-dependent KaiB fold-switch that our biochemistry reveals. Our data identify distinct, temperature-sensitive steps: the dawn dissolution of the KaiB–dpKaiC complex and its autocatalytic acceleration by free KaiA, together with the shift of the KaiB fold-switch equilibrium toward the fold-switched state at low temperature, which further reduces free KaiA through inhibition. The fact that both models, each emphasizing different temperature-dependent processes, can recapitulate the experimental data suggests that all of these processes contribute to the overall mechanisms that underlie temperature compensation and the low-temperature failure of the oscillator.

## Materials and Methods

Full experimental procedures for protein expression and purification, fluorescent labeling **(Supplementary Table 4-5)**, SEC-MALS **(Supplementary Table 11)**, solution and ¹⁹F NMR **(Supplementary Table 8)**, KaiC A-loop binding assays **(Supplementary Table 7)**, KaiB–KaiC equilibrium binding assays, *in vitro* oscillator reconstitution, the Autocatalytic Dawn (AD) model, bistability model, cyanobacterial strain construction **(Supplementary Table 1-3)**, and bioluminescence recording and analysis are provided in the SI Appendix, Materials and Methods.

## Acknowledgments

This work was supported by the US National Institutes of Health (NIH) grant R35GM144110 and US Army grant W911NF-23-1-0248 awarded to AL, and R35GM118290 to S.S.G. We thank D. Rice for maintaining the NMR facility at UC Merced. The KaiABC schematic cycle in Fig. 1A was created using BioRender.com^45^.

## Author Contributions

A.L., A.C.C., A.T., M.F., L.M.V.P., L.F.L., and S.S.G. conceptualized the project. A.C.C., A.G.C., N.Z., and M.F. designed and performed the biochemical assays. A.A.G.C., A.T., and C.A.O. designed and performed the *in vivo* assays. A.C.C., A.T., and M.F. analyzed and interpreted the experimental data. A.C.C., A.T., M.F., S.S.G., and A.L. wrote the manuscript.

