## Supporting information for "A KaiA Conformational Switch Links Day-Night Transitions to Clock Robustness"

**This file includes:**

Materials and Methods

Supplementary Figures 1 to 19

Supplementary Tables 1 to 11

References

Supplementary Figure 18. Schematic representation of the Autocatalytic Dawn (AD) model. ... 54
Supplementary Figure 19. Schematic representation of genetic modifications introduced into the S. elongatus chromosome for the construction of the KaiA mutant strains used in this study. .. 56

#### Materials & Methods

##### Cloning, expression, Protein Purification, and Fluorescent Labeling.

All the protein constructs used in the in vitro experiments are listed in **Supplementary Table 4.** For KaiA, KaiB, and KaiC, their overexpression and purification were performed as previously reported [1] in BL21(DE3) *Escherichia coli* (Novagen). Cells were harvested and lysed using an Avestin C3 Emulsiflex homogenizer (Avestin Inc, Canada). Cell lysate was clarified by centrifugation before affinity purification on Ni-NTA resin (QIAGEN) in polypropylene gravity columns using the buffers specified in **Supplementary Table 5**. All purification steps were carried out at 4°C with freshly prepared buffers. For Strep-tag KaiC purification, Strep-Tactin flow resin gravity columns were used. The KaiC proteins were eluted with 50 mM Biotin. [2] For cleavage of the 6x-His-SUMO fusion protein, 6x-His-ULP1 protease was added and incubated at 4 °C for 15 hours. 6x-His-SUMO was removed by loading the cleaved protein onto a Ni-NTA column, followed by purification by gel-filtration chromatography. For KaiA, KaiB, and KaiC, clock reaction buffer (20 mM Tris, 150 mM NaCl, 1 mM ATP, 5 mM $MgCl_2$, 0.5 mM EDTA, pH 8.0) was used as the final elution buffer for the gel-filtration chromatography. For fluorescent labeling of the KaiB K25C protein, the elution buffer was 20 mM Tris, 150 mM NaCl, pH 7.5. The protein was reduced with 20 mM tris(2-carboxyethyl) phosphine (TCEP) before labeling. The fluorophore thiol dye 6-iodoacetamidofluorescein (6-IAF) (Invitrogen) was mixed in a 5:1 dye-to-protein ratio and incubated overnight at 4°C. Labeled protein fractions were purified further by gel-filtration chromatography using a Sephadex 75 16/600 column in clock buffer. [1,3]

**SEC MALS**

Size-exclusion chromatography coupled with multi-angle light scattering (SEC-MALS) was performed using a Cytiva FPLC system coupled in-line to a multi-angle light scattering detector (miniDAWN, Wyatt Technology) and a differential refractive index detector (Optilab). Protein samples were filtered through a 0.22 $\mu$m filter before being injected into the columns. All experiments were performed on a Superdex 200 10/300 GL column equilibrated in clock reaction buffer (20 mM Tris, 150 mM NaCl, 1 mM ATP, 5 mM MgCl2, 0.5 mM EDTA, pH 8.0) at a flow rate of 0.5 ml/min at room temperature. Molar mass was calculated from the SEC-MALS data using the instrument software Astra with $dn/dc = 0.185$ mL/g for protein samples. Peak regions corresponding to homogeneous species were selected for analysis, and together, this combined MALS/QELS/dRI approach provided absolute molar mass, RMS radius, and hydrodynamic radius across the elution peak without reliance on UV extinction coefficients. (**Supplementary Table 11)**

**Reconstituting in vitro oscillations using fluorescence anisotropy**

The reconstitution of the in vitro clock reactions was prepared in a 20 mM Tris, 150 mM NaCl, 1 mM ATP, 5 mM MgCl2, 0.5 mM EDTA, pH 8.0 buffer by mixing unlabeled proteins at the final concentrations as KaiC WT (3.5 µM), KaiB WT (3.45 µM), KaiA mutants (1.2 µM), and 50 nM fluorescently labeled KaiB probe. After filtering through a 0.22 $\mu$m filter, aliquots (70 µL) of reaction mix were pipetted into black, non-binding 384-well microplates (Greiner BioOne). The plates were sealed with a clear adhesive film to avoid evaporation. All reactions were measured in three replicates per run. The fluorescence polarization was measured pre-equilibrated (at the experimental temperature) in a CLARIOstar BMG Plate Reader. The target polarization was set to 10% to a maximum polarization of 200 milli-polarization units. Fluorescence polarization was measured in kinetic mode every 15 minutes for 900 cycles. All the anisotropy values are reported in millianisotropy (mA) units. Period and amplitude analysis were performed as previously reported [1,3].

**KaiB-KaiC Equilibrium binding assays**

KaiB equilibrium binding assays were performed in clock buffer (20 mM Tris, 150 mM NaCl, 1 mM ATP, 1 mM $MgCl_2$, and 0.5 mM EDTA, pH 8). 60 mL of KaiB probe mix (50 nM fluorescently labeled KaiB and 3.45 µM KaiB WT) was pipetted into 22 adjacent wells of a 384-well flat-bottom black polystyrene assay plate (Corning). Master mixes were prepared by adding KaiB probe mix to thawed KaiC WT aliquots. Serial dilution was performed by transferring 30 µL of KaiC master mix to the first well, mixing and pipetting up and down more than 8 times, then transferring 30 µL to the subsequent well and repeating the process for all 22 wells. A multi-channel pipette was used for the serial dilution assay, and three replicate measurements were collected. After sealing, the plate was transferred to a pre-equilibrated CLARIOstar multimode microplate reader (BMG Labtech), and Fluorescence polarization was measured in kinetic mode every 15 minutes for 48 hours. [1,4] Similar assays were conducted with KaiC S431E T432E as well at 20°C, 25°C, 30°C, and 35°C. [5]

The fitting model quantifies KaiB–KaiC binding at each time point using a five-parameter "bell-shaped" equation that combines a Hill-type saturable binding term with a high-concentration attenuation (prozone) term:

$$FA(C) = base + A \cdot [C^n / (Kd1^n + C^n)] \cdot [Kd2 / (Kd2 + C)]$$

Here, base is the unbound baseline anisotropy, A the binding amplitude, Kd1 the apparent half-maximal binding concentration (*µ*M monomer), n the Hill coefficient describing cooperativity, and Kd2 a second dissociation constant that captures the decline in signal at high KaiC concentration (self-association or prozone effect). This isn't a mechanistic ODE model of the KaiA/KaiB/KaiC oscillator; it's a static algebraic binding isotherm applied independently at each sampled time point along the circadian time course, so cooperativity and affinity are tracked as they evolve rather than being derived from reaction kinetics. [1,4,6,7]

Parameters were estimated by nonlinear least squares with a custom Levenberg–Marquardt algorithm (numerically estimated Jacobian, adaptive damping, up to 300 iterations, convergence tolerance $1\times10^{-8}$ on the parameter step), minimizing the sum of squared residuals between measured and predicted anisotropy across the KaiC concentration series at each time point. [8,9] To avoid local minima, each fit was seeded from multiple starting points (a warm start from the previous time point's converged parameters plus three fixed initial guesses spanning a range of Kd1 and n), with parameters constrained to physically valid ranges (A, Kd1, Kd2 > 0; 0.3 < n < 25) and the best valid candidate retained by lowest SSE. Goodness of fit was tracked as $R^2 = 1 - SS\_res/SS\_tot$. Repeating this at every time point yields, per condition (variant × temperature × replicate), full time trajectories of Kd1, n, and $R^2$, showing how apparent binding affinity and cooperativity change over the circadian cycle. [10,11] (**Supplementary Table 10**) The Python plotting scripts used to generate these figures were written with the assistance of Claude Anthropic.

##### KaiC A-loop anisotropy binding assay

KaiC A-loop binding assays were performed in clock buffer (20 mM Tris, 150 mM NaCl, 1 mM ATP, 1 mM $MgCl_2$, and 0.5 mM EDTA, pH 8). The N-terminal 5-FAM (5-carboxyfluorescein)-labeled KaiC A-loop peptide from residues 488-519 was ordered from GenScript (RIISGSPTRITVDEKSELSRIVRGVQEKGPES). 25 nM labeled peptide was used for the assays with increasing concentrations of KaiA mutants. Reaction mixes were pipetted into a 384-well flat-bottom black polystyrene assay plate (Corning), and sealed with transparent film (MicroAmp™, Applied Biosystems™). After sealing, the plate was transferred to a Spark 10 M (Tecan) plate reader, and Fluorescence polarization was measured using 485 ± 20 nm and 535 ± 25 nm excitation and emission filters. The assay was repeated in triplicate at 20°C, 30°C, and 40°C.

Fluorescence polarization data were analyzed in Python using nonlinear least-squares fitting. Polarization values were converted to anisotropy as

$$r = \frac{2P}{3 - P}$$

and, for each KaiA concentration, three replicate anisotropy values were averaged to obtain a mean response; variability was summarized as the SEM. Binding isotherms were fit to a two-state binding model in which the concentration of peptide–KaiA complex is

$$[PL] = \frac{(K_d + P_t + 2A_t) - \sqrt{(K_d + P_t + 2A_t)^2 - 8A_tP_t}}{2}$$

and anisotropy is described by

$$r(A_t) = r_f + (r_b - r_f)\left(\frac{[PL]}{P_t}\right).$$

Here, $A_t$ denotes total KaiA concentration, $P_t$ denotes total peptide concentration, $K_d$ is the apparent dissociation constant, and $r_f$ and $r_b$ are the anisotropy values of the free and bound peptide, respectively. Fits were obtained with bounded nonlinear least squares (curve_fit), and parameter standard errors were estimated from the covariance matrix of the fit. Residuals were defined as $r_{\mathrm{obs}} - r_{\mathrm{fit}}$, and 95% confidence bands for the fitted curves were computed by propagating parameter uncertainty using the covariance matrix and the appropriate Student-$t$ critical value. The dissociation constants calculated are listed in **Supplementary Table 7.**

**The Autocatalytic Dawn (AD) model and Temperature-dependent bistability model of the reconstituted oscillator**

Two deterministic kinetic models were compared with the KaiB fluorescence-anisotropy (FA) time courses: the Autocatalytic Dawn (AD) model developed here (Fig. S18) and the bistability model of Liu et al. (2026). Both were implemented in Python and integrated with SciPy. Model equations, parameter values, code, and data are available on Zenodo[12]**.** The AD model describes the KaiC phosphorylation cycle and its binding to KaiB (**Supplementary Figure 18**). KaiC is phosphorylated from the unphosphorylated state (U) to the Thr432-phosphorylated (T) and doubly phosphorylated (ST) states. It is then activated by the CI step to a KaiB-binding-competent form, binds fold-switched KaiB, and is dephosphorylated within the complex before KaiB is released and the cycle restarts. Phosphorylation and dephosphorylation rates depend on free KaiA as described by Rust et al. (2007)[13], with basal rate constants taken from Rust et al. (2007) [13] and Chang et al. (2015)[14]. Each KaiB-bound complex sequesters KaiA, so free KaiA falls as complexes accumulate. Free KaiA in turn accelerates KaiB release, which makes the dawn reset autocatalytic. KaiB interconverts between its ground state and the binding-competent fold-switched state with temperature-dependent rates. Concentrations matched the experiments (KaiC 3.5 $\mu$M, KaiB 3.5 $\mu$M, KaiA 1.2 $\mu$M). In-complex dephosphorylation was given a $Q_{10}$ of 2.6 below 30 °C (Liu et al., 2026)[15]. Ten parameters were fitted to the oscillation amplitudes at the four temperatures: a global rate factor, the KaiA feedback strength, the $Q_{10}$ of the CI step, the fold-switch rates and their Q10 and the KaiB release rate at 30, 25, and 20 °C.

Period was not used in the fit. The fitted KaiB release rate at 30 °C (0.14 $h^{-1}$) is higher than the measured disassembly rate of the KaiB–dephosphorylated KaiC complex (0.0875 $h^{-1}$; Simon et al., 2022[16]). To test whether the KaiB fold switch is required for the loss of rhythm at 20 °C, the fold-switch rates were made temperature independent (constant KaiB supply) with all other parameters unchanged.

The bistability model is a Python implementation of the six-state model of Liu et al. (2026)[15], built from their reported equations and parameter values at 20 and 30 °C. KaiA-dependent phosphorylation and dephosphorylation are coupled to KaiB binding through KaiA sequestration. Parameters at other temperatures were obtained by linear interpolation between 20 and 30 °C and by linear extrapolation to 35 °C, and no mechanistic parameter was refitted.

Plate-reader fluorescence polarization was converted to anisotropy (mA = 2000·mP/(3000 − mP)). Each temperature was measured in triplicate and the mean trace was used, excluding the first 24 h after mixing. Oscillation amplitude was defined as twice the standard deviation of the detrended trace, and period from its autocorrelation. The model FA signal is the amount of KaiB-bound KaiC. In the AD model, the KaiB–dephosphorylated KaiC complex is weighted by its relative anisotropy, which was measured at each temperature from steady-state FA references. All computations used Python with NumPy, SciPy, pandas and Matplotlib. Claude (Anthropic) was used to assist in verifying the model code, checking parameter sources, and writing and testing the analysis scripts. All code and results were reviewed by the authors, and are available on Zenodo.[12]

#### KaiA-dependent KaiC autophosphorylation assays

For phosphorylation assays, KaiC protein was dephosphorylated overnight at 30°C in clock buffer. The assay began by adding 1.2 μM KaiA protein to 3.5 μM KaiC and transferring samples to preincubated water baths at 20°C, 25°C, 30°C, and 35°C. Aliquots (10mL) were collected every 30 minutes for 4 hours. Each aliquot was mixed with 10 mL 2X SDS-PAGE loading buffer, spun for 20 seconds, and stored at -20°C. Frozen aliquots were thawed and loaded with 5 mL of samples onto freshly made Phos-tag gels. The electrophoresis apparatus was kept in an ice box during the entire run. The phosphorylation rates calculated are listed in **Supplementary Table 6.**

#### KaiA mutants 15N-1H HSQC NMR spectroscopy

Uniformly $^{15}$N-labeled ΔN KaiA and ΔN KaiA L156A were expressed in *E. coli* grown in M9 minimal medium prepared with $^{15}NH_4Cl$ as the sole nitrogen source and 99.8% $D_2O$ (v/v). The purified protein was concentrated to 50 *μ*M in NMR buffer containing 20 mM Tris, 50 mM NaCl, pH 7.0, 10 *μ*M DSS (4,4-dimethyl-4-silapentane-1-sulfonic acid) as an internal chemical shift reference, and 0.02% NaN3 to prevent microbial growth, prepared in 95% H2O/5% D2O. All NMR experiments were performed on a Bruker AVANCE III spectrometer operating at 600 MHz $^{1}$H Larmor frequency, equipped with a triple-resonance cryoprobe (TCI) to enhance sensitivity[17].

#### Expression and purification of $^{19}$F Tryptophan-labeled KaiA mutants

For $^{19}$F NMR of KaiA mutants (N-KaiA, KaiA WT, KaiA L156A), 7-F-tryptophan was labeled based on [18] with modifications. KaiA pET-28b plasmids were expressed in *Escherichia coli* BL21(DE3) cells (Novagen). Cells were grown in M9 (H2O) medium with N-15-enriched $NH_4Cl$ at 37°C, 220 rpm, until OD 600 reached 0.6. Shaker temperature was lowered to 18°C, and cells were shaken for 1 hour at 220 rpm. 60 mg of 7-fluoroindole dissolved in 200 µL DMSO was added, and cells were shaken for another hour. Induction with 0.5 mM IPTG at 18°C was performed for 18 hours. Cells were harvested by centrifugation at 5,000 rpm for 10 min, followed by cell lysis and protein purification. Proteins were purified by Ni-NTA affinity chromatography and gel filtration chromatography as described in **Supplementary Table 6**. For NMR experiments, final samples contained ~ 100 µM protein in 20 mM sodium phosphate buffer, 150 mM NaCl, pH 8, 5 mM DSS, 0.02% $NaN_3$, 5% $D_2O$. A Bruker 600 MHz AVANCE III spectrometer equipped with a TCI cryoprobe was used for all the NMR experiments described here. Chemical shifts were referenced to internal 2,2-dimethyl-2-silapentane-5-sulfonate. All data were processed and analyzed with Topspin 3. (**Supplementary Table 8)**

**Culture conditions, strains, plasmids and molecular methods for in vivo experiments**

Growth conditions and strain construction largely followed established protocols described previously [19] and summarized here (**Supplementary Fig. 19, Supplementary Table 1,2,3**). *Escherichia coli* strains were cultured in LB medium at 37 °C, either in liquid culture or on agar plates, with antibiotic selection as appropriate. *Synechococcus elongatus* PCC 7942 and derivative strains were grown in BG-11 medium [20] at 30 °C, unless otherwise noted, either in shaking BG-11 liquid cultures or on BG-11 agar plates supplemented with antibiotics when appropriate, under constant illumination or 12 h:12 h light–dark cycles at light intensity ranging between 50 and 150 *µ*mol photon $m^{-2}$ $s^{-1}$.

Recombinant *S. elongatus* strains were generated primarily by natural transformation [21]. Briefly, cultures at an optical density at 750 nm ($OD_{750}$) of 0.3-0.5 were concentrated (~ 5 times), washed, and incubated with plasmid DNA overnight in the dark, then spread on selective BG-11 agar plates. Colonies were isolated after ~5 days of growth under constant illumination of 100 *µ*mol photon $m^{-2}$ $s^{-1}$.

Ectopic expression of the *kaiA* alleles was established prior to deletion of the native locus (**Supplementary Fig. 17**) in *S. elongatus* AMC2747 that expresses the firefly luciferase gene from the class 1 P*kaiBC* promoter (P*kaiB*-*luc*) [22].

CRISPR editing in the native *kaiA* locus were conducted in *S. elongatus* AMC2752 carrying P*kaiBC*-*luc* at NS2 and whose *ago* gene has been deleted. The Argonaute nuclease (SeAgo) in *S. elongatus* was previously shown to reduce natural transformation and impede the maintenance of RSF1010-based plasmids required for CRISPR editing [22].

For CRISPR editing in the native *kaiA* locus, plasmids carrying the *kaiA* alleles were first introduced by natural transformation into *S. elongatus* AMC2752 [22]. After incubation in darkness with plasmid DNA, the cells were used directly as recipients for a subsequent biparental conjugation to introduce the cargo plasmid carrying the *cpf1* nuclease and crRNA targeting the native *kaiA* allele. Conjugation followed previously published methods [22,23] from *E. coli* donor strain AM1359, which harbors the conjugal plasmid pRL443 and the helper plasmid pRL623 for mobilization of ColE1 plasmids. Briefly, AM1359 donor cells carrying the cargo plasmid were washed 3 times, mixed with the recipient cells and spotted onto BG-11 agar supplemented with 5% (v/v) LB and incubated at 30°C under low light (10–20 $\mu$mol photons $m^{-2}$ $s^{-1}$) for 24 h, after which cells were recovered and plated on BG-11 agar with selective antibiotics until isolated colonies appeared (5–9 days).

Plasmids pAM5903–pAM5907, used to ectopically express the *kaiA* alleles at NS1, were constructed by seamless assembly following the CyanoVECTOR cloning design [24]. Briefly, plasmid pCVD020 (carrying homologous DNA fragments for recombination at the neutral site 1, NS1TC) was digested with ZraI, and plasmid pCVD064 (carrying the nourseothricin resistance gene *nat*, codon-optimized for *S. elongatus*, *nat7942*) was digested with EcoRV to generate the vector backbone and resistance marker fragments, respectively. The P*kaiA* promoter and different *kaiA* alleles: P*kaiA*-*kaiA*(aa 148-284), P*kaiA*-*kaiA*(aa 148-284, L156A), P*kaiA*-*kaiA*(aa 181-284), P*kaiA*-*kaiA*(aa L156A), and P*kaiA*–*kaiA* were excised by EcoRV digestion from the pCVD015 derivatives described below: pAM5832, pAM5833, pAM5834, pAM5847, and pAM5848. The digested fragments were then assembled by seamless ligation with the NS1TC backbone with the NAT7942 selection marker to produce pAM5903 (P*kaiA*-*kaiA*(aa 148-284)), pAM5904 (P*kaiA*-*kaiA*(aa 148-284, L156A)), pAM5905 (P*kaiA*-*kaiA*(aa 181-284)), pAM5906 (P*kaiA*-*kaiA*(aa L156A)), and pAM5907 (P*kaiA*–*kaiA*).

The pCVD015 derivatives carrying the P*kaiA* promoter driving the different *kaiA* alleles were constructed by PCR amplification followed by seamless assembly into pCVD015 digested with SwaI. All fragments were amplified from *S. elongatus* genomic DNA unless otherwise noted. The wild-type allele in pAM5847 was amplified as a single fragment using primers pkaiA_F and kaiA_Down_R. The L156A allele in pAM5848 was amplified with the same primers using pAM5293 as template. The ΔN alleles were each assembled from two fragments: for pAM5832 (*kaiA*(aa 148-284)), fragments were amplified with pkaiA_F/kaiA_AA1R_1 and kaiA_148F/kaiA_Down_R; for pAM5833 (*kaiA*(aa 148-284, L156A)), fragments were amplified with pkaiA_F/kaiA_AA1R_1 from genomic DNA and kaiA_148F/kaiA_Down_R from pAM5293; and for pAM5834 (*kaiA*(aa 181-284)), fragments were amplified with pkaiA_F/kaiA_AA1R_2 and kaiA_181F/kaiA_Down_R.

A control plasmid carrying only the nourseothricin resistance marker targeting NS1, pAM5613, was derived from pAM5604 by SwaI digestion to release the ccdB toxic gene cassette, followed by re-ligation of the backbone. pAM5604 was assembled from three fragments by seamless cloning according to the CyanoVECTOR strategy: the NS1 recombination fragments with an

RK2 origin of mobilization from pCVD019 (NS1RK2BOM), the NAT7942 nourseothricin resistance marker from pCVD064, and the SwaI-flanked *ccdB* counterselection cassette from pCVD015.

Plasmids pAM5944–pAM5946 were constructed for ectopic expression of *kaiA* alleles from *Anabaena* and *S. elongatus* (used as a positive control) at NS1. The *kaiA* coding sequences and their respective promoter regions were PCR-amplified from genomic DNA and cloned directly by seamless assembly into pCV0052 digested with SwaI. The *Anabaena kaiA* allele with its native promoter in pAM5944 (P*kaiA*_An–*kaiA*_An) was amplified as a single fragment using primers pKaiAana_F and kaiAa_D433R. The *S. elongatus kaiA* allele with its native promoter in pAM5945 (P*kaiA*_Se–*kaiA*_Se) was using primers pkaiA_F and kaiA_Down_R. The chimeric construct in pAM5946 (P*kaiA*_Se–*kaiA*_An) was assembled from two fragments amplified independently - the *S. elongatus* promoter region with primers pKaiAse_F/pKaiAse_R and the *Anabaena kaiA* coding sequence with primers kaiAa_3F/kaiAa_D433R - and cloned simultaneously into pCV0052 by seamless assembly. A control plasmid, pAM5943 (S7942NS1_AACC1), carrying only the gentamycin resistance marker at NS1, was derived from pCV0052 by SwaI digestion to release the ccdB toxic gene cassette, followed by re-ligation of the backbone.

Plasmid pAM5900 (pUC19–*kaiA*Δ(296–623)::*aadA*) was used to knock out *kaiA* and was derived from pAM5831 (pUC19–*kaiA*Δ(296–623)::*aadA*-*sacB*) digested by AgeI and SacI, end-blunted, and re-ligation to remove the *sacB* counterselection marker. Plasmid pAM5831 was constructed by seamless assembly of three PCR fragments into pUC19. A DNA fragment of ~ 900 bp usptream of *kaiA* base 296 was amplified from *S. elongatus* genomic DNA using primers kaiA_U607F and kaiA_296R; the *aadA*-*sacB* cassette was amplified from pAM5001 (culture collection, R. Simkovsky) using primers aadA-pCVD2_R and ATP0121; and a ~1100 bp downstream DNA fragment starting at base 623 of *kaiA* was amplified from genomic DNA using primers kaiA_623F and kaiA_D1763R.

Plasmids pAM5908 and pAM5909 were used to replace the wild-type *kaiA* allele in its native locus with the *kaiA*(aa 148-284) and *kaiA*(aa 148-284, L156A) alleles, respectively. Both plasmids were constructed by seamless assembly following the CyanoVECTOR strategy (Taton et al., 2014) using three fragments: the RSF1010 backbone from pAM5406 digested with ZraI, the NAT7942 nourseothricin resistance marker from pCVD064 digested with EcoRV, and the P*kaiA*–*kaiA* inserts obtained by EcoRV digestion of pAM5832 (*kaiA*(aa 148-284)) and pAM5833 (*kaiA*(aa 148-284, L156A)), yielding pAM5908 (RSF1010BB–NAT7942–P*kaiA*-*kaiA*(aa 148-284)) and pAM5909 (RSF1010BB–NAT7942–P*kaiA*-*kaiA*(aa 148-284, L156A)).

Plasmids pAM5819 and pAM5820 were constructed to express crRNAs targeting the *kaiA* coding sequence for CRISPR-Cas12a editing. For each construct, pAMT004 was digested with AarI to release the lacZα fragment, and phosphorylated oligonucleotides were annealed and ligated into the linearized backbone. For pAM5819 (pAMT004-crRNA[KaiA350–369]), oligonucleotides 112_Fwd_SA and 112_Rev_SA were used, and for pAM5820 (pAMT004–crRNA[*kaiA*469–488]), oligonucleotides 156_1_Fwd_SA and 156_1_Rev_SA.

Plasmid pAMT004 (RSF1010BB–APH1–CPF1–CRISPR) was constructed by seamless assembly of three fragments following the CyanoVECTOR strategy [24]: the RSF1010 backbone from pAM5406 digested with ZraI, the kanamycin resistance marker APH1 from pCVD003 digested with EcoRV, and the Cas12a (Cpf1) expression cassette with its CRISPR array from pAMT001 digested with EcoRV.

Complete segregation was confirmed by PCR, and 3 to 4 independent clones were selected for downstream analyses.

Standard molecular biology techniques were used for plasmid construction and verification. PCRs were performed with a high-fidelity DNA polymerase, and plasmid DNA was isolated using commercial miniprep kits. DNA fragments were assembled using either restriction/ligation or seamless cloning methods. All plasmids were propagated in *E. coli* DH5α or TOP10 strains.

**Bioluminescence measurements**

*Synechococcus elongatus* strains expressing firefly luciferase (luc) under the control of the PkaiBC promoter were used as reporters of circadian rhythms. For pre-culture, cells were grown (inoculated at an $OD_{750}$ of 0.05) in 6-well plates under 12 h light:12 h dark cycles at 50 $\mu$mol photons $m^{-2}$ $s^{-1}$ for 3–4 days, reaching an $OD_{750}$ of ~0.4. Cultures were then adjusted to $OD_{750}$ = 0.2, and 20 $\mu$L of cell suspension was dispensed into 96-well plates containing 270 $\mu$L of BG-11 medium solidified with 1.5% (w/v) agar and supplemented with 10 $\mu$L of 5 mM D-luciferin per well [25].

Bioluminescence was recorded using one of two platforms. In the first setup, plates were loaded into a Tecan Spark plate reader equipped with a stacker microplate handling system fitted with LEDs and housed in a large temperature-controlled incubator. In the second setup, plates were incubated in a temperature-controlled plate incubator (MIO) fitted with LED illumination and transferred to a Tecan Spark plate reader by the robotic arm of a Tecan EVO 100 liquid handling platform. In both configurations, bioluminescence readings were acquired at 2 h intervals over several days.

#### Circadian bioluminescence analysis

*Model fitting.* Bioluminescence time-series from circadian reporter strains were analyzed in MATLAB R2023a (MathWorks).

Bioluminescence traces from several replicates and strains on multiple plates were aligned on a common time axis corresponding to the nominal sampling interval. Readings were sorted by time, and any empty entries (missing values for time measured on different plates) were replaced by linear interpolation along this shared axis, after which rows with remaining missing values were discarded to ensure complete time-series. Optionally, traces were linearly detrended by subtracting a best-fitting straight line from each time-series.

To determine period and phase, each time series within an analysis window was fitted to a chirped damped cosine model:

$$y(u) = A\,e^{-\lambda u} \cdot \cos\left(\omega_0 u + \frac{1}{2}\kappa u^2 + \phi_0\right) + B,$$

where $u = t - t_0$ is elapsed time since the start of the window, $A$ is the amplitude at $t_0$, $\lambda$ is the exponential damping rate ($\mathrm{h}^{-1}$), $\omega_0$ is the angular frequency at $t_0$ (rad $\mathrm{h}^{-1}$), $\kappa$ is a small chirp rate (rad $\mathrm{h}^{-2}$) describing gradual change in apparent period over time, $\phi_0$ is the initial phase (rad), and $B$ is the constant offset (mesor). When $\kappa = 0$, this reduces to the standard damped cosine [1]. Allowing $\kappa \neq 0$ lets the model capture slow period drift within a recording without changing how period and phase are defined and interpreted.

The instantaneous angular frequency of this model is

$$\omega(u) = \omega_0 + \kappa u,$$

and the instantaneous period at elapsed time $u$ is

$$T(u) = \frac{2\pi}{\omega_0 + \kappa u}.$$

Initial frequency estimates were obtained by applying a fast Fourier transform to mean-subtracted traces and selecting the dominant spectral peak within a circadian range (typically 18–30 h); the phase and magnitude of this peak provided initial values for oscillation phase and amplitude. Model parameters $(A, \omega_0, \phi_0, B, \lambda, \kappa)$ were then refined by nonlinear least squares with parameter bounds chosen to keep the instantaneous period within the specified range and to limit the amount of allowed period variation.

For each fit, we defined the primary period $T_{\text{mid}}$ as the instantaneous period at the temporal midpoint of the analysis window:

$$T_{\text{mid}} = T(u_{\text{mid}}) = \frac{2\pi}{\omega_0 + \kappa u_{\text{mid}}},$$

where $u_{\text{mid}}$ is the elapsed time at the midpoint of the window. In addition, a time-averaged period $T_{\text{mean}}$ over the window was computed analytically from the fitted $\omega_0$ and $\kappa$.

*Acrophase, phase statistics, and ensemble waveforms*. Acrophase (peak time) was defined relative to the experiment-specific reference time (typically the last dark-to-light transition before the cells were placed under constant light). For each fitted curve, the chirp model was evaluated on a dense grid (10,000 points) across the analysis window, and local maxima were detected numerically. Acrophase was taken as the time $t_{\text{peak}}$ of the first model peak at or after the reference time that lay within the analysis window; when no such peak was present, the peak closest to the window midpoint was used. Acrophase was reported in hours as:

$$\phi_{\text{adj}} = \text{mod}(t_{\text{peak}} - t_{\text{ref}}, T_{\text{mid}}),$$

where $t_{\text{peak}}$ is the time of the selected model peak, $t_{\text{ref}}$ is the reference time, and $T_{\text{mid}}$ is the period at the segment midpoint.

For each condition, fitted curves from all replicates were interpolated onto a common 100-point grid spanning the analysis window and averaged to obtain ensemble waveforms, with pointwise standard deviations quantifying across-replicate variability. Periods, acrophases, and damping rates across replicates were summarized as mean ± standard deviation. Mean acrophase and its dispersion were computed by mapping $\phi_{\text{adj}}$ into angles $2\pi\phi_{\text{adj}}/T_{\text{mid}}$, calculating the circular mean direction and resultant vector length, and converting back to hours, following standard circular statistics approaches.

*Temperature dependence and* $Q_{10}$. Temperature compensation was assessed from period estimates obtained at multiple constant temperatures. For each strain or condition, the mean period and associated variability were calculated at each temperature. A regression model (typically linear in temperature, with optional higher-order terms specified a priori) was fitted to period versus temperature, and the quality of fit was assessed by the coefficient of determination $R^2$.

The temperature coefficient $Q_{10}$ was then estimated in two ways. First, a global $Q_{10}$ was obtained from the regression by evaluating the fitted period at the lowest and highest experimental temperatures and applying

$$Q_{10} = \left(\frac{\tau(T_1)}{\tau(T_2)}\right)^{\frac{10}{T_2 - T_1}}.$$

Second, pairwise $Q_{10}$ values between adjacent temperatures were computed using the same expression applied to empirical mean periods. Uncertainties in $Q_{10}$ were estimated by propagating the regression parameter covariance for the regression-based estimate and by combining period standard deviations for the pairwise estimates.

*Phase-response curves (PRCs).* For each dark-pulse time, strain, and condition, the acrophases from individual replicates were aggregated, and circular mean acrophases were computed. Phase shifts were defined as the difference in mean acrophase between pulsed (4h dark pulse) and control groups plotted against the pulsed ZT to construct phase-response curves, with error bars reflecting circular dispersion.

Custom MATLAB code implementing all steps (preprocessing, chirp model fitting, acrophase calculation, $Q_{10}$ estimation, and PRC construction) is available at https://github.com/aataton/circadian_bioluminescence_matlab. Development of the code for the analyses and initial corresponding Methods section was assisted by the AI language model Claude Sonnet 4.6 (Anthropic; accessed via Perplexity), with all final implementations and wording verified by the authors.

**Serial dilution spot plate assays to determine growth and LD sensitivity under different conditions**

Cells were inoculated into 8 mL BG11 medium supplemented with the appropriate antibiotics at an initial $OD_{750}$ of 0.05. Cultures were grown in 6-well plates at 30 °C and 40 $\mu$mol photons $m^{-2}$ $s^{-1}$ with shaking under continuous light for 3 days, followed by two 12-h light:12-h dark cycles, until cultures reached an $OD_{750}$ of approximately 0.5. Then cell concentration was determined by flow cytometry, and cultures were adjusted to 2,000 cells $\mu L^{-1}$. Flow cytometer (Beckman Coulter, CytoFLEX platform B2-R2-V2) gains for FSC and SSC were set to 1,000, with a primary FSC threshold of 100,000 and a secondary SSC threshold of 20,000; *S. elongatus* cells were gated by autofluorescence of photosynthetic pigments detected in the APC channel (Ex

638 nm/Em 660/20 nm, gain 600). Cultures were serially diluted 4-folds, and 4-$\mu$L aliquots of each dilution were spotted onto BG11 agar plates. Plates were incubated for 5 to 7 days at 25, 30, 35, or 40 °C under light intensities of 75, 150, or 300 $\mu$mol photons $m^{-2}$ $s^{-1}$. Pictures of the plates were taken, and colonies within each spot with isolated colonies, were quantified using a custom Fiji/ImageJ macro (https://github.com/aatatton/spot_assay_colony_counter_Fiji).

**Use of artificial intelligence**

ChatGPT (OpenAI), Claude (Anthropic), and Perplexity AI were used to assist with the development, modification, and troubleshooting of Python scripts used for data plotting and visualization in the Spyder (Scientific Python Development Environment) environment, as well as for grammar and language editing of the manuscript. All AI-assisted code and text were reviewed, edited, and verified by the authors. The authors remain responsible for the accuracy, integrity, and interpretation of the data and results and for the final content of the manuscript. All data analysis, interpretation of results, and scientific conclusions were performed and determined by the authors.

#### Supplementary Figures

(a)

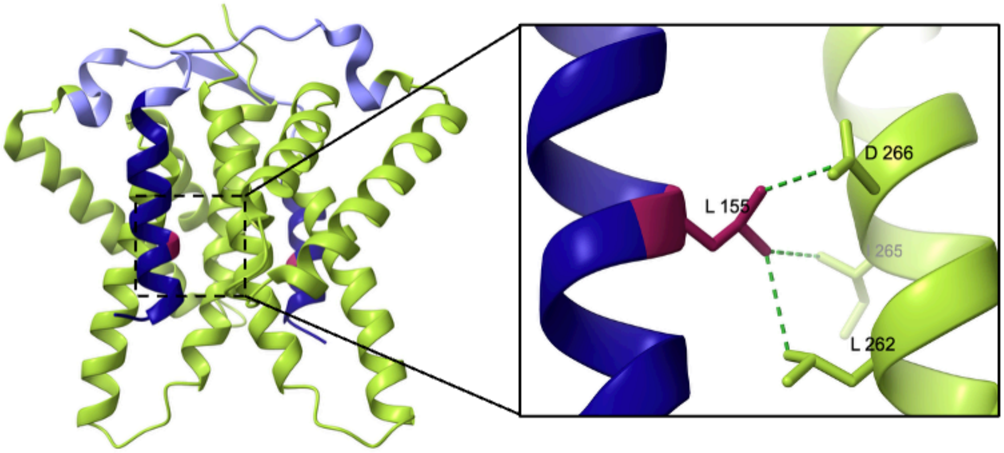

(b)

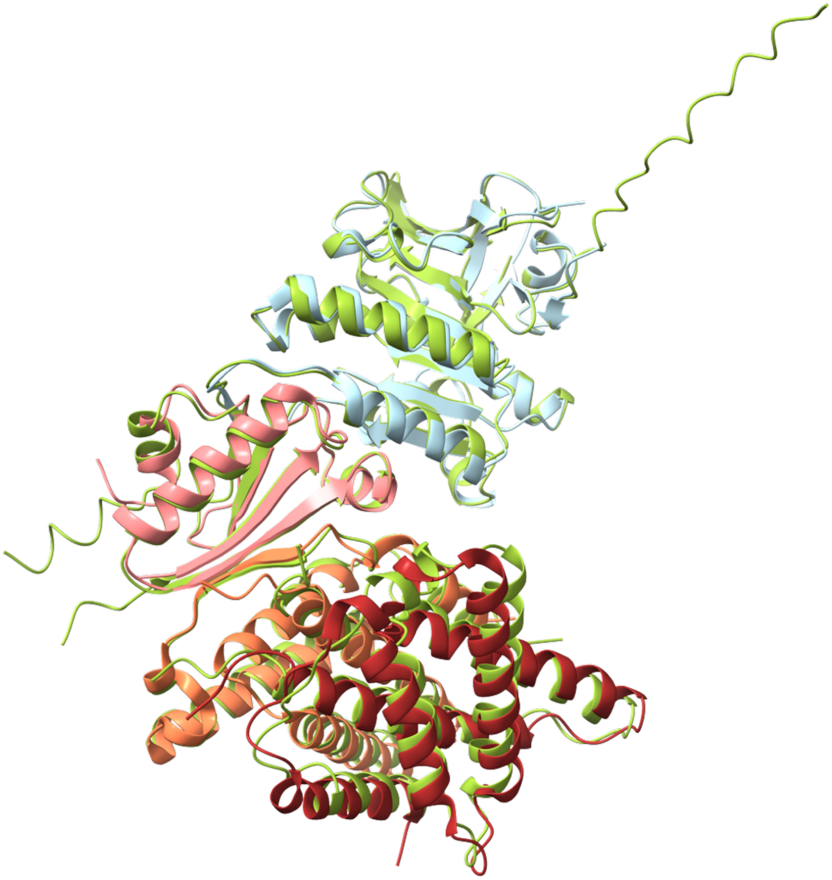

**Supplementary Figure 1. Structural analysis of the closed KaiA conformations.**

(a) L155 residue forms multiple hydrophobic contacts with residues in C-KaiA. The inactive conformation of *T. elongatus* KaiA (PDB ID 5JWR.E, F) shows carbon-carbon contacts between α5 residue L155 and L262, I265, and D266. (b) Structural alignment of ΔN-KaiA (orange, maroon), fold-switched KaiB (pink), KaiC CI (light blue) complex (PDB ID 5JWR) with AlphaFold-predicted *S. elongatus* complex (green). The alignment has a Root Mean Square Deviation (RMSD) across 227 pairs of 0.72.

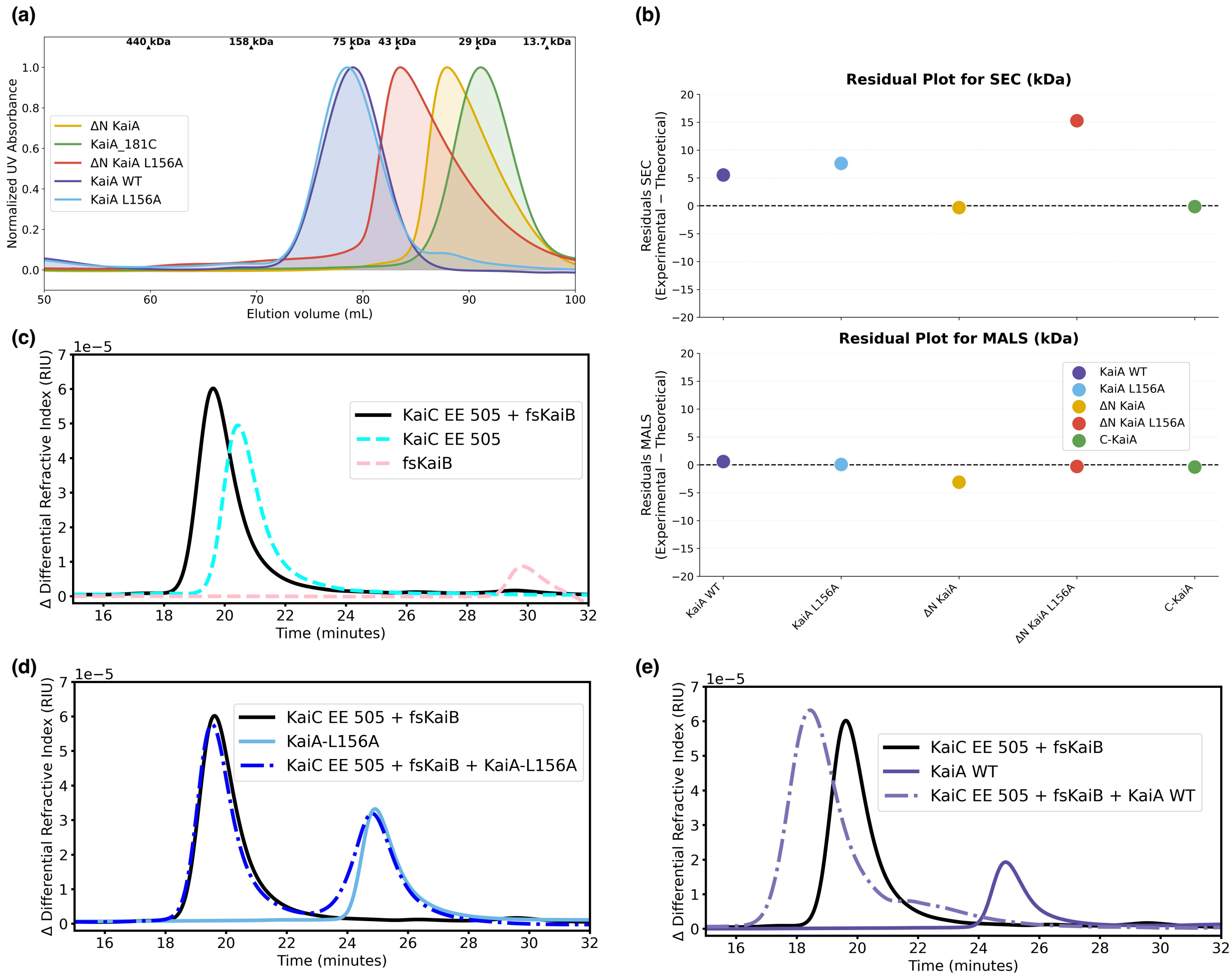

**Supplementary Figure 2. SEC-MALS characterization of KaiA mutants.**

**(a)** Normalized UV absorbance profiles of KaiA mutants using a Superdex 200 16/600 column. (**b)** Residuals of experimental versus theoretical molecular weights for KaiA variants measured by SEC (top) and MALS (bottom). For each protein, the residual is defined as experimental minus theoretical molecular weight (kDa). **(c)** SEC-MALS of KaiC S431E T432E 1-505 (35 μM) and fold-switched KaiB (35 μM) complex using a Superdex 200 10/300 analytical column. **(d)** SEC-MALS of KaiA L156A (35 μM), KaiC S431E T432E 1-505 (35 μM), and fold-switched KaiB (35 μM) complex. KaiA L156A could not be sequestered by the KaiB-KaiC complex. **(e)** SEC-

MALS of KaiA WT (35 µM), KaiC S431E T432E 1-505 (35 µM), and the fold-switched KaiB (35 µM) complex.

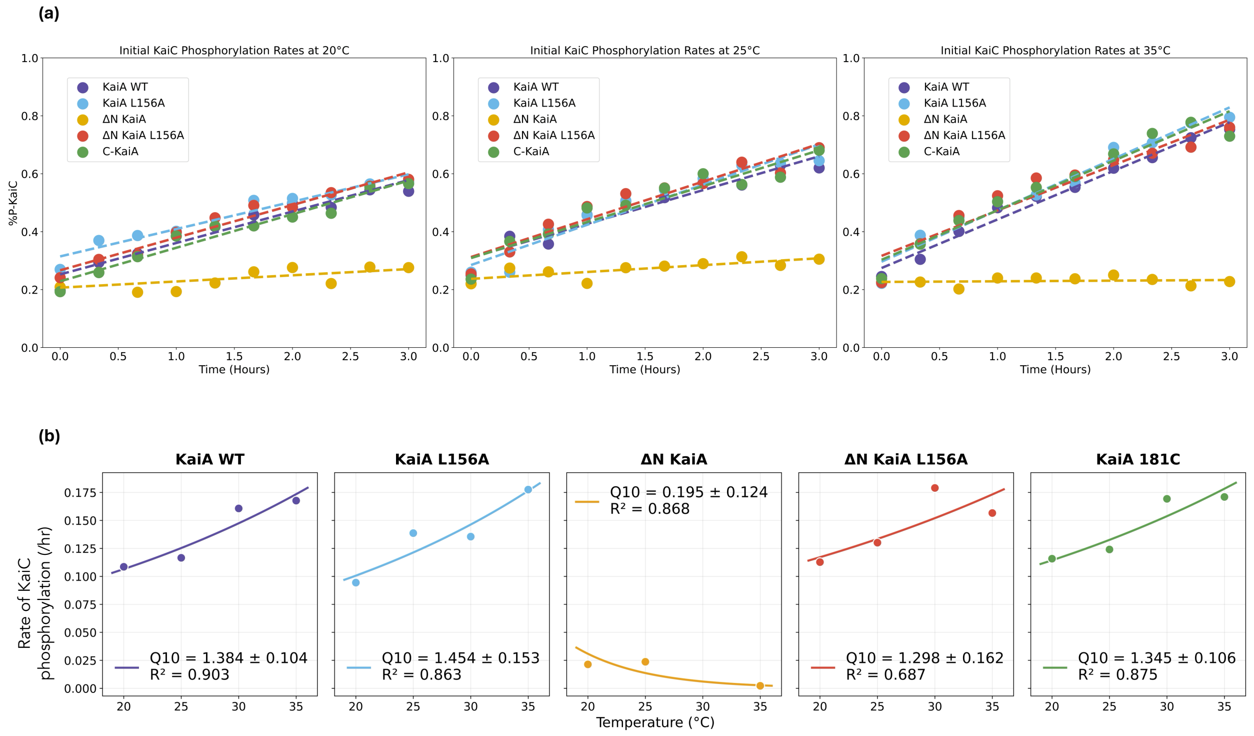

**Supplementary Figure 3. KaiA-dependent KaiC initial autophosphorylation kinetics.**

KaiC autophosphorylation rate was measured by gel assays in the presence of wild-type or mutant KaiA constructs: KaiA WT, KaiA L156A, ΔN KaiA, ΔN KaiA L156A, and KaiA 181C. **(a)** Level of phosphorylated KaiC (% P-KaiC) as a function of time across different temperatures. **(b)** Q10 plots of each KaiA mutant-dependent KaiC autophosphorylation rate. For each construct, phosphorylation rate is plotted against temperature on a semi-log scale. Solid lines show the log-linear regression fit, $\ln(\text{rate}) = (\ln(Q10)/10)\cdot T + b$, from which the temperature coefficient $Q10 = e^{(10*\text{slope})}$ was calculated for each construct. Since the KaiC phosphorylation rate itself is weak, the Q10 of the ΔN KaiA mutant is unreliable.

**(a)**

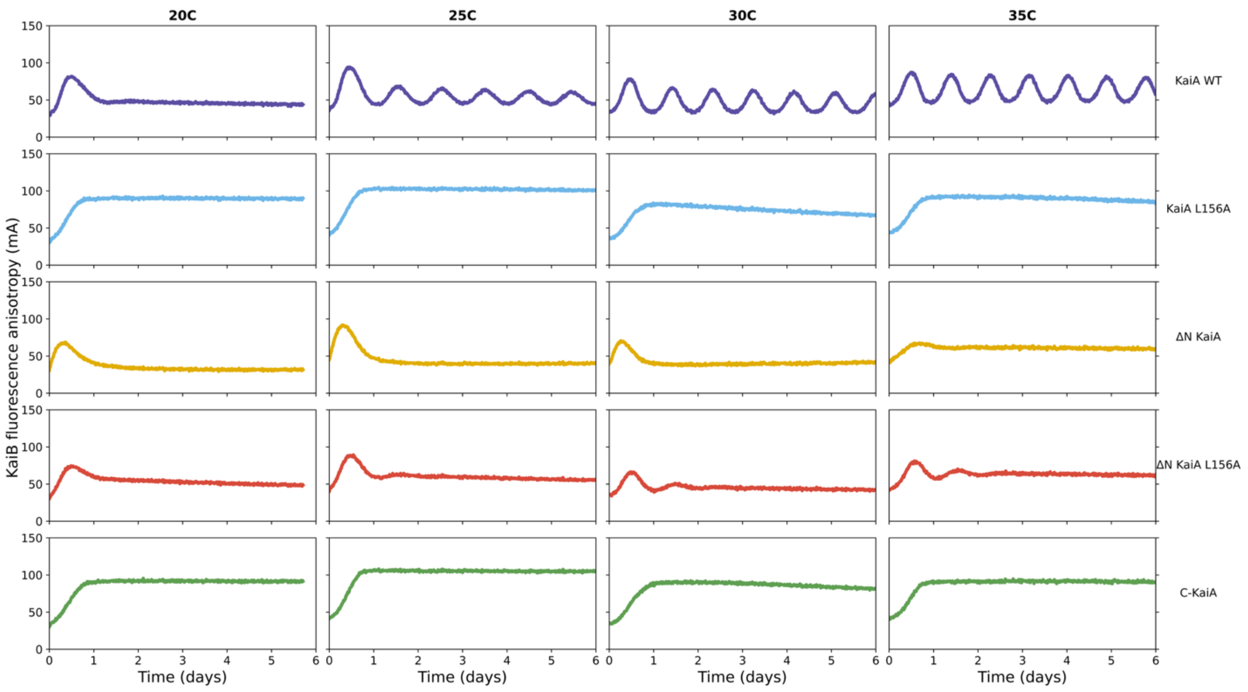

**(b)**

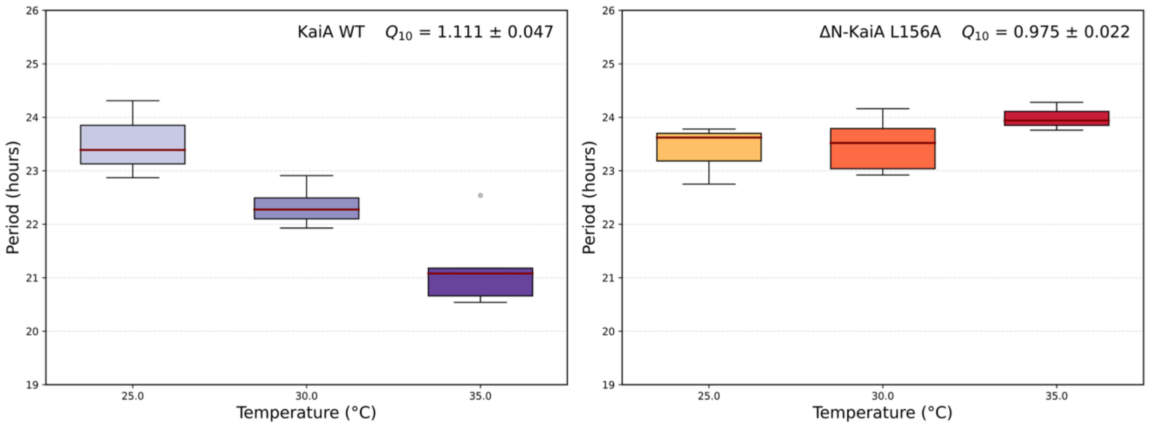

**Supplementary Figure 4. In vitro reconstituted KaiABC fluorescence anisotropy assays.**

**(a)** IVO with different KaiA mutants across temperatures with 50 nM fluorescently labeled KaiB as a probe. Each reaction includes KaiC WT (3.5 µM), KaiB WT (3.45 µM), and KaiA mutant (1.2 µM). **(b)** Q10 plots of Period against Temperature for KaiA WT (left) and ΔN-KaiA L156A (right). Box plots show in vitro oscillation periods (h) measured per temperature. Boxes span the interquartile range (IQR), dark red lines mark the median, whiskers extend to the most extreme values within 1.5 × IQR, and gray circles show values outside this range. Box shading indicates increasing temperature. $Q_{10}$ was calculated as $Q_{10} = (f_2/f_1)$^(10/($T_2$–$T_1$)), where f = 1/period, for every pairwise combination of individual replicates at the lowest ([$T_1$] °C) and highest ([$T_n$] °C) temperatures. The value shown is the mean ± SD of these pairwise estimates.

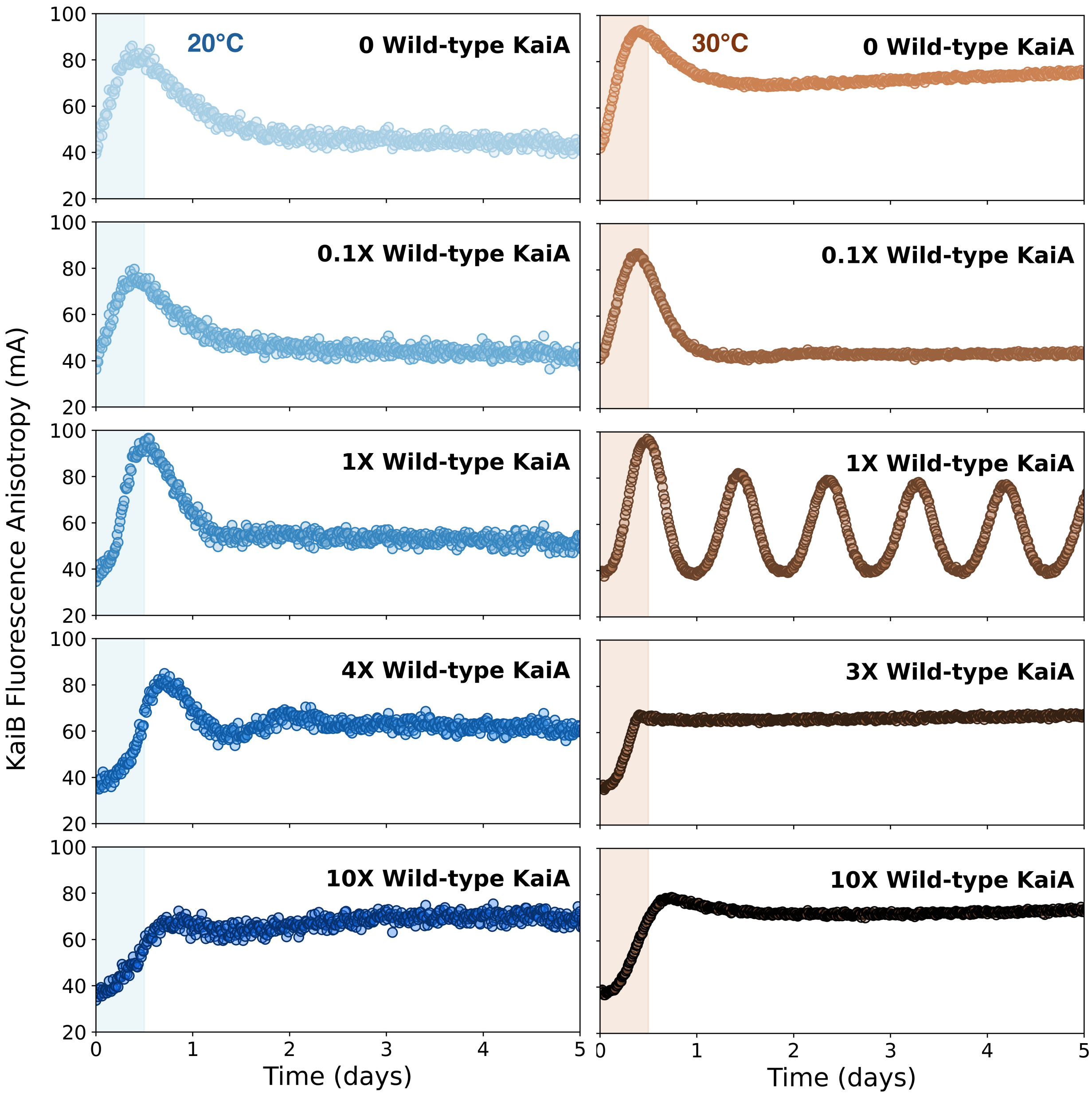

**Supplementary Figure 5. In vitro oscillations with varying KaiA WT concentrations.**

(Left) 20°C and (Right) 30°C. The initial 12 hours are highlighted in the shaded region. Increasing KaiA WT concentrations cause a stronger antagonistic effect on initial KaiB-KaiC interaction (KaiB anisotropy level) and later sustain higher KaiB-KaiC binding (1X is 1.2 μM of

KaiA). Higher KaiA WT concentrations mimic the constitutively open KaiA mutants like KaiA L156A and C-KaiA (Supplementary Fig. 4).

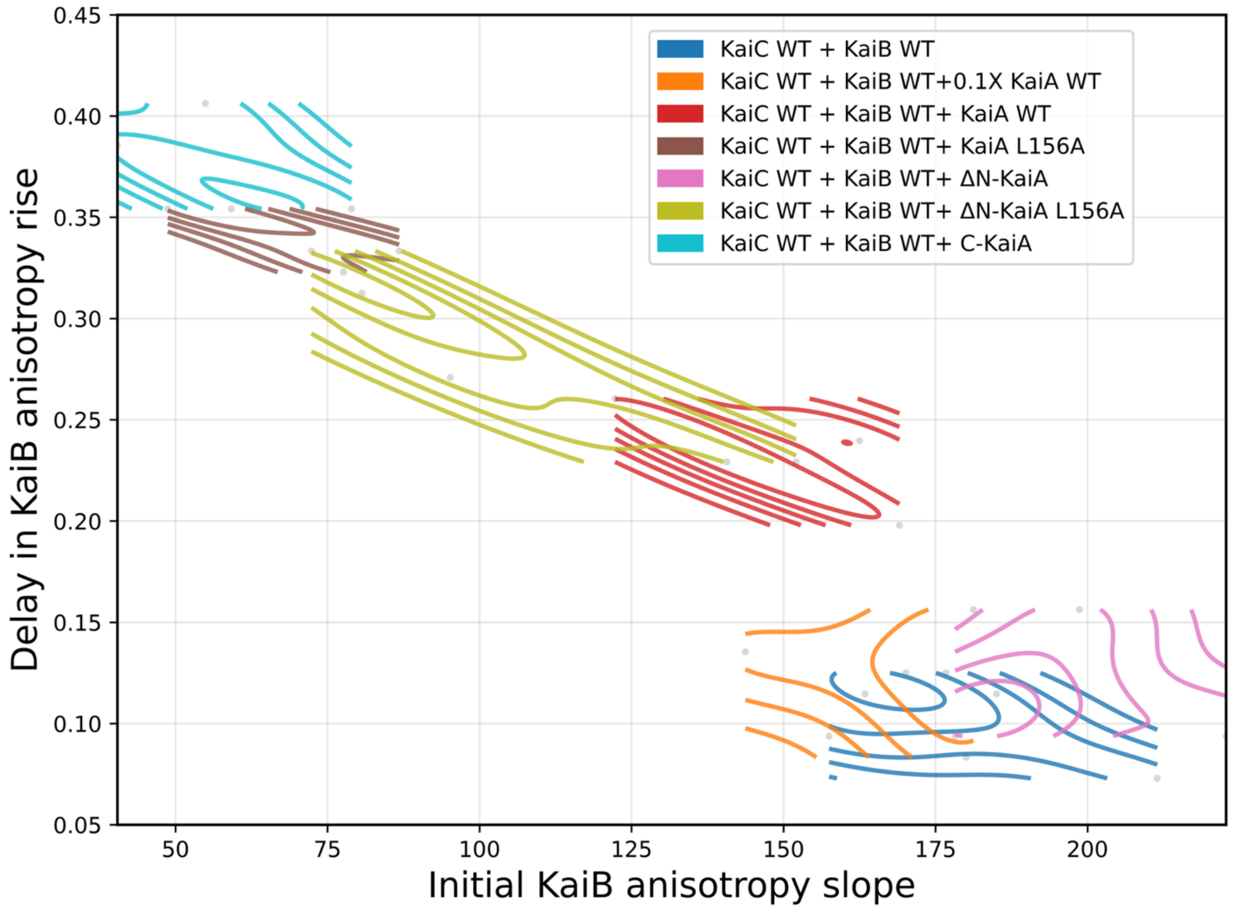

**Supplementary Figure 6. Density distribution of KaiB anisotropy kinetics with KaiA antagonists.**

Time delay induced for KaiB-KaiC binding (KaiB anisotropy increase) plotted against the slope of Initial KaiB anisotropy rise in the presence of each KaiA mutant across temperatures. Contour lines represent Gaussian KDE isodensity levels for each KaiA mutant, changing from 20°C to 35°C (color-coded per legend).

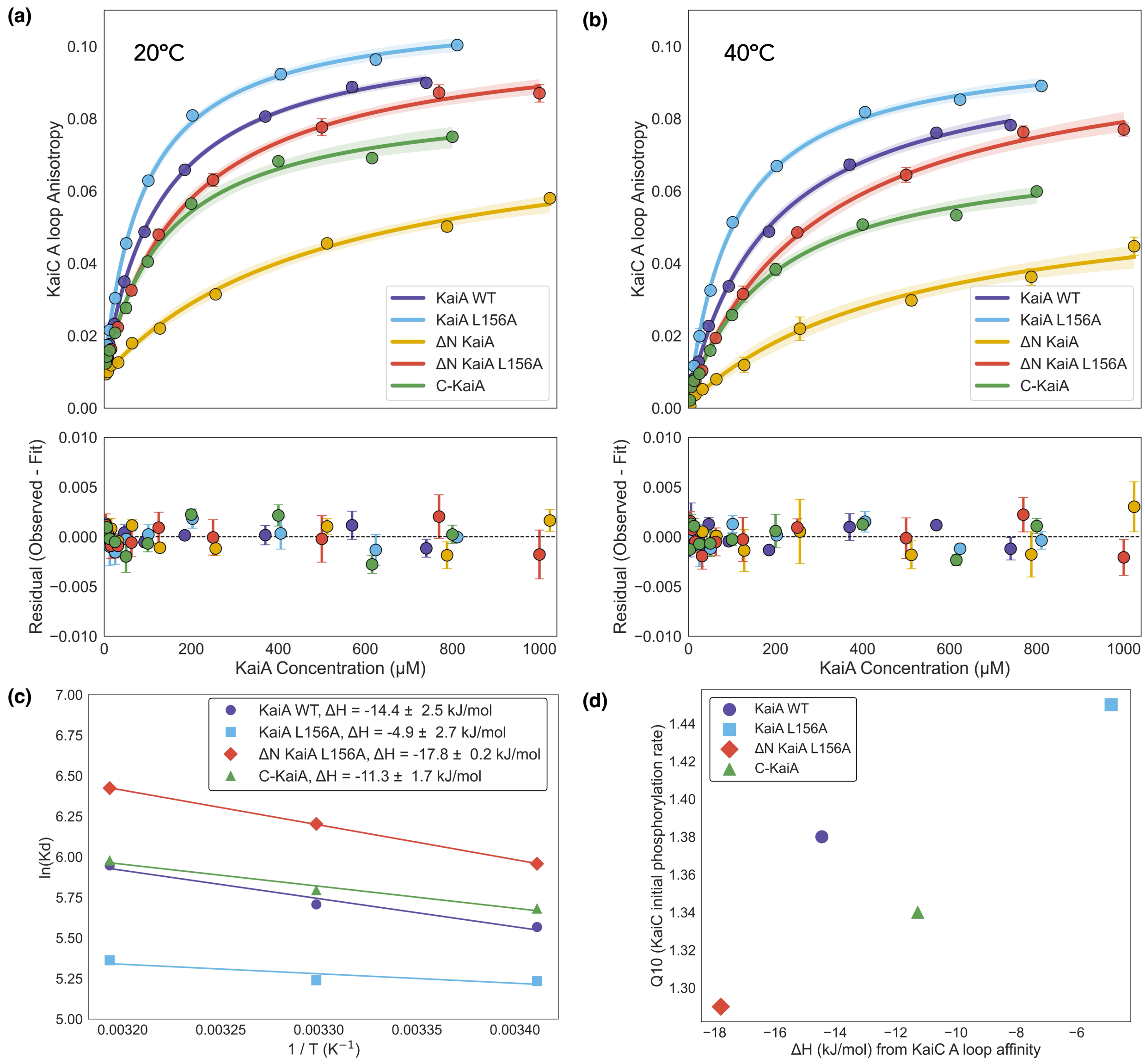

**Supplementary Figure 7. Fluorescence anisotropy profiles of KaiC A-loop peptides as a function of KaiA mutants.**

**(a) 20 °C, (b) 40 °C.** KaiA–KaiC A-loop binding fits with confidence bands and residuals. Fluorescence anisotropy of an A-loop peptide (0.05 $\mu$M) was measured as a function of KaiA concentration for KaiA WT, KaiA L156A, ΔN KaiA, ΔN KaiA L156A, and C-KaiA constructs at 20 °C, and data were fit to a 1:1 binding model that explicitly accounts for ligand depletion. Top panel: mean anisotropy values (symbols) ± SEM from up to three anisotropy replicates are shown together with best-fit binding curves (solid lines) and 95% confidence bands (shaded regions) obtained from the parameter covariance matrix and a t-based critical value. Bottom panel, residuals (observed – fit anisotropy) for each construct are plotted with the same color scheme and ± SEM error bars, with a dashed horizontal line indicating zero residual to visualize systematic deviations from the model. KaiA concentration (μM) is shown on the shared x-axis. **(c)** van 't Hoff plot of KaiA mutants and KaiC binding showing $\ln(1/K_D)$versus $1/T$, with each point representing the fitted equilibrium dissociation constant at a given temperature. The linear fit is used to extract $\Delta H^{\circ}$from the slope and $\Delta S^{\circ}$from the intercept. **(d)** Comparison of Q10 obtained from initial KaiC autophosphorylation and $\Delta H^{\circ}$obtained from the KaiC A loop anisotropy assay for each KaiA mutant.

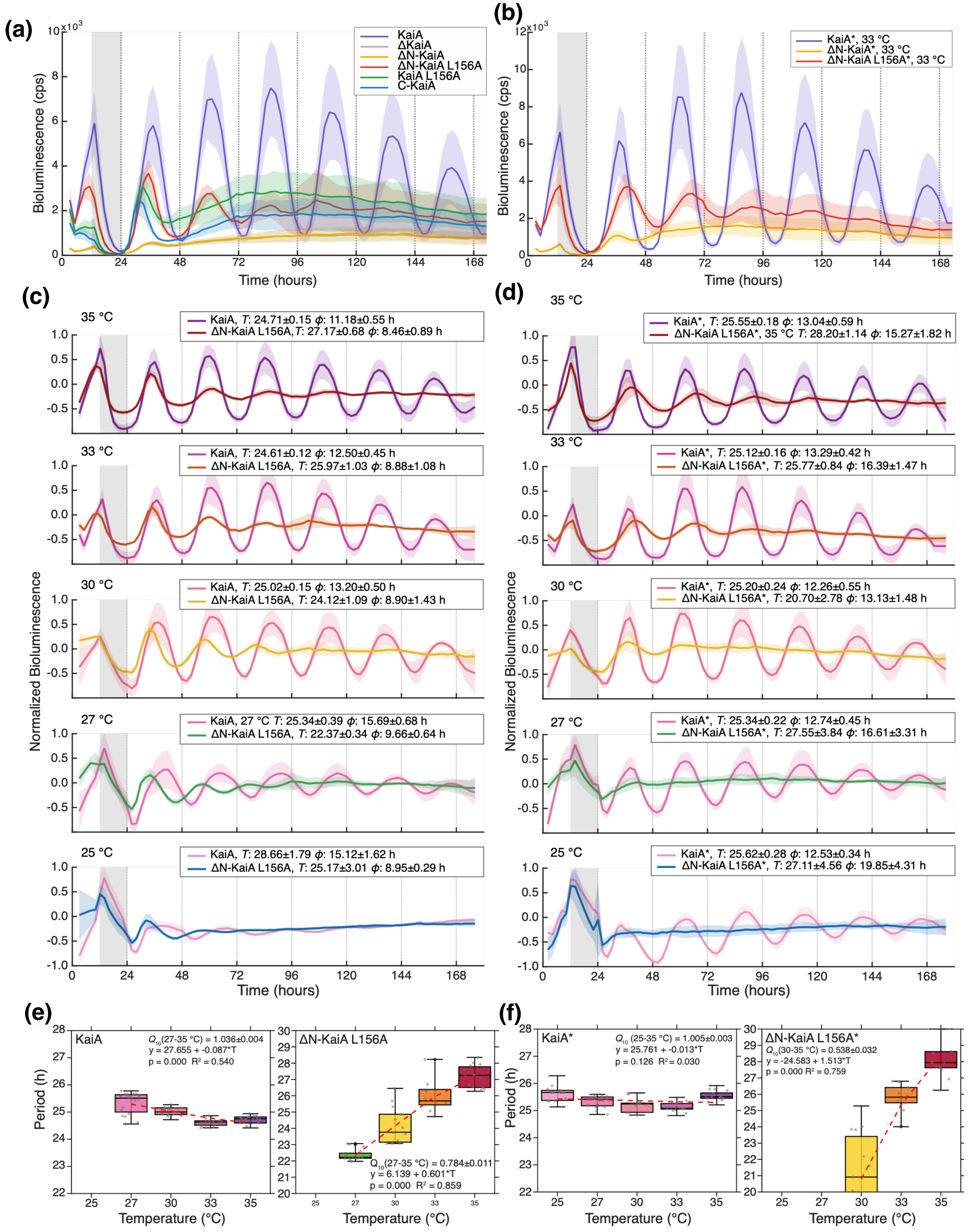
(a)
KaiA
ΔKaiA
ΔN-KaiA
ΔN-KaiA L156A
KaiA L156A
C-KaiA
Bioluminescence (cps)
Time (hours)
(b)
KaiA*, 33 °C
ΔN-KaiA*, 33 °C
ΔN-KaiA L156A*, 33 °C
Bioluminescence (cps)
Time (hours)
(c)
35 °C
KaiA, T: 24.71±0.15 ϕ: 11.18±0.55 h
ΔN-KaiA L156A, T: 27.17±0.68 ϕ: 8.46±0.89 h
33 °C
KaiA, T: 24.61±0.12 ϕ: 12.50±0.45 h
ΔN-KaiA L156A, T: 25.97±1.03 ϕ: 8.88±1.08 h
30 °C
KaiA, T: 25.02±0.15 ϕ: 13.20±0.50 h
ΔN-KaiA L156A, T: 24.12±1.09 ϕ: 8.90±1.43 h
27 °C
KaiA, 27 °C T: 25.34±0.39 ϕ: 15.69±0.68 h
ΔN-KaiA L156A, T: 22.37±0.34 ϕ: 9.66±0.64 h
25 °C
KaiA, T: 28.66±1.79 ϕ: 15.12±1.62 h
ΔN-KaiA L156A, T: 25.17±3.01 ϕ: 8.95±0.29 h
Normalized Bioluminescence
Time (hours)
(d)
35 °C
KaiA*, T: 25.55±0.18 ϕ: 13.04±0.59 h
ΔN-KaiA L156A*, 35 °C T: 28.20±1.14 ϕ: 15.27±1.82 h
33 °C
KaiA*, T: 25.12±0.16 ϕ: 13.29±0.42 h
ΔN-KaiA L156A*, T: 25.77±0.84 ϕ: 16.39±1.47 h
30 °C
KaiA*, T: 25.20±0.24 ϕ: 12.26±0.55 h
ΔN-KaiA L156A*, T: 20.70±2.78 ϕ: 13.13±1.48 h
27 °C
KaiA*, T: 25.34±0.22 ϕ: 12.74±0.45 h
ΔN-KaiA L156A*, T: 27.55±3.84 ϕ: 16.61±3.31 h
25 °C
KaiA*, T: 25.62±0.28 ϕ: 12.53±0.34 h
ΔN-KaiA L156A*, T: 27.11±4.56 ϕ: 19.85±4.31 h
Normalized Bioluminescence
Time (hours)
(e)
KaiA
Q10(27-35 °C) = 1.036±0.004
y = 27.655 + -0.087*T
p = 0.000 R² = 0.540
Period (h)
Temperature (°C)
ΔN-KaiA L156A
Q10(27-35 °C) = 0.784±0.011
y = 6.139 + 0.601*T
p = 0.000 R² = 0.859
Temperature (°C)
(f)
KaiA*
Q10 (25-35 °C) = 1.005±0.003
y = 25.761 + -0.013*T
p = 0.126 R² = 0.030
Period (h)
Temperature (°C)
ΔN-KaiA L156A*
Q10(30-35 °C) = 0.538±0.032
y = -24.583 + 1.513*T
p = 0.000 R² = 0.759
Temperature (°C)

**Supplementary Figure 8. Temperature dependence of circadian rhythms in S. elongatus strains expressing ΔN-KaiA L156A vs. KaiA WT.**

**(a)** $P_{kaiBC}$-*luc* bioluminescence over a 7-day time course at 33 °C in Δ*kaiA* strains at 33 °C ectopically expressing KaiA WT (*T*: 24.56±0.13 *ϕ*: 12.59±0.43 h), ΔN-KaiA, ΔN-KaiA L156A (*T*: 25.81±1.25 *ϕ*: 9.09±1.14 h), KaiA L156A, and C-KaiA. **(b)** $P_{kaiBC}$-*luc* bioluminescence at 33 °C in *S. elongatus* strains carrying *kaiA* WT and CRISPR-edited *kaiA*(aa 148-284), and *kaiA*(aa 148-284, L156A) alleles, denoted as KaiA WT* (*T*: 25.40±0.98 *ϕ*: 16.62±1.66 h), ΔN-KaiA*, and ΔN-KaiA L156A* (*T*: 25.40±0.98 *ϕ*: 16.62±1.66 h). **(c)** Normalized bioluminescence (–1 to +1) across a 25-35°C temperature range (each row corresponds to a different temperature: 35, 33, 30, 27, and 25 °C, from top to bottom) for Δ*kaiA* strains expressing KaiA WT and ΔN-KaiA L156A. **(d)** Normalized bioluminescence across the temperature range in KaiA WT* and ΔN-KaiA L156A* *S. elongatus* strains. **(e)** Period and Q10 analyses of KaiA WT and ΔN-KaiA L156A strains across the temperature range. (**f)** Period and Q10 analyses of KaiA WT* and ΔN-KaiA L156A* *S. elongatus* strains across 25–35 °C. Period (*T*) and acrophase (*ϕ*) were estimated by fitting a chirped damped cosine model to the detrended bioluminescence time series; values are shown as mean ± SD (n>9, incl. 3-4 individual clones). The temperature dependence of the period was assessed by linear regression (dashed red line), with the Q10 coefficient, regression equation, slope p-value, and $R^2$ displayed within each panel. Grey shaded areas indicate the period during which the cells were incubated in darkness, corresponding to the night of light/dark entrainment cycles, after which measurements were conducted under constant illumination. Prior to being distributed in 96-well plates, the cells were entrained for three 12 h light: 12 h dark cycles as liquid cultures in 6 well-plates at 50 $\mu$mol photon $m^{-2}s^{-1}$.

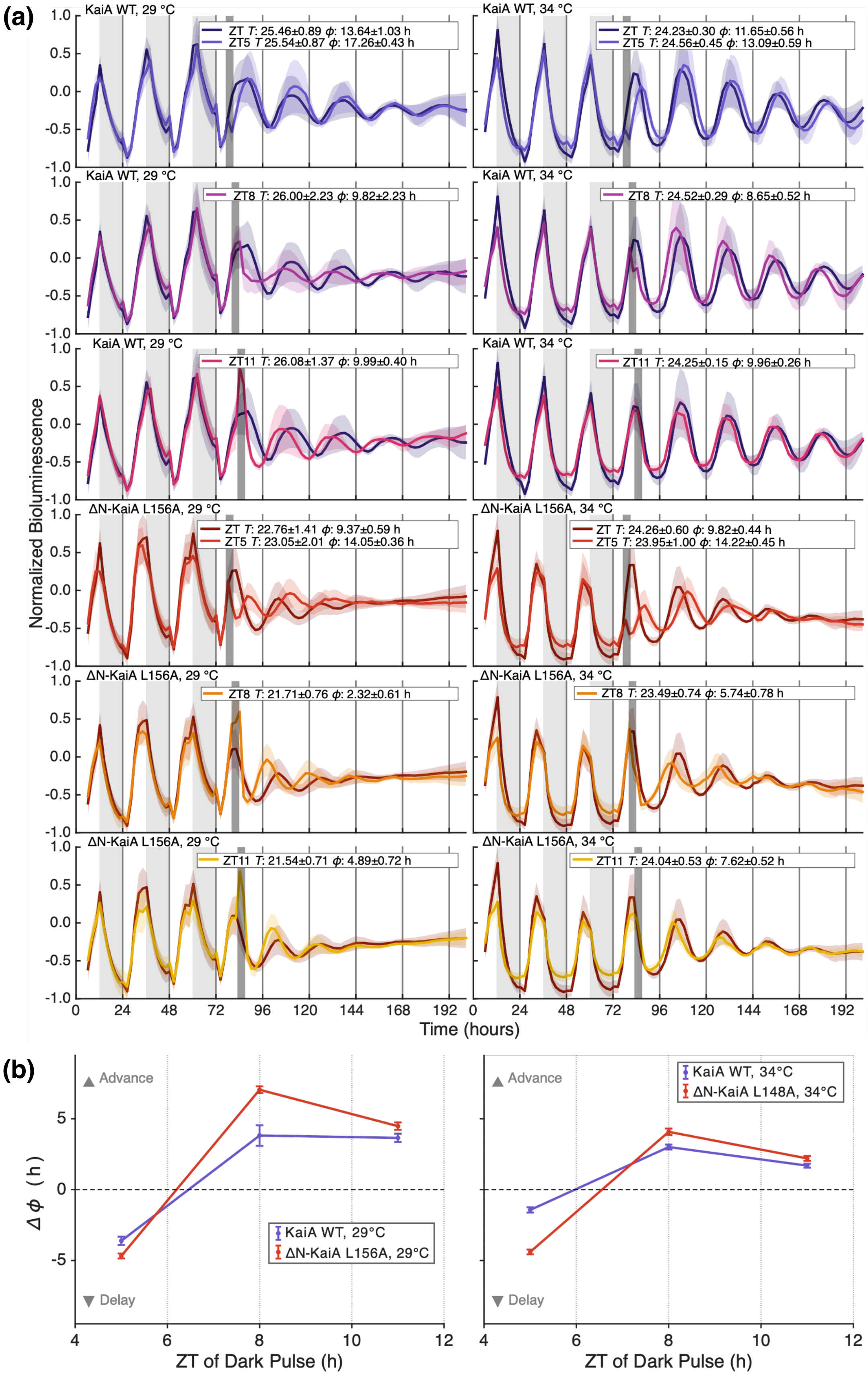

(a)
KaiA WT, 29 °C
ZT T: 25.46±0.89 ϕ: 13.64±1.03 h
ZT5 T 25.54±0.87 ϕ: 17.26±0.43 h
KaiA WT, 34 °C
ZT T: 24.23±0.30 ϕ: 11.65±0.56 h
ZT5 T: 24.56±0.45 ϕ: 13.09±0.59 h
ZT8 T: 26.00±2.23 ϕ: 9.82±2.23 h
ZT8 T: 24.52±0.29 ϕ: 8.65±0.52 h
ZT11 T: 26.08±1.37 ϕ: 9.99±0.40 h
ZT11 T: 24.25±0.15 ϕ: 9.96±0.26 h
ΔN-KaiA L156A, 29 °C
ZT T: 22.76±1.41 ϕ: 9.37±0.59 h
ZT5 T: 23.05±2.01 ϕ: 14.05±0.36 h
ΔN-KaiA L156A, 34 °C
ZT T: 24.26±0.60 ϕ: 9.82±0.44 h
ZT5 T: 23.95±1.00 ϕ: 14.22±0.45 h
ZT8 T: 21.71±0.76 ϕ: 2.32±0.61 h
ZT8 T: 23.49±0.74 ϕ: 5.74±0.78 h
ZT11 T: 21.54±0.71 ϕ: 4.89±0.72 h
ZT11 T: 24.04±0.53 ϕ: 7.62±0.52 h
Normalized Bioluminescence
Time (hours)
(b)
Advance
Delay
Δϕ (h)
KaiA WT, 29°C
ΔN-KaiA L156A, 29°C
KaiA WT, 34°C
ΔN-KaiA L148A, 34°C
ZT of Dark Pulse (h)

**Supplementary Figure 9. S. elongatus strains expressing the ΔN-KaiA L156A allele resets circadian phase similarly to WT.**

(**a)** $P_{kaiBC}$ bioluminescence over an 8-day time-course at 29 °C and 34 °C in Δ*kaiA* strains expressing ectopically KaiA WT or ΔN-KaiA L156A. Period (*T*) and acrophase (*ϕ*) were estimated by fitting a chirped damped cosine model to the detrended bioluminescence time series; values are shown as mean ± SD (n>9, incl. 3-4 individual clones). Cultures were maintained under 12 h light:12 h dark conditions during the first 3 days, then maintained under constant illumination and given a 4-hour dark pulse on the fourth day at ZT 5 h, 8 h, or 11 h. Grey shaded areas indicate the period during which the strains were incubated in darkness, corresponding to the night of light/dark entrainment cycles. Dark grey shaded areas correspond to the 4 h dark pulses. (**b)** Phase response curves (PRCs) at 29 °C and 34 °C for the Δ*kaiA* strains expressing ectopically WT KaiA and ΔN-KaiA L156A. Phase shifts (Δ *ϕ*, hours) in response to the 4-hour dark pulses were calculated. Positive values indicate phase advances; negative values indicate phase delays.

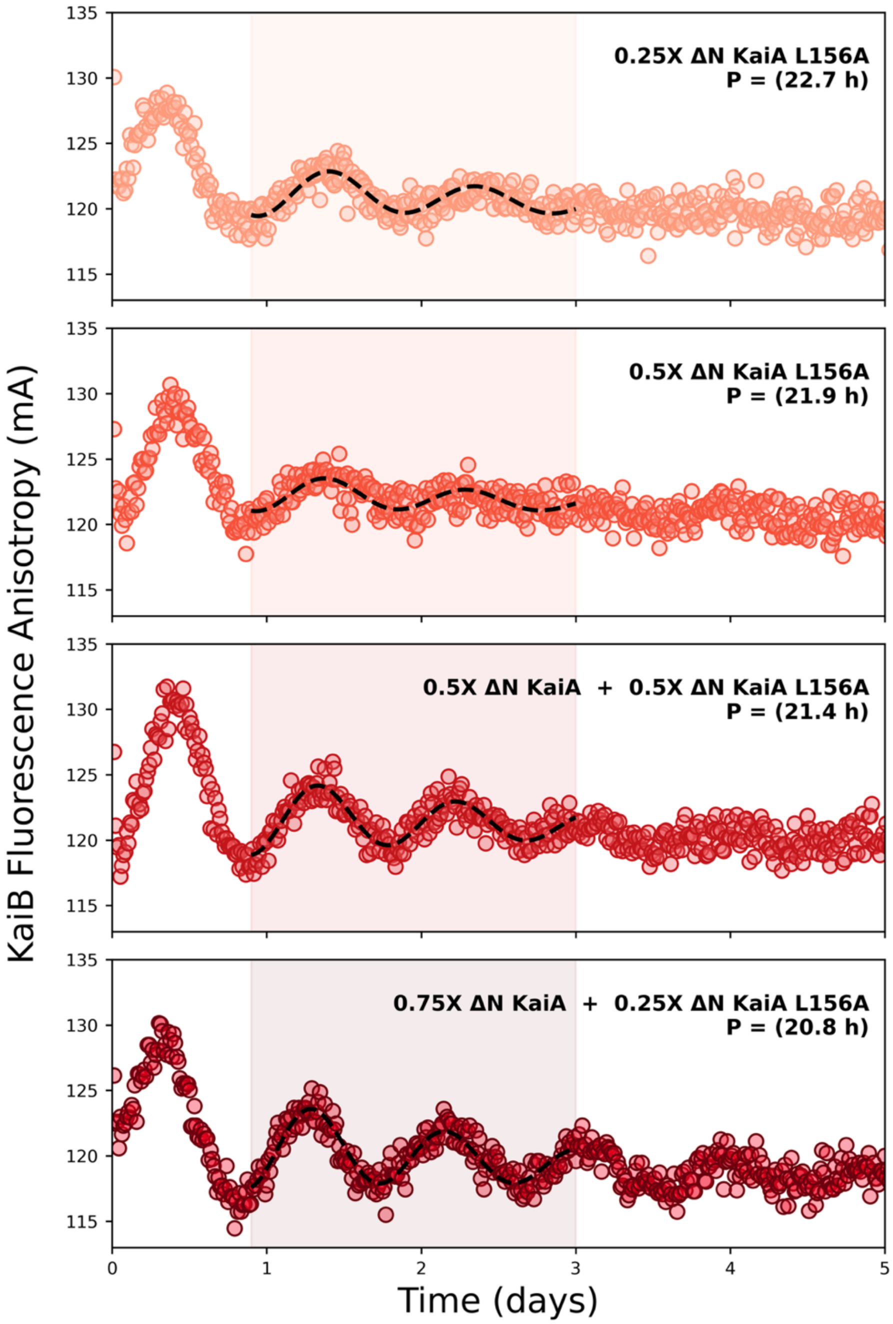

**Supplementary Figure 10. In vitro oscillations of ΔN-KaiA L156A.**

Titrating ΔN-KaiA with ΔN-KaiA L156A in different ratios gives more robust circadian rhythms with a shorter period. (1X is 1.2 μM of KaiA). The shaded region indicates the time range for period estimation.

(a)

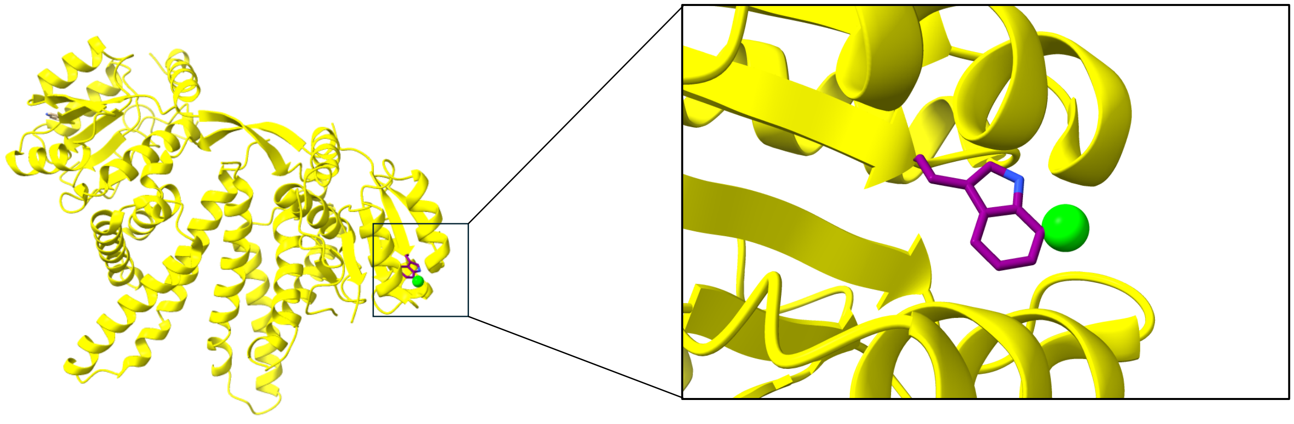

(b)

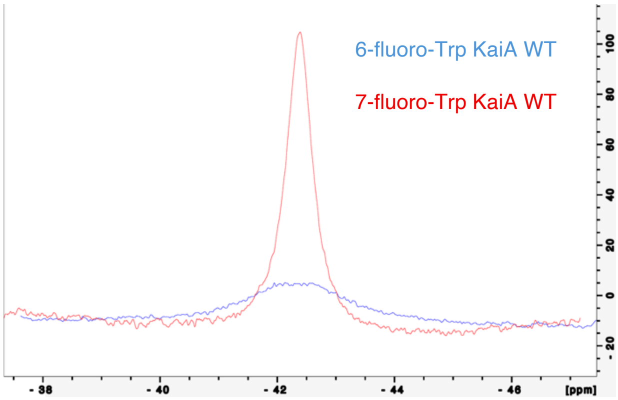

(c)

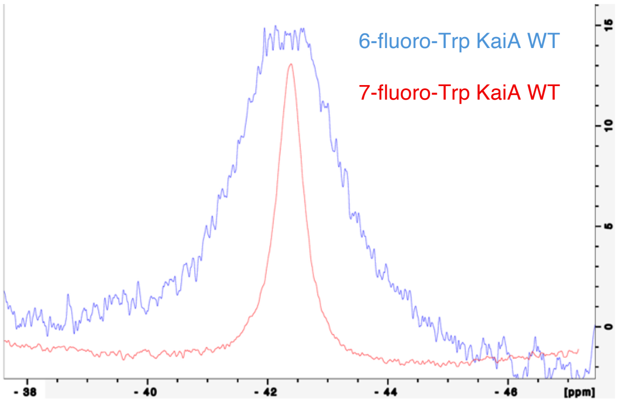

**Supplementary Figure 11. 19F NMR characterization of KaiA mutants.**

**(a)** $^{19}$F Tryptophan incorporation in KaiA (from *S. elongatus;* PDB ID 1R8J). Inlet box showing the modification of W10 as a 7-fluoro-tryptophan. **(b)** $^{19}$F NMR profile of KaiA WT with modification of W10 as a 6-fluoro-tryptophan aligned with 7-fluoro-tryptophan, same scale. **(c)** 6-fluoro-tryptophan KaiA WT aligned with 7-fluoro-tryptophan KaiA WT scaled down to 1/8$^{th}$. For both (b) and (c), the 7-fluoro-tryptophan spectrum was shifted 8.3718 ppm to match a 6-fluoro-tryptophan x-axis.

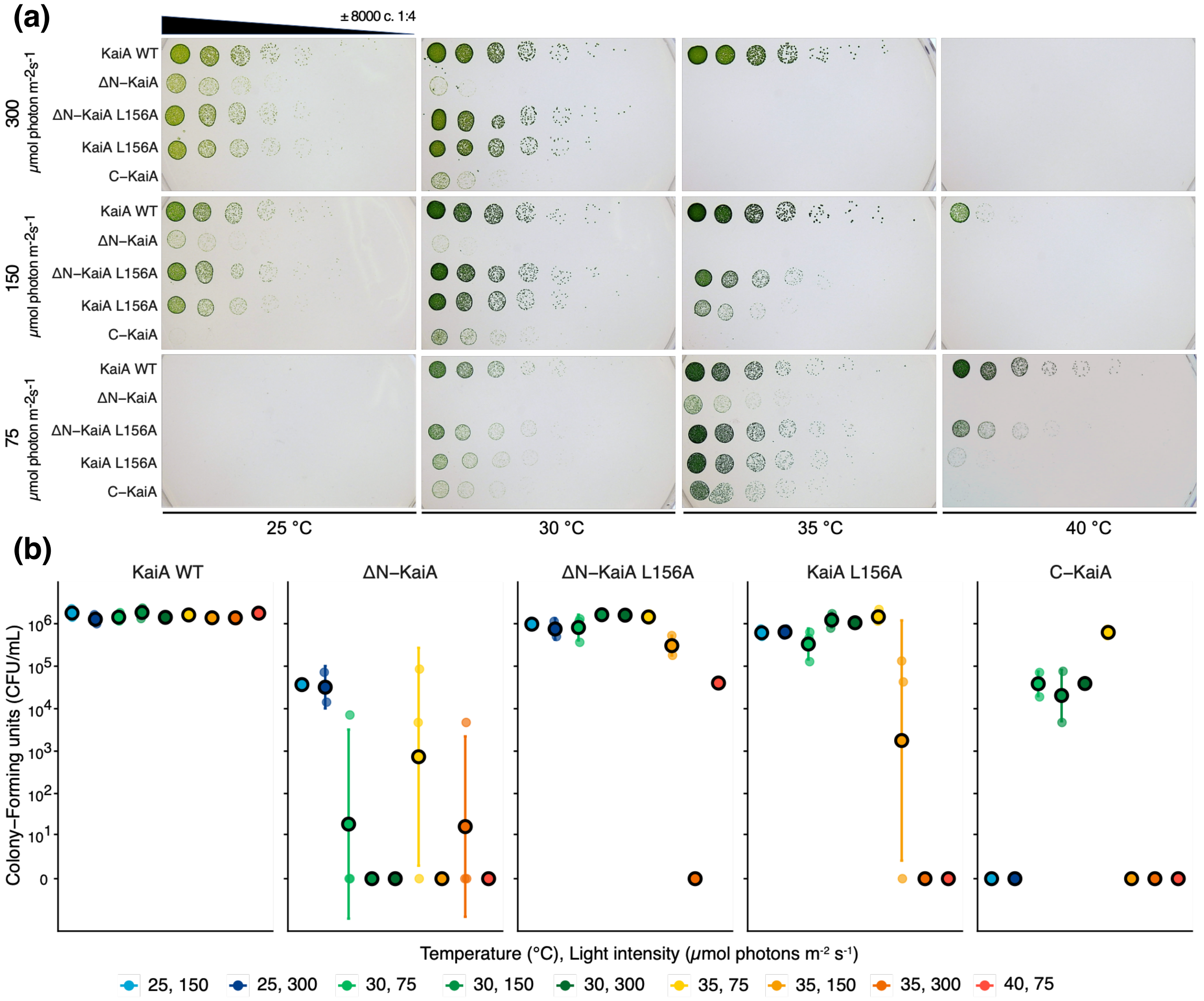

**Supplementary Figure 12. Growth (light–dark sensitivity) of KaiA WT and mutant strains under 12-h light:12-h dark cycles at different temperatures and light intensities.**

**(a)** Representative set of serial dilution spot plates incubated at 25, 30, 35, or 40 °C (left to right) under light intensities of 300, 150, or 75 $\mu$mol photons m$^{-2}$ s$^{-1}$ (top to bottom). Each row of a plate corresponds to a 4-fold serial dilution of Δ*kaiA* strains ectopically expressing KaiA WT, ΔN-KaiA, ΔN-KaiA L156A, KaiA L156A, or C-KaiA. Within each row, the number of cells spotted decreases from left to right, allowing the formation of isolated colonies that can be counted after incubation. Growth density of each spot and colony number across the dilution series shows the ability of each strain to form colonies under the indicated condition. Reduced colony number or loss of visible growth relative to KaiA WT, particularly at lower cell densities, indicates impaired growth under that condition. **(b)** Number of colonies in CFU/mL calculated from colony counts, dilution factors, and the 4-$\mu$L spotted volume on the dilution spot plates. Filled circles are biological-replicate means of spots with countable colonies; filled circles with a black outline and error bars show mean ± SD (n=3) calculated on a ln(1 + count) scale.

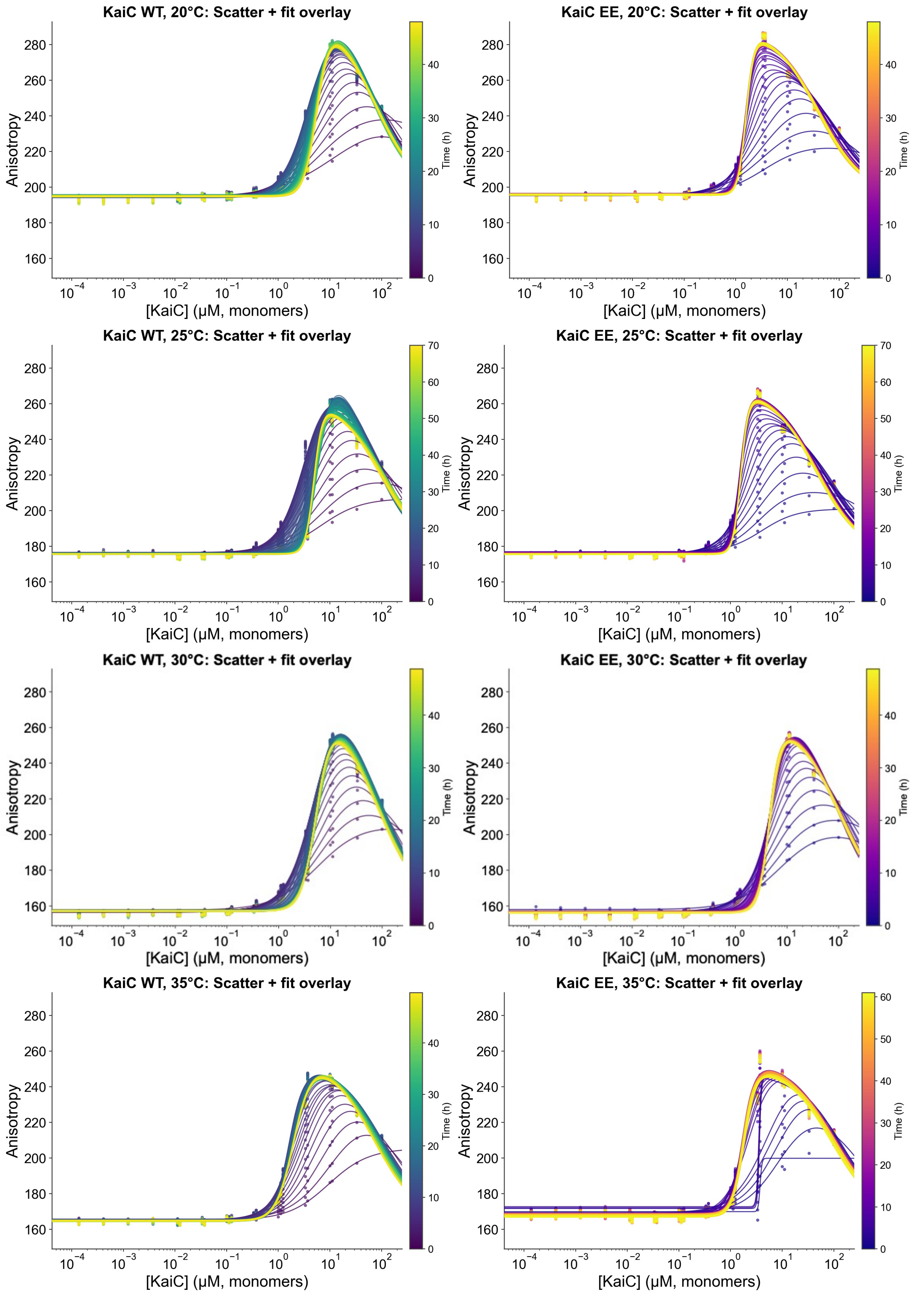
KaiC WT, 20°C: Scatter + fit overlay
KaiC EE, 20°C: Scatter + fit overlay
KaiC WT, 25°C: Scatter + fit overlay
KaiC EE, 25°C: Scatter + fit overlay
KaiC WT, 30°C: Scatter + fit overlay
KaiC EE, 30°C: Scatter + fit overlay
KaiC WT, 35°C: Scatter + fit overlay
KaiC EE, 35°C: Scatter + fit overlay
Anisotropy
[KaiC] (μM, monomers)
Time (h)

**Supplementary Figure 13. KaiB-KaiC equilibrium binding assay with fluorescence anisotropy.**

Time-resolved evolution of the KaiB-KaiC binding isotherm across temperature. Each panel corresponds to equilibrium binding assays of fluorescently labeled KaiB (50 nM), unlabeled KaiB WT (3.45 $\mu$M) with KaiC-EE (left) and KaiC WT (right). Anisotropy versus [KaiC] ($\mu$M, monomer, log scale) is shown for every 30-min time point, colored by elapsed time (color bar, h), with corresponding Hill fits overlaid as solid lines. The dotted gray line marks mean anisotropy at [KaiC] = 0. Progressive changes in curve position and shape reflect time-dependent shifts in apparent affinity (Kd1) and cooperativity (nH).

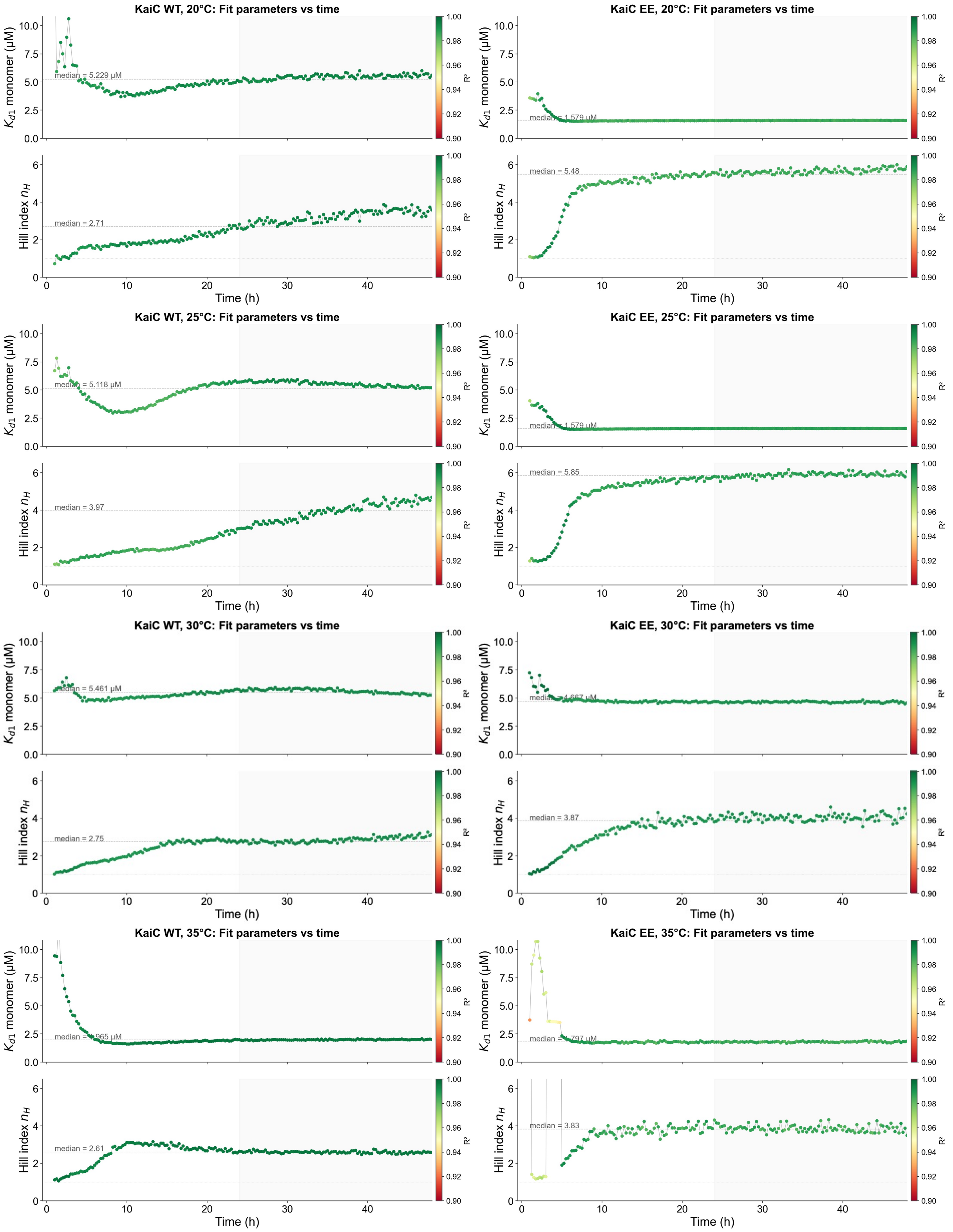
KaiC WT, 20°C: Fit parameters vs time
KaiC EE, 20°C: Fit parameters vs time
KaiC WT, 25°C: Fit parameters vs time
KaiC EE, 25°C: Fit parameters vs time
KaiC WT, 30°C: Fit parameters vs time
KaiC EE, 30°C: Fit parameters vs time
KaiC WT, 35°C: Fit parameters vs time
KaiC EE, 35°C: Fit parameters vs time
$K_{d1}$ monomer (μM)
Hill index $n_H$
Time (h)
$R^2$
median = 5.229 μM
median = 2.71
median = 5.48
median = 5.118 μM
median = 3.97
median = 5.85
median = 5.461 μM
median = 2.75
median = 3.87
median = 2.61
median = 3.83

**Supplementary Figure 14. Time course of fitted binding parameters for KaiB - KaiC [WT/EE] equilibrium binding assays.**

(From Supplementary Figure 13). Each panel corresponds to parameters from independent Hill fits at each time point for KaiC-EE (left) and KaiC WT (right). Top: Kd1 ($\mu$M, monomer). Middle: Hill cooperativity index n_H (dotted line, n_H = 1, marks the non-cooperative threshold). Bottom: $R^2$ of each fit (dashed red line, R¬≤ = 0.95). Points are colored by $R^2$ (color bar); the gray line connects successive time points, the dashed horizontal line marks the median, and gray shading indicates alternating 24-hour windows.

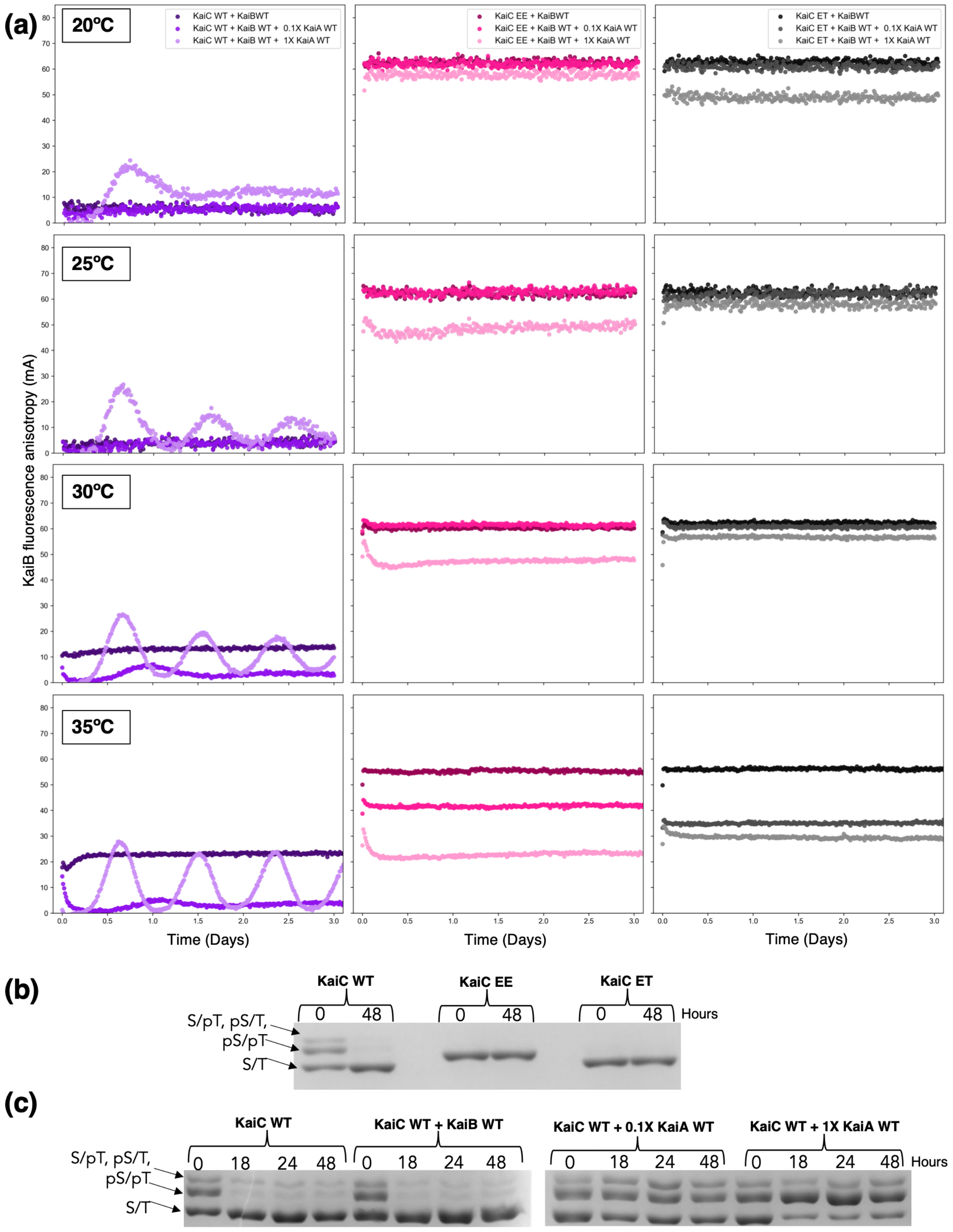

(a)
20ºC
25ºC
30ºC
35ºC
KaiC WT + KaiBWT
KaiC WT + KaiB WT + 0.1X KaiA WT
KaiC WT + KaiB WT + 1X KaiA WT
KaiC EE + KaiBWT
KaiC EE + KaiB WT + 0.1X KaiA WT
KaiC EE + KaiB WT + 1X KaiA WT
KaiC ET + KaiBWT
KaiC ET + KaiB WT + 0.1X KaiA WT
KaiC ET + KaiB WT + 1X KaiA WT
KaiB fluorescence anisotropy (mA)
Time (Days)
(b)
KaiC WT
KaiC EE
KaiC ET
0 48
Hours
S/pT, pS/T,
pS/pT
S/T
(c)
KaiC WT
KaiC WT + KaiB WT
KaiC WT + 0.1X KaiA WT
KaiC WT + 1X KaiA WT
0 18 24 48
Hours
S/pT, pS/T,
pS/pT
S/T

**Supplementary Figure 15. Fluorescence anisotropy assays and phosphorylation assays of KaiC mutants with KaiA.**

**(a)** Anisotropy assays with pre-incubated KaiB – KaiC mutants. Fluorescence anisotropy profiles of KaiB WT with dpKaiC WT, KaiC EE (S431E T432E), and KaiC ET (S431E). 0.1X or 1X KaiA WT was mixed with a 48-hour pre-incubated KaiB-KaiC mix. All rows contain KaiB WT (3.45 µM) and fluorescently labeled KaiB (50 nM) as the probe, and the columns increase in temperature from top to bottom: 20°C, 25°C, 30°C, and 35°C. **(b)** Phos-tag gel analysis of KaiC mutants (3.5 µM) phosphorylation states at time zero and after 48 hours (before mixing with 0.1X or 1X KaiA WT) at 35°C. **(c)** Phos-tag gel analysis of KaiC WT phosphorylation states over time (samples were not pre- dephosphorylated) at 35°C: (left to right), KaiC WT alone, KaiC WT and KaiB WT, KaiC WT, KaiB WT and 1/10$^{th}$ of KaiA WT(0.12 µM), and KaiB WT and KaiA WT(1.2 µM).

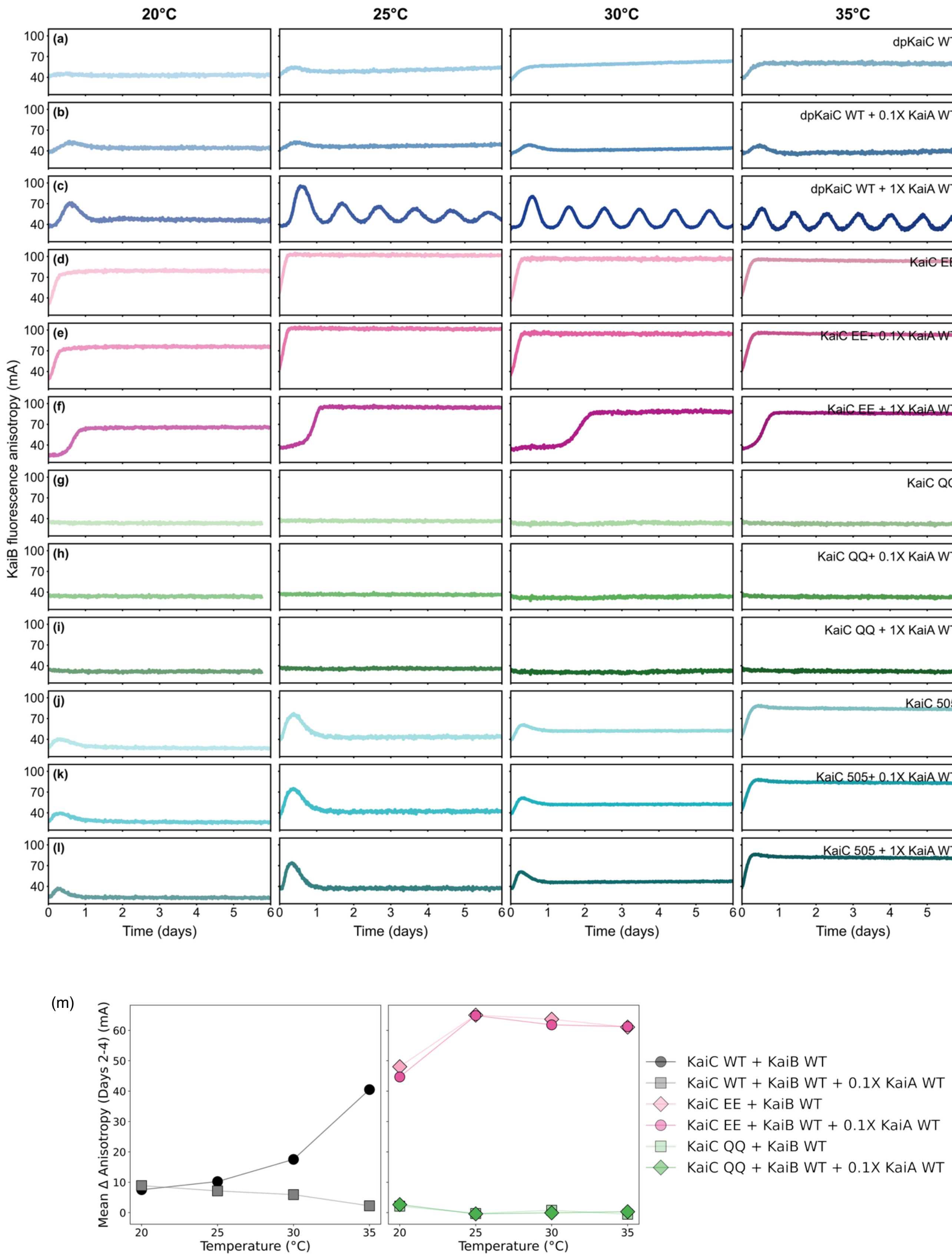
20°C
25°C
30°C
35°C
(a) dpKaiC WT
(b) dpKaiC WT + 0.1X KaiA WT
(c) dpKaiC WT + 1X KaiA WT
(d) KaiC EE
(e) KaiC EE+ 0.1X KaiA WT
(f) KaiC EE + 1X KaiA WT
(g) KaiC QQ
(h) KaiC QQ+ 0.1X KaiA WT
(i) KaiC QQ + 1X KaiA WT
(j) KaiC 505
(k) KaiC 505+ 0.1X KaiA WT
(l) KaiC 505 + 1X KaiA WT
KaiB fluorescence anisotropy (mA)
Time (days)
(m)
Mean Δ Anisotropy (Days 2-4) (mA)
Temperature (°C)
KaiC WT + KaiB WT
KaiC WT + KaiB WT + 0.1X KaiA WT
KaiC EE + KaiB WT
KaiC EE + KaiB WT + 0.1X KaiA WT
KaiC QQ + KaiB WT
KaiC QQ + KaiB WT + 0.1X KaiA WT

**Supplementary Figure 16. In vitro reconstituted fluorescence anisotropy assays with KaiC mutants (3.5 µM) across temperatures.**

Note that all the rows have KaiB WT (3.45 µM) and fluorescently labeled KaiB (50 nM) as the probe, and the columns from left to right increase in temperature: 20°C, 25°C, 30°C, and 35°C. **(a-c)** The initial three rows depict the dephosphorylated KaiC WT (blue) at 30°C overnight, which was subsequently mixed with KaiB WT (3.45 µM), fluorescently labeled KaiB (50 nM), and KaiA WT (1.2 µM (1X) or 0.12 µM (0.1X)). **(d-f)** The subsequent three rows show the double phosphomimic KaiC S431E T432E (pink), demonstrating enhanced KaiB-KaiC binding across various temperatures. Notably, higher concentrations of KaiA WT exhibit a more pronounced antagonistic effect on the initial KaiB-KaiC EE binding. **(g-i)** The KaiC CI ATPase inactive mutant, KaiC E77Q E78Q, exhibits no apparent increase in KaiB anisotropy even in the presence or absence of KaiA WT. **(j-l)** The final three rows present the KaiC 1-505 mutant, which lacks the C-terminal tail and is therefore incapable of interacting with KaiA. Interestingly, KaiC 505 also demonstrates temperature-dependent KaiB assembly, similar to dpKaiC WT, but fails to exhibit KaiA-dependent KaiB disassembly. **(m)** The mean change in Δ anisotropy was averaged over days 2-4 and plotted against temperature. KaiC WT anisotropy shows a strong temperature dependence compared to KaiC EE. Points show the mean of these replicate averages, and error bars show the standard deviation (n = 3). Lines connecting the points are a visual guide only to see the trend with temperature.

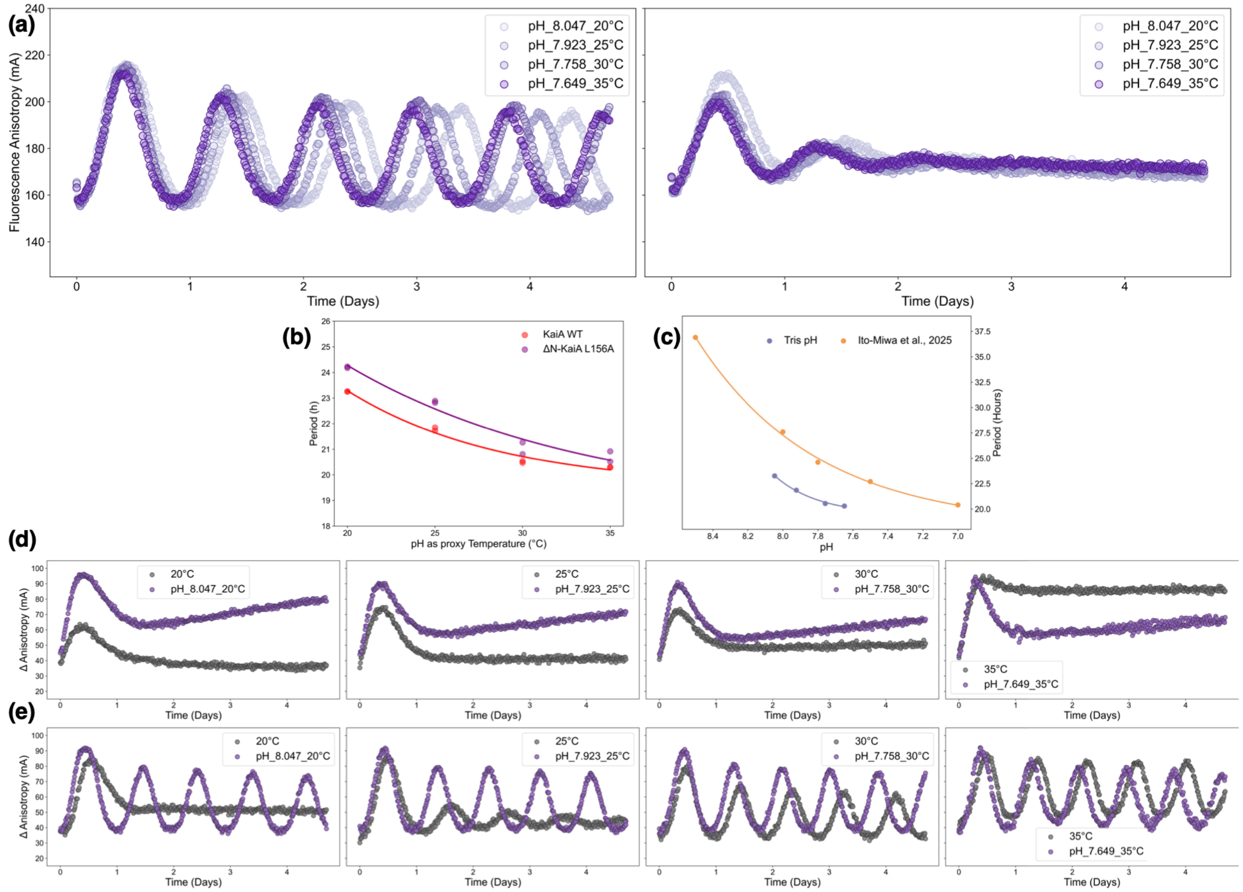

**Supplementary Figure 17. Effect of pH on reconstituted in vitro fluorescence anisotropy oscillations with Tris buffers.**

**(a)** (Left) KaiA WT and (Right) ΔN-KaiA L156A in vitro oscillations measured at 30°C with different pH Tris buffers spanning the temperature range 20°C, 25°C, 30°C, and 35°C. Oscillations are robust across the pH range, but the period is significantly affected by pH. **(b)** Period of the in vitro oscillation with KaiA WT and ΔN-KaiA L156A plotted against pH as a proxy for temperature. **(c)** Period of the in vitro oscillation plotted against buffer solution pH. Purple circles indicate data from this paper, and range circles indicate the pH dependence of the period of in vitro KaiC phosphorylation from Ito-Miwa et al. (2025) [26]. **(d)** KaiB-KaiC Disassembly is temperature-dependent with little pH-sensitivity. **(e)** The KaiABC IVO rhythms are predominantly temperature-dependent, and pH only influences the period.

(a)

(b)

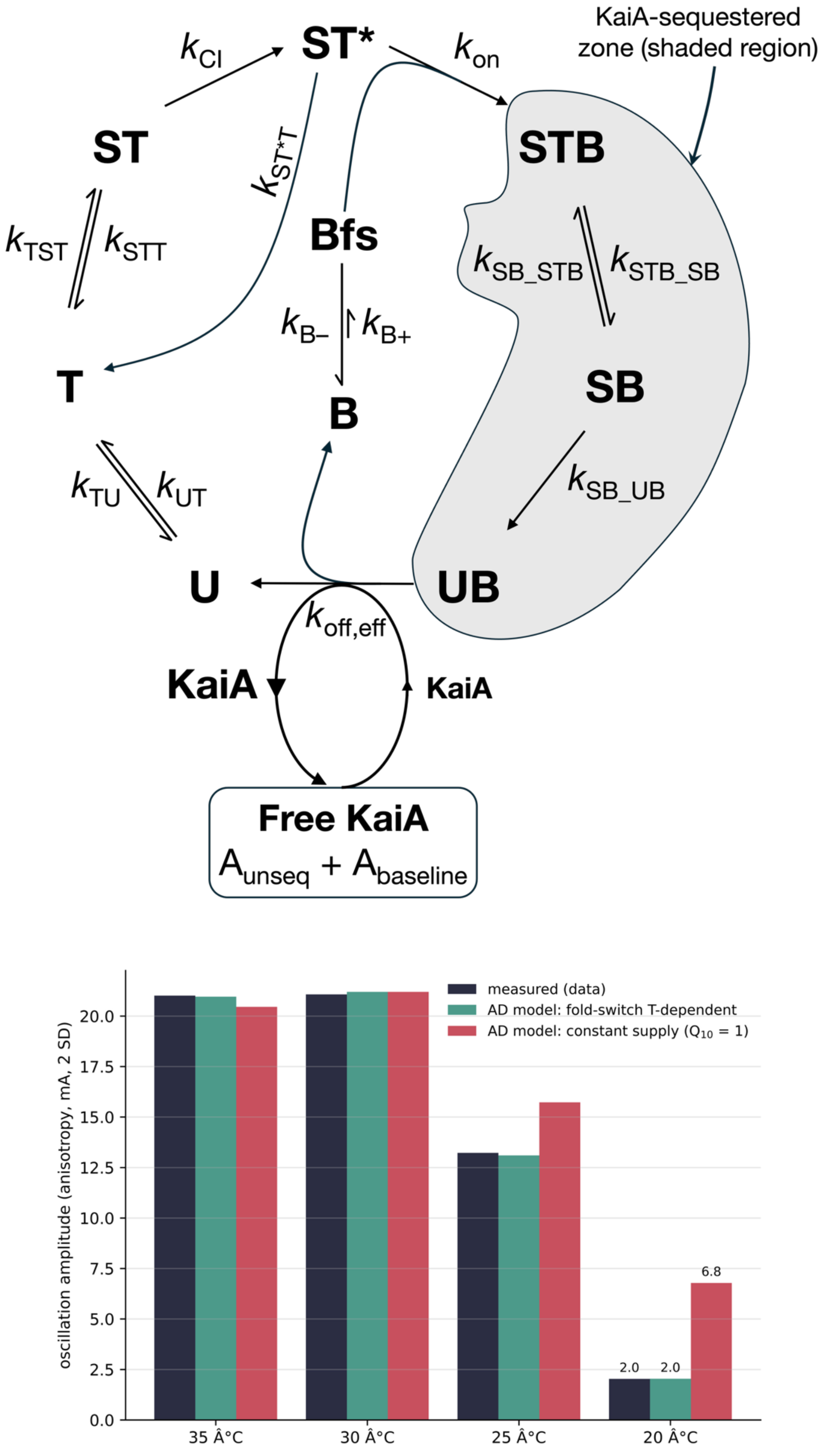

KaiA-sequestered zone (shaded region)
ST*
ST
STB
Bfs
T
B
SB
U
UB
$k_{CI}$
$k_{on}$
$k_{ST*T}$
$k_{TST}$
$k_{STT}$
$k_{SB\_STB}$
$k_{STB\_SB}$
$k_{B-}$
$k_{B+}$
$k_{TU}$
$k_{UT}$
$k_{SB\_UB}$
$k_{off,eff}$
KaiA
KaiA
Free KaiA
$A_{unseq} + A_{baseline}$
measured (data)
AD model: fold-switch T-dependent
AD model: constant supply ($Q_{10}$ = 1)
oscillation amplitude (anisotropy, mA, 2 SD)
0.0
2.5
5.0
7.5
10.0
12.5
15.0
17.5
20.0
6.8
2.0
2.0
35 Â°C
30 Â°C
25 Â°C
20 Â°C

**Supplementary Figure 18. Schematic representation of the Autocatalytic Dawn (AD) model.**

**(a)** STB, SB, and UB denote KaiA·KaiB·KaiC complexes; each sequesters one KaiA dimer per KaiC hexamer. KaiA is released as these complexes dissociate, replenishing the free-KaiA pool that drives the dawn cascade. STB, SB, and UB each sequester one KaiA dimer per KaiC hexamer biologically; the model represents this with an effective coupling A_unseq = KaiA_total – m·(STB+SB+UB), m = 3 (phenomenological, not literal stoichiometry). **(b)** The fold-switch temperature dependence is required for the model to reproduce the loss of rhythm at 20 °C

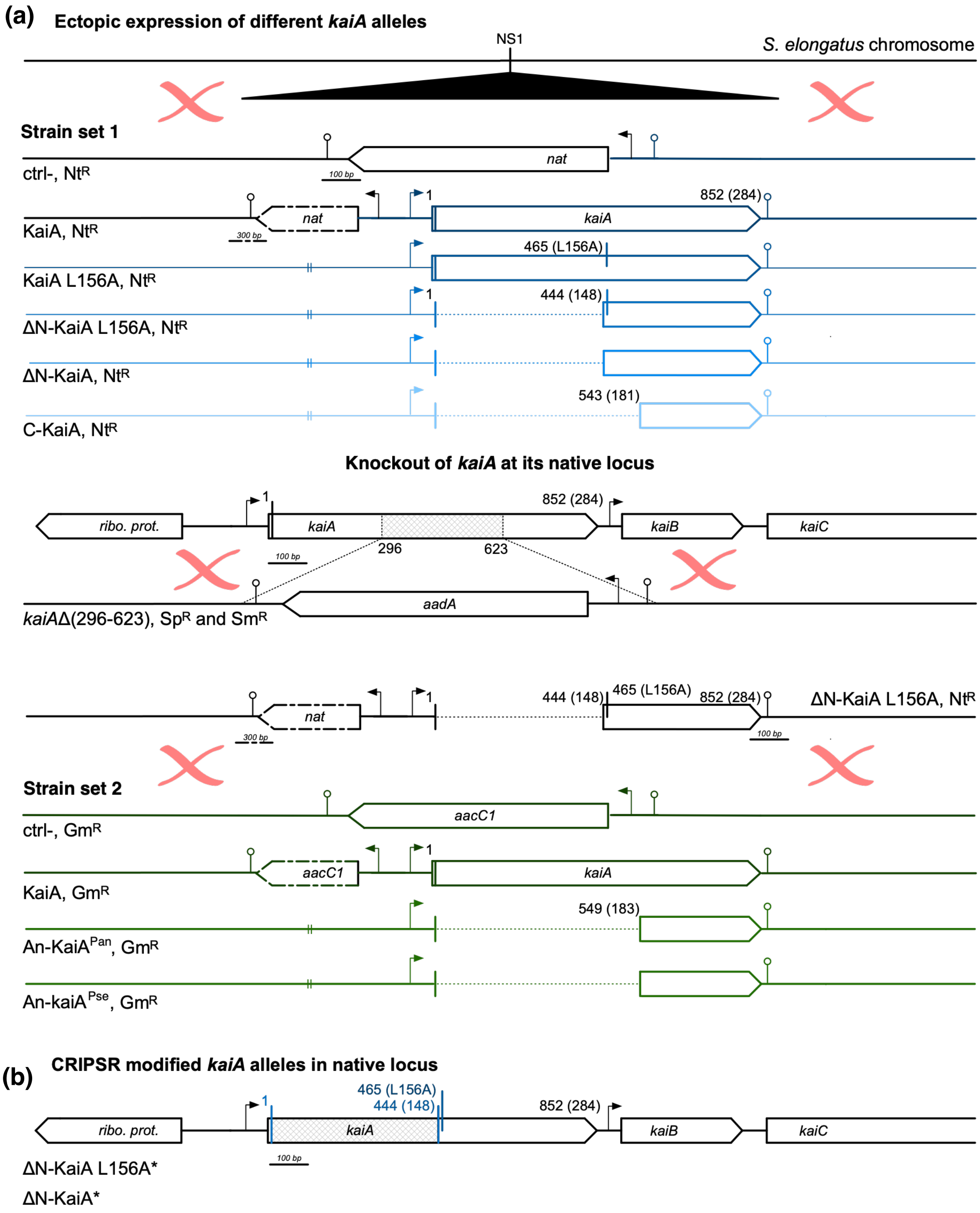

(a)
Ectopic expression of different kaiA alleles
NS1
S. elongatus chromosome
Strain set 1
nat
100 bp
ctrl-, NtR
1
852 (284)
nat
kaiA
300 bp
KaiA, NtR
465 (L156A)
KaiA L156A, NtR
1
444 (148)
ΔN-KaiA L156A, NtR
ΔN-KaiA, NtR
543 (181)
C-KaiA, NtR
Knockout of kaiA at its native locus
1
852 (284)
ribo. prot.
kaiA
kaiB
kaiC
296
623
100 bp
aadA
kaiAΔ(296-623), SpR and SmR
1
444 (148)
465 (L156A)
852 (284)
ΔN-KaiA L156A, NtR
nat
300 bp
100 bp
Strain set 2
aacC1
ctrl-, GmR
1
aacC1
kaiA
KaiA, GmR
549 (183)
An-KaiAPan, GmR
An-kaiAPse, GmR
(b)
CRIPSR modified kaiA alleles in native locus
1
465 (L156A)
444 (148)
852 (284)
ribo. prot.
kaiA
kaiB
kaiC
100 bp
ΔN-KaiA L156A*
ΔN-KaiA*

**Supplementary Figure 19. Schematic representation of genetic modifications introduced into the S. elongatus chromosome for the construction of the KaiA mutant strains used in this study.**

**(a)** The antibiotic resistance gene only or P*kaiA-kaiA* WT, P*kaiA-kaiA*(aa L156A), P*kaiA-kaiA*(aa 148-284, L156A), P*kaiA-kaiA*(aa 148-284), and P*kaiA-kaiA*(aa 181-284) alleles along with a Nt resistance gene (*nat*) were inserted at the NS1 of the *S. elongatus* chromosomes. The native *kaiA* locus was knocked out after insertion of the *kaiA* alleles at NS1. The *kaiA* coding sequence from *Anabaena* PCC 7120 was cloned either with its native promoter (An-$kaiA^{Pan}$) or placed downstream of the *S. elongatus kaiA* promoter (An-$kaiA^{Pse}$) and inserted along with a Gm resistance gene (*aacC1*) at NS1. Strain construction was facilitated by using the previously validated Δ*kaiA* expressing ΔN-KaiA L156A strain as an intermediate. **(b)** For ΔN-KaiA* and ΔN-KaiA L156A* strains, the *kaiA* alleles were edited by CRISPR at the native *kaiA* locus. A Δ*ago* strain of *S. elongatus* was used for CRISPR editing. Coordinates are given relative to the *kaiA* locus; corresponding amino-acid residues are indicated in between parentheses.

### Supplementary Tables

##### Supplementary Table 1. Cyanobacterial strains used in this study

| AMC# | Background strain | Plasmid(s) | Genotype | Resistance | Ref. |
|---|---|---|---|---|---|
| 2747 | AMC2665 | pAM2105 | NS2::(*cat*-P*kaiB*-*luc*) | Cm | [22] |
| 2749 | AMC2752 | pAMT010, pAM2105 | *ago*Δ(-117..2075), NS2::(*cat*-P*kaiB*-*luc*) | Cm | [22] |
| 2766 | AMC2747 | pAM5903, pAM5901 | NS2::(*cat*-$P_{kaiB}$-*luc*); NS1::(nat7942-$P_{kaiA}$-*kaiA*(aa 148-284)); *kaiA*Δ(296-623)::*aadA* | Cm, Nt, Sp + Sm | This study |
| 2767 | AMC2747 | pAM5909, pAM5902 | NS2::(*cat*-$P_{kaiB}$-*luc*); NS1::(nat7942-$P_{kaiA}$-*kaiA*(aa 148-284, L156A)); *kaiA*Δ(296-623)::*aadA* | Cm, Nt, Sp + Sm | This study |
| 2768 | AMC2747 | pAM5910, pAM5903 | NS2::(*cat*-$P_{kaiB}$-*luc*); NS1::(nat7942-$P_{kaiA}$-*kaiA*(aa 181-284)); *kaiA*Δ(296-623)::*aadA* | Cm, Nt, Sp + Sm | This study |
| 2769 | AMC2747 | pAM5904, pAM5900 | NS2::(*cat*-$P_{kaiB}$-*luc*); NS1::(nat7942-$P_{kaiA}$- *kaiA(*aa L156A)); *kaiA*Δ(296-623)::*aadA* | Cm, Nt, Sp + Sm | This study |

| | | | | | |
|---|---|---|---|---|---|
| 2770 | AMC2747 | pAM5907, pAM5900 | NS2::(*cat*-$P_{kaiB}$-*luc*); NS1::(nat7942-$P_{kaiA}$-*kaiA*$^{+}$); *kaiA*Δ(296-623)::*aadA* | Cm, Nt, Sp + Sm | This study |
| 2785 | AMC2767 | pAM5946 | NS2::(*cat*-$P_{kaiB}$-*luc*); NS1::aacC1-P*kaiA*Se-*kaiA*An; *kaiA*Δ(296-623)::*aadA* | Sp + Sm, Cm, Gm | This study |
| 2786 | AMC2767 | pAM5944 | NS2::(*cat*-$P_{kaiB}$-*luc*); NS1::aacC1-P*kaiA*$_{An}$-*kaiA*$_{An}$; *kaiA*Δ(296-623)::*aadA* | Sp + Sm, Cm, Gm | This study |
| 2787 | AMC2767 | pAM5945 | NS2::(*cat*-$P_{kaiB}$-*luc*); NS1::aacC1-P*kaiA*Se-*kaiA*Se; *kaiA*Δ(296-623)::*aadA* | Sp + Sm, Cm, Gm | This study |
| 2788 | AMC2767 | pAM5943 | NS2::(*cat*-$P_{kaiB}$-*luc*); NS1::aacC1; *kaiA*Δ(296-623)::*aadA* | Sp + Sm, Cm, Gm | This study |
| 2802 | AMC2747 | pAM5915, pAM5904 | NS2::(*cat*-$P_{kaiB}$-*luc*); NS1::(nat7942-$P_{kaiA}$-*kaiA*$^{[1,164-*]}$); *kaiA*Δ(296-623)::*aadA* | Cm, Nt, Sp + Sm | This study |
| 2803 | AMC2747 | pAM5613, pAM5905 | NS2::(*cat*-$P_{kaiB}$-*luc*); NS1::nat7942; *kaiA*Δ(296-623)::*aadA* | Cm, Nt, Sp + Sm | This study |

| | | | | | |
|---|---|---|---|---|---|
| 2804 | AMC2749 | pAM5908, pAM5819 | agoΔ(-117..2075), NS2::PkaiB-luc, *kaiA*(aa 148-284) | Cm | This study |
| 2805 | AMC2749 | pAM5909, pAM5819 | agoΔ(-117..2075), NS2::PkaiB-luc; *kaiA*(aa 148-284, L156A) | Cm | This study |

**Supplementary Table 2. Plasmid used in this study**

| **AM#** | **Backbone/ source(s)** | **Plasmid name** | **Plasmid genotype** | **Resistance** | **Ref.** |
|---|---|---|---|---|---|
| 4819 | | pCVD003 | APHI | Ap, Km | [24] |
| 4831 | pMA-RQ | pCVD015 | ccdB/SwaI | Ap | [24] |
| 4835 | pUC19 | pCVD019 | NS1RK2BOM | Ap | [24] |
| 4836 | pUC19 | pCVD020 | NS1TC | Ap, Tc | |
| 4880 | pMA-RQ | pCVD064 | NAT7942 | Ap, Nt | |
| 4940 | | pCV0052 | S7942NS1_AACC1_CCDB | | [24] |

| | | | | | |
|---|---|---|---|---|---|
| 5001 | | pAM5001 | *aadA-sacB* | SpSm | Lab collection (R. Simkovsky) |
| 5293 | | pAM5293 | Template for L156A | | Lab collection (S. Cohen) |
| 5406 | | pAM5406 | RSF1010BB | Ap | [27] |
| 5600 | pCVD015 | pAM5600 | CRISPR ARRAY-cpf1 | Ap | [22] |
| 5604 | pCVD019, pCVD015, pCVD064 | pAM5604 | NS1RK2B-NAT7942-*ccdB*/SwaI | Nt | This study |
| 5613 | pAM5604 | pAM5613 | NS1RK2B-NAT7942 | Nt | This study |
| 5819 | pAMT004 | pAM5819 | pAMT004-gRNA(*kaiA_112*) | Km | This study |
| 5820 | pAMT004 | pAM5820 | pAMT004-gRNA(*kaiA_156*) | | |

| | | | | | |
|---|---|---|---|---|---|
| 5831 | pUC19 | pAM5831 | pUC19-..*kaiA*Δ(296-623)::(*aadA-sacB*).. | Ap, Sp + Sm | This study |
| 5832 | pCVD015 | pAM5832 | pCVD015-$P_{kaiA}$-*kaiA*(aa 148-284) | Ap | This study |
| 5833 | pCVD015 | pAM5833 | pCVD015-$P_{kaiA}$-*kaiA*(aa 148-284, L156A) | Ap | This study |
| 5834 | pCVD015 | pAM5834 | pCVD015-$P_{kaiA}$-*kaiA*(aa 181-284) | Ap | This study |
| 5847 | pCVD015 | pAM5847 | pCVD015-$P_{kaiA}$-*kaiA* | Ap | This study |
| 5848 | pCVD015 | pAM5848 | pCVD015-$P_{kaiA}$-*kaiA(aa L156A)* | Ap | This study |
| 5900 | pAM5831 | pAM5900 | pUC19-..*kaiA*Δ(296-623)::*aadA..* | Ap, Sp + Sm | This study |
| 5903 | pCVD020, pCVD064, pAM5832 | pAM5903 | NS1TC-NAT7942-$P_{kaiA}$-*kaiA*(aa 148-284) | Tc*, Nt | This study |

| | | | | | |
|---|---|---|---|---|---|
| 5904 | pCVD020, pCVD064, pAM5833 | pAM5904 | NS1TC-NAT7942-$P_{kaiA}$-*kaiA*(aa 148-284, L156A) | Tc*, Nt | This study |
| 5905 | pCVD020, pCVD064, pAM5834 | pAM5905 | NS1TC-NAT7942-$P_{kaiA}$- *kaiA*(aa 181-284) | Tc*, Nt | This study |
| 5906 | pCVD020, pCVD064, pAM5848 | pAM5906 | NS1TC-NAT7942-$P_{kaiA}$-*kaiA*(aa L156A) | Tc*, Nt | This study |
| 5907 | pCVD020, pCVD064, | pAM5907 | NS1TC-NAT7942-$P_{kaiA}$-*kaiA*$^{+}$ | Tc*, Nt | This study |

| | | | | | |
|---|---|---|---|---|---|
| | pAM5847 | | | | |
| 5908 | pAM5406, pCVD064, pAM5832 | pAM5908 | RSF1010BB-NAT7942-..P*kaiA*-*kaiA*(aa 148-284).. | Nt | This study |
| 5909 | pAM5406, pCVD064, pAM5833 | pAM5909 | RSF1010BB-NAT7942-..P*kaiA*-*kaiA*(aa 148-284, L156A).. | Nt | This study |
| 5943 | pCV0052 | pAM5943 | S7942NS1_AACC1 | Gm | This study |
| 5944 | pCV0052 | pAM5944 | S7942NS1_AACC1_P*kaiA*An-*kaiA*An) | Gm | This study |
| 5945 | pCV0052 | pAM5945 | S7942NS1_AACC1_P*kaiA*Se-*kaiA*Se) | Gm | This study |
| 5946 | pCV0052 | pAM5946 | S7942NS1_AACC1_P*kaiA*Se-*kaiA*An) | Gm | This study |

| | | | | | |
|---|---|---|---|---|---|
| 5997 | pAM5406, pCVD003, pAMT001 | pAMT004 | RSF1010BB-APH1-CPF1-CRISPR | Km | This study |

**Supplementary Table 3. Primers used in this study**

| name | ATP | sequence 5'-3' |
|---|---|---|
| 112_Fwd_SA | | agatAGCCAGTGCAGCATCAACTT |
| 112_Rev_SA | | agacAAGTTGATGCTGCACTGGCT |
| 156_1_Fwd_SA | | agatGCTCAGCGACTACAAGAGCG |
| 156_1_Rev_SA | | agacCGCTCTTGTAGTCGCTGAGC |
| aadA-pCVD2_R | | GATATCGCGCGCGCGCGCGGATGCTAGCGGCGCGCCAAAAAAAA |
| aadA-sacB_F | [ATP0121] | GATATCCCCCCGGGGGCCCCCGATTCTAGAAAGCTTCAAAAAGGCCA |
| kaiA_148F | [ATP1491] | CGACTGCGTGGAGCTATCGAGCCAGCAGC |
| kaiA_181F | [ATP1492] | CGACTGCGTGAACCTACCCGCCTACGAAAG |

| | | |
|---|---|---|
| kaiA_296R | [ATP1465] | GCGCGCGCGCGCGATATCTAGAGCTGTTCTTTGGCTGGTTC |
| kaiA_623F | [ATP1466] | GGGGCCCCGGGGGGATATCCGAATAGCAACCTCAACCAGAGC |
| kaiA_AA1R_1 | [ATP1488] | TCGATAGCTCCACGCAGTCGCTCCTGTC |
| kaiA_AA1R_2 | [ATP1489] | CGGGTAGGTTCACGCAGTCGCTCCTGTC |
| kaiA_D1763R | [ATP1467] | CGACGGCCAGTGAATTCGAGTATTGCTGGAAAACGGACGTGAC |
| kaiA_Down_R | [ATP1490] | GCCGGGGAGCTCCTTCATTTCAACGGGGTAGCAACAGCAATC |
| kaiA_U607F | [ATP1464] | GAGGATCCCCGGGTACCGAGCAATGTCCGCTTCTGATTTCGGT |
| kaiAa_3F/kaiAa_F | | aggagcgactgcgtgACTCAAGAAGTTGATCAGCAAATAT |
| kaiAa_D433R | | gccggggagctccttcatttTCTTTCTGGCTTTATTCATAAAGGT |
| pkaiA_F/pKaiAse_F | [ATP1487] | GGCCAATAACCCAGGGATTTTCCAAGTCTGCGTCGAGGC |
| pKaiAana_F | | ataacccagggatttGTTTAAGTATCTTATCAGGGAGCAG |
| pKaiAse_R | | CAACTTCTTGAGTcacgcagtcgctcctgtcagga |

**Supplementary Table 4. Constructs and shorthand names used in the in vitro experiments.**

| Shorthand name | Protein constructs full name |
|---|---|
| KaiA WT | *S.e.* KaiA 1 - 284 |
| KaiA L156A | *S.e.* KaiA 1 - 284 L156A |
| ΔN KaiA | *S.e.* KaiA 148 - 284 |
| ΔN KaiA L156A | *S.e.* KaiA 148 - 284 L156A |
| C-KaiA | *S.e.* KaiA 181 - 284 |
| KaiB WT | *S.e.* KaiB 1 - 102 FLAG |
| KaiB -K25C-6IAF | *S.e.* KaiB 1 - 102 K25C – FLAG - 6-iodoacetamido-fluorescein |
| fsKaiB | *S.e.* KaiB 1 - 99 - Y7A - I87A - Y93A - FLAG |
| KaiC WT | FLAG *S.e.* KaiC 1 - 519 |
| KaiC EE | FLAG *S.e.* KaiC 1 - 519 – S431E – T432E |
| KaiC ET | FLAG *S.e.* KaiC 1 - 519 – S431E |
| KaiC QQ | FLAG *S.e.* KaiC 1 - 519 – E77Q – E78Q |
| KaiC 505 | FLAG *S.e.* KaiC 1 - 505 |
| KaiC EE 505 | FLAG *S.e.* KaiC 1 – 505 - S431E – T432E |

**Supplementary Table 5. Protein expression and purification conditions**

| Protein construct | Expression temperature and duration | Affinity column Loading buffer | Affinity column Wash buffer | Affinity column Elution buffer | Gel Filtration Column and Elution Buffer |
|---|---|---|---|---|---|
| KaiA WT | 22 ºC, 18 h | 50 mM NaH2PO4, 500 mM NaCl, pH 8 | 50 mM NaH2PO4, 500 mM NaCl, 20 mM imidazole pH 8 | 50 mM NaH2PO4, 500 mM NaCl, 250 mM imidazole pH 8 | Superdex-200-16/600 20 mM Tris, 150 mM NaCl, 5 mM MgCl2, 1 mM ATP, 0.5 mM EDTA, pH 8 |
| KaiA L156A | 22 ºC, 18 h | 50 mM NaH2PO4, 500 mM NaCl, pH 8 | 50 mM NaH2PO4, 500 mM NaCl, 20 mM imidazole pH 8 | 50 mM NaH2PO4, 500 mM NaCl, 250 mM imidazole pH 8 | Superdex-200-16/600 20 mM Tris, 150 mM NaCl, 5 mM MgCl2, 1 mM ATP, 0.5 mM EDTA, pH 8 |
| ΔN KaiA | 22 ºC, 18 h | 50 mM NaH2PO4, | 50 mM NaH2PO4, 500 | 50 mM NaH2PO4, 500 | Superdex-75-16/600 |

| | | | | | |
|---|---|---|---|---|---|
| | | 500 mM NaCl, pH 8 | mM NaCl, 20 mM imidazole pH 8 | mM NaCl, 250 mM imidazole pH 8 | 20 mM Tris, 150 mM NaCl, 5 mM MgCl2, 1 mM ATP, 0.5 mM EDTA, pH 8 |
| ΔN KaiA L156A | 22 °C, 18 h | 50 mM NaH2PO4, 500 mM NaCl, pH 8 | 50 mM NaH2PO4, 500 mM NaCl, 20 mM imidazole pH 8 | 50 mM NaH2PO4, 500 mM NaCl, 250 mM imidazole pH 8 | Superdex-75-16/600 20 mM Tris, 150 mM NaCl, 5 mM MgCl2, 1 mM ATP, 0.5 mM EDTA, pH 8 |
| C-KaiA | 22 °C, 18 h | 50 mM NaH2PO4, 500 mM NaCl, pH 8 | 50 mM NaH2PO4, 500 mM NaCl, 20 mM imidazole pH 8 | 50 mM NaH2PO4, 500 mM NaCl, 250 mM imidazole pH 8 | Superdex-75-16/600 20 mM Tris, 150 mM NaCl, 5 mM MgCl2, 1 mM ATP, 0.5 mM EDTA, pH 8 |
| KaiB WT | 22 °C, 18 h | 50 mM NaH2PO4, | 50 mM NaH2PO4, 500 | 50 mM NaH2PO4, 500 | Superdex-75-16/600 |

| | | | | | |
|---|---|---|---|---|---|
| | | 500 mM NaCl, pH 8 | mM NaCl, 20 mM imidazole pH 8 | mM NaCl, 250 mM imidazole pH 8 | 20 mM Tris, 150 mM NaCl, 5 mM MgCl2, 1 mM ATP, 0.5 mM EDTA, pH 8 |
| fsKaiB | 22 °C, 18 h | 50 mM NaH2PO4, 500 mM NaCl, pH 8 | 50 mM NaH2PO4, 500 mM NaCl, 20 mM imidazole pH 8 | 50 mM NaH2PO4, 500 mM NaCl, 250 mM imidazole pH 8 | Superdex-75-16/600<br>20 mM Tris, 150 mM NaCl, 5 mM MgCl2, 1 mM ATP, 0.5 mM EDTA, pH 8 |
| KaiB K25C | 22 °C, 18 h | 50 mM NaH2PO4, 500 mM NaCl, pH 8 | 50 mM NaH2PO4, 500 mM NaCl, 20 mM imidazole pH 8 | 50 mM NaH2PO4, 500 mM NaCl, 250 mM imidazole pH 8 | Superdex-75-16/600<br>Labeling buffer: 20 mM Tris, 150 mM NaCl, pH 7<br><br>Final buffer: |

| | | | | | |
|---|---|---|---|---|---|
| | | | | | 20 mM Tris, 150 mM NaCl, 5 mM MgCl2, 1 mM ATP, 0.5 mM EDTA, pH 8 |
| KaiC WT | 22 °C, 18 h | 50mM Tris,150mM NaCl, 1mM ATP, 5mM MgCl2, 5% Glycerol, pH 8 | 50mM Tris,150mM NaCl, 1mM ATP, 5mM MgCl2, 5% Glycerol, pH 8 | 50mM Tris,150mM NaCl, 50 mM Biotin, 1mM ATP, 5mM MgCl2, 5% Glycerol, pH 8 | Superdex-200-16/600 20 mM Tris, 150 mM NaCl, 5 mM MgCl2, 1 mM ATP, 0.5 mM EDTA, pH 8 |
| KaiC QQ | 22 °C, 18 h | 50mM Tris,150mM NaCl, 1mM ATP, 5mM MgCl2, 5% Glycerol, pH 8 | 50mM Tris,150mM NaCl, 1mM ATP, 5mM MgCl2, 5% Glycerol, pH 8 | 50mM Tris,150mM NaCl, 50 mM Biotin, 1mM ATP, 5mM MgCl2, 5% Glycerol, pH 8 | Superdex-200-16/600 20 mM Tris, 150 mM NaCl, 5 mM MgCl2, 1 mM ATP, 0.5 mM EDTA, pH 8 |

| | | | | | |
|---|---|---|---|---|---|
| KaiC EE | 22 ºC, 18 h | 50mM Tris,150mM NaCl, 1mM ATP, 5mM MgCl2, 5% Glycerol, pH 8 | 50mM Tris,150mM NaCl, 1mM ATP, 5mM MgCl2, 20 mM Imidazole, 5% Glycerol, pH 8 | 50mM Tris,150mM NaCl, 1mM ATP, 5mM MgCl2, 250 mM Imidazole, 5% Glycerol, pH 8 | Superdex-200-16/600 20 mM Tris, 150 mM NaCl, 5 mM MgCl2, 1 mM ATP, 0.5 mM EDTA, pH 8 |
| KaiC 505 | 22 ºC, 18 h | 50mM Tris,150mM NaCl, 1mM ATP, 5mM MgCl2, 5% Glycerol, pH 8 | 50mM Tris,150mM NaCl, 1mM ATP, 5mM MgCl2, 5% Glycerol, pH 8 | 50mM Tris,150mM NaCl, 50 mM Biotin, 1mM ATP, 5mM MgCl2, 5% Glycerol, pH 8 | Superdex-200-16/600 20 mM Tris, 150 mM NaCl, 5 mM MgCl2, 1 mM ATP, 0.5 mM EDTA, pH 8 |
| KaiC EE 505 | 22 ºC, 18 h | 50mM Tris,150mM NaCl, 1mM ATP, 5mM | 50mM Tris,150mM NaCl, 1mM ATP, 5mM MgCl2, 20 mM | 50mM Tris,150mM NaCl, 1mM ATP, 5mM MgCl2, 250 | Superdex-200-16/600 20 mM Tris, 150 mM NaCl, 5 mM |

| | | | | | |
|---|---|---|---|---|---|
| | | MgCl2, 5% Glycerol, pH 8 | Imidazole, 5% Glycerol, pH 8 | mM Imidazole, 5% Glycerol, pH 8 | MgCl2, 1 mM ATP, 0.5 mM EDTA, pH 8 |
| KaiC ET | 22 ºC, 18 h | 50mM Tris,150mM NaCl, 1mM ATP, 5mM MgCl2, 5% Glycerol, pH 8 | 50mM Tris,150mM NaCl, 1mM ATP, 5mM MgCl2, 20 mM Imidazole, 5% Glycerol, pH 8 | 50mM Tris,150mM NaCl, 1mM ATP, 5mM MgCl2, 250 mM Imidazole, 5% Glycerol, pH 8 | Superdex-200-16/600 20 mM Tris, 150 mM NaCl, 5 mM MgCl2, 1 mM ATP, 0.5 mM EDTA, pH 8 |

**Supplementary Table 6. KaiA-dependent KaiC initial phosphorylation rates (per hour) across varying temperatures (Fig. 2C, Supplementary Fig. 8)**

| Temperature | 20°C | 25°C | 30°C | 35°C |
|---|---|---|---|---|
| KaiA WT | 0.11 | 0.12 | 0.16 | 0.17 |
| KaiA L156A | 0.09 | 0.14 | 0.14 | 0.18 |
| ΔN KaiA | 0.02 | 0.02 | -0.01 | 0 |
| ΔN KaiA L156A | 0.11 | 0.13 | 0.18 | 0.16 |
| C-KaiA | 0.12 | 0.12 | 0.17 | 0.17 |

**Supplementary Table 7. Dissociation constants (Kd) in *μ*M calculated in KaiC A-loop anisotropy experiments (Fig. 2D, Supplementary Fig. 9)**

| Temperature | 20°C | 30°C | 40°C |
|---|---|---|---|
| KaiA WT | 261.83 ± 10.78 | 300.98 ± 12.81 | 382.8 ± 28.39 |
| KaiA L156A | 187.56 ± 10.99 | 188.51 ± 10.02 | 213.31 ± 12.34 |
| ΔN KaiA | 1024 ± 160.32 | 1024 ± 272.99 | 1024 ± 245.83 |
| ΔN KaiA L156A | 386.63 ± 28.46 | 494.2 ± 52.09 | 616.23 ± 62.95 |
| C-KaiA | 293.26 ± 36.89 | 328.44 ± 47.8 | 394.35 ± 46.29 |

**Supplementary Table 8. $^{19}$F NMR spectrum analysis of 7-fluoro-Trp labeled KaiA mutants**

| Mutant | Peak | Line width | Δ$\delta$ (ppm) | Δn (Hz) |
|---|---|---|---|---|
| **KaiA_135N** | -50.513 | 0.33171 | | |
| **KaiA_WT** | -50.561 | 0.4763 | 0.048 | 28.8 |
| **KaiA_L156A** | -50.608 | 0.51979 | 0.095 | 57 |
| **KaiA_WT_FsKaiB** | -50.511 | 0.58032 | -0.002 | -1.2 |

**Supplementary Table 9. Q10 calculated for KaiA mutants in temperature-dependent experiments**

| | Experiment | KaiA mutants | Q10 |
|---|---|---|---|
| 1. | In vitro oscillations (fluorescence anisotropy) | KaiA WT | 1.1 |
| | | ΔN KaiA L156A | 0.97 |
| 2. | In vivo oscillations (bioluminescence) | KaiA WT | 1.04 |
| | | ΔN KaiA L156A | 0.78 |
| 3. | KaiC phosphorylation assay (rate of phosphorylation) | KaiA WT | 1.38 |
| | | $KaiA_{L156A}$ | 1.45 |

| | | | |
|---|---|---|---|
| | | ΔN KaiA | - |
| | | ΔN KaiA L156A | 1.29 |
| | | C-KaiA | 1.34 |
| 4. | KaiC A loop anisotropy assay (dissociation constant) | KaiA WT | 1.21 |
| | | KaiA L156A | 1.06 |
| | | ΔN KaiA | - |
| | | ΔN KaiA L156A | 1.26 |
| | | C-KaiA | 1.16 |

**Supplementary Table 10. Kd1 (*μ*M, monomer) from KaiB-KaiC equilibrium binding assay anisotropy experiments**

| Kd1 (*μ*M, monomer) | 20°C | 25°C | 30°C | 35°C |
|---|---|---|---|---|
| KaiC WT | 5.60 ± 0.51 | 5.20 ± 0.76 | 5.25 ± 0.63 | 2.01 ± 0.08 |
| KaiC EE | 1.58 ± 0.25 | 1.58 ± 0.26 | 4.56 ± 0.04 | 1.85 ± 0.17 |

**Supplementary Table 11. Molecular weight calculated from SEC-MALS**

| Protein | Experimental MW (kDa) | Theoretical MW monomer (kDa) | |
|---|---|---|---|
| KaiA WT | 64.7 | 32.6 | KaiA Dimer |
| KaiA L156A | 65 | 32.56 | KaiA Dimer |
| ΔN KaiA | 31.91 | 16.25 | KaiA Dimer |
| ΔN KaiA L156A | 32.15 | 16.21 | KaiA Dimer |
| C-KaiA | 24.37 | 12.2 | KaiA Dimer |
| fsKaiB | 10.14 | 11.83 | KaiB Monomer |
| KaiC EE 505 | 340 | 57.54 | Hexamer |
| KaiC EE 505 + fsKaiB | 400 | | Hexamer KaiC + 6 KaiB monomer |
| KaiC EE 505 + fsKaiB + KaiA WT peak 1 | 513 | | Hexamer KaiC + 6 KaiB monomer + KaiA WT |
| KaiC EE 505 + fsKaiB + KaiA L156A peak 1 | 400 | | Hexamer KaiC + 6 KaiB monomer |
| KaiC EE 505 + fsKaiB + KaiA L156A peak 2 | 65 | | KaiA dimer |
